# A dynamic proteomic HECT E3 ligase screen reveals substrate exclusivity and redundancies

**DOI:** 10.64898/2026.11.24.768805

**Authors:** Bhavesh S. Parmar, Napyin Ho, Christina Schülein-Völk, Parviz Ghezellou, Nikita A. Verheyden, Ursula Eilers, Martin Eilers, Christian Münch

## Abstract

Cells maintain proteostasis through constant purge of proteins, complexes and large assemblies via the ubiquitin proteasome system (UPS). As a catalyst of ubiquitin transfer, E3 ligases determine the fate of their substrates, and to an extent, of the cell. Current approaches in identifying E3-substrate pairs largely overlook the dynamic nature of their interaction and the functional implications on cellular proteostasis. To address this, we incorporated a pSILAC-enabled dynamic proteomics into a knockdown (KD) screen for the HECT E3 ligase family. Our approach resolved the protein-level accumulation dynamics in response to HECT E3 KD and revealed extensive UPS-wide redundancy. Interestingly, our data also revealed compensation amongst E3s, as well as compensation of individual protein synthesis rate, in response to KD-induced accumulation. Statistical, machine learning and network analyses revealed unique functional niches and hierarchy of KD response for individual HECT E3s. As a mechanistic proof of principle, we show compensation along an uncovered HUWE1, HECTD1 and TRIP12 compensation axis that affects MCL1 turnover and mitochondrial quality control. Overall, we here provide a high-confidence list of putative HECT E3 substrates and dependencies along with the interaction network that can be explored via an interactive tool. Our analysis framework is adaptable to larger screens and may positively inform the targeted protein degradation field towards precise therapeutic targets.

## INTRODUCTION

Cells maintain protein homeostasis (proteostasis), by tightly controlling the different processes along the protein life-cycle, including protein synthesis and degradation via autophagy or the ubiquitin proteasome system (UPS)^1^. The UPS ubiquitously governs 80% of the mammalian cellular proteome through a ubiquitin transfer cascade involving ubiquitin-activating (E1), ubiquitin-conjugating (E2) and ubiquitin ligases (E3) enzymes^2,3^. As a catalyst of the ubiquitin transfer, E3 ligases are believed to confer specificity for substrate degradation. RING (really interesting new gene)-finger, HECT (homologous to E6AP carboxyl terminus), RBR (ring between ring) and U-BOX are the four key E3 families that together account for >600 enzymes, influencing over 250,000 ubiquitination sites in the human proteome^4,5^. However, resolving the specificity of E3-substrate relationship has remained a challenge as one E3 can ubiquitinate multiple substrates and a substrate can be ubiquitinated by multiple E3s. Additionally, proteins can be monoubiquitinated at multiple lysine residues or polyubiquitinated at few, further diversifying the distinct biological outcomes^6^. Moreover, E3 architecture and specificity can be modulated by specific adaptors, receptors and scaffold proteins^7^. Mere ubiquitination is not enough, cells also possess ubiquitin editing enzymes called deubiquitinases (DUBs)^8^, and other ubiquitin-like molecules^9^ that contribute to the dynamic nature of cellular adaptation.

Biochemical approaches to identify E3 substrates have largely relied either on genetic manipulation or use of recombinant proteins^10,11^. While scalable, these approaches suffer from off-target effects and do not account for true proteome or proteoform diversity in a cell. Proteomic approaches mainly focus on protein-protein interaction (PPI) mapping techniques like affinity purification, immunoprecipitation, proximity labelling and ubiquitinated-substrate trapping^12–18^. These approaches are labour intensive, low-throughput and struggle to differentiate substrates from non-substrate interactors. Furthermore, these approaches are inherently confined to a steady-state measurement and thus limited in capturing transient interactions and proteome reshuffling. In an alternative approach, azidohomoalanine-pulse chase (AHA) proteomics combined with CRISPR knockout^19^ have been used to efficiently capture the dynamic nature of E3-substrate relationship. This approach has yielded a novel level of insight. However, due to the requirement of cell culture in methionine free media, which perturb protein translation, and click-chemistry for labelled-protein enrichment, which necessitates large cell volumes, this approach has been impractical for higher throughput screening platforms. Substrate/interaction repositories and prediction tools such as Biogrid^20^, Ubibrowser^21^ and DeepUSI^22^ lack the cellular context and are often discordant. Moreover, these E3-substrate interactions are cascading, often mired with spatial and temporal distance between the cause and effect. Thus, the investigation of an E3 activity demands investigation of its dynamic functional landscape rather than individual encounters.

To address this, we combined siRNA induced knockdown screen along with a dynamic proteomics screening of HECT E3 ligases. The 28 annotated HECT E3 ligases in human regulate crucial intracellular pathways such as Wnt, Notch and TGFβ and are implicated in various cancers, viral infections and neurological disorders^23^. Due to evidence of HECT E3 involvement in cell death, tumorigenesis, inflammation and DNA damage response^24,25^, there is an increasing interest and efforts towards deconvolution of HECT E3 activity and efficacy as a therapeutic target^26,27^. With the aim to capture putative substrate accumulation as well as proteome synthesis influenced by each HECT E3, we incorporated pulse stable isotope labelling in cell culture (pSILAC) and quantified the proteome pools over ten, independent tandem-mass tag (TMTpro) multiplexes. Using robust statistics and machine learning, our data extrapolates dynamic proteome-wide behaviour in response to the knockdown of individual HECT E3s and identifies exclusivity and overlap within their functional landscape, including putative substrates. Our findings reveal extensive compensation amongst HECT E3s and translation suppression in response to protein accumulation. In addition, our data support the notion of wide-spread redundancy within the E3 functional landscape. Exploring these redundancies for one such redundancy axis among the HUWE1-TRIP12-HECTD1 E3 ligases by combinatorial knockdown revealed their intertwined regulation of mitochondria quality control. Together, we provide a proof-of-concept for a scalable, proteo-genomic screen in a 96-well cell culture format carried out for the HECT E3 ligase family and revealing the E3-substrate relationships with temporal resolution.

## RESULTS

### HECT E3 essentiality is coupled with proteome turnover and cell proliferation

Numerous *in vitro* and *in silico* strategies have been developed to identify E3 ligase substrates^46,47^. Yet, the essentiality and influence of an individual E3 within a cell is rarely investigated in a proteome wide context. Typically, steady-state protein levels are observed instead of monitoring changes in protein dynamics that reveal changes in enzyme kinetics. To bridge this gap, we carried out a proteomic translation and degradation screen for the HECT E3 ligase family using knockdown (KD) in a 96-well format (Fig.1a). First, we measured global protein turnover kinetics by performing a time-course pSILAC experiment (1-96 h) (Fig.S1a), resulting in 59,797 peptides in total and 20,262 peptides identified in both SILAC channels (Fig.S1b). Protein turnover curves derived from these 20,262 peptides revealed moderate to slow turnover of the 16 identified HECT E3s with apparent half-lives between 8-28 h (Fig.S1c), revealing an optimal KD window of 48-72 h post siRNA transfection coupled with SILAC heavy pulse-labeling during the last 24 h (Fig.1a). In total, 27 HECT were individually targeted for KD based on a commercial library with pooled siRNA guides, interspersed with a pooled non-targeting control as a baseline reference. The plates were then processed based on an optimized sample preparation and data acquisition methods^28^, and the resulting data was computationally integrated to resolve direct and indirect HECT E3-substrate relationships.

**Fig. 1:**
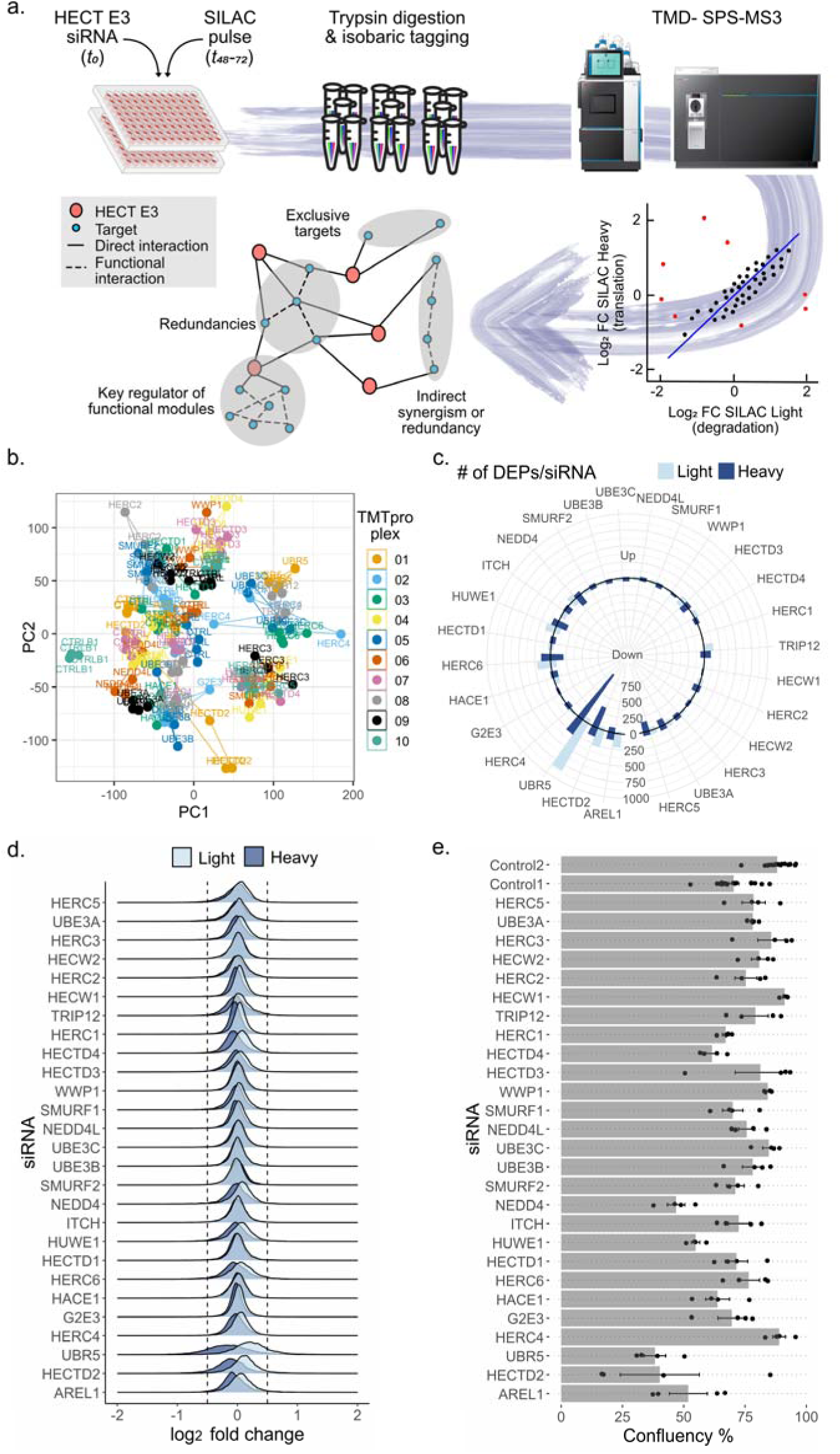
HECT E3 essentiality is coupled with proteome turnover and cell proliferation. **a.** Combined knockdown (KD) and pSILAC screen of HECT E3 ligase workflow. U2OS were subjected to siRNA transfection targeting 27 of the 28 annotated HECT E3s for 24h followed by 24h silencing in light media and 24h heavy SILAC pulse. The samples were digested, labelled with TMTpro, analyzed with LC-MS and using Targeted mass difference (TMD) triggered SPS-MS3. Log_2_FC were calculated per TMT plex with internal control and integrated into robust statistics and machine learning pipeline to identify overlapping and exclusive HECT E3 modalities. b. Principle component analysis of all biological replicates for siRNA HECT E3 KD and non-targeting controls (CTRL), colored based of TMTpro plex ID. c. Number of differentially up-(outwards from 0) or down-regulated (inwards from 0) light and heavy proteins for each KD vs control comparison, log_2_FC cutoff ± 0.58 q value < 0.05. d. distribution of log_2_FC (vs control) in light and heavy protein population and e. Cellular confluency at the time of harvest for each HECT E3 KD and non-targeting controls. Shown are mean values with standard deviation.

On average, we quantified ∼8,000 proteins in both light and heavy populations, supported by >75,000 peptides each (Fig.S1d-e), and confirmed effective KD across all detected and quantified targets (Fig.S1f). Principal component analysis revealed homogeneous distribution of the biological replicates across all siRNAs and TMTpro plexes (Fig.1b), indicating unbiased quantitative measurements. In total, 6,338 light proteins and 6,551 heavy proteins were quantified across all 27 KDs (Fig.S1g-h). The number of differentially expressed proteins (DEPs) (log_2_FC cut off ±0.58, q value < 0.05) varied substantially, with UBR5 KD yielding the most (n>1,000) and HERC1 KD yielding the least (n=14) DEPs (Fig.1c) (Table.S2). For a majority of KDs, the light and heavy proteome fold changes were symmetrically distributed around log_2_(0), indicating balanced proteome turnover and minimal impact of the KD on cellular proteostasis. Interestingly, increased accumulation of light proteome and reduced synthesis of heavy proteome was observed in KD of AREL1, HECTD2, UBR5, HUWE1, NEDD4 and HECTD4 (Fig.1d). This reveals a highly coordinated translation adaptation in response to substrate accumulation upon specific HECT E3 KD. These same KDs also showed reduced cell confluency, suggesting a systemic reduction in cell proliferation and/or viability (Fig.1e) linked to proteostasis imbalance. Reduced viability or proliferation with UBR5, NEDD4 or HUWE1 KD have been reported previously in different cell lines and animal models^48–53^. Collectively, our data links HECT E3 essentiality to cellular viability through measurement of global proteome translation in response to the KD of HECT E3 ligase, with compelling concordance to known HECT E3 phenotypes.

### Many UPS-related processes are co-regulated by HECT E3s

Our data captured dynamic expression from light to heavy proteomes, broadly categorized into six clusters (Fig.2a). DEPs within ±25% absolute fold change difference between light and heavy were classified as following a linear up- and down-regulation. DEPs that fell outside this heavy-light difference range threshold were classified into four non-linear categories (namely *stabilized, reverse stabilized, late response* and *depleted*), reflecting a divergence between the early (light) and late (heavy) proteome abundance (Fig.2a). This categorization allowed us to identify proteins that fell within the same cluster across multiple KDs, signified by their mean log_2_FC in Fig.2b. Specific proteins recurred across up to 11 of the 27 HECT E3 KDs. Amongst the top five shared candidates within each cluster, we found prominent known HECT E3 interactors/substrates being differentially regulated, notably H1-1^54^, CDKN1A^55,56^, TGFβI^57^, DCAF5^20,58^. Additionally, predicted HECT E3 substrates^59^, such as PALM3, ANTXR1, COL11A1 and PLCD1, were also experimentally identified to be differentially regulated in multiple KDs.

**Fig. 2:**
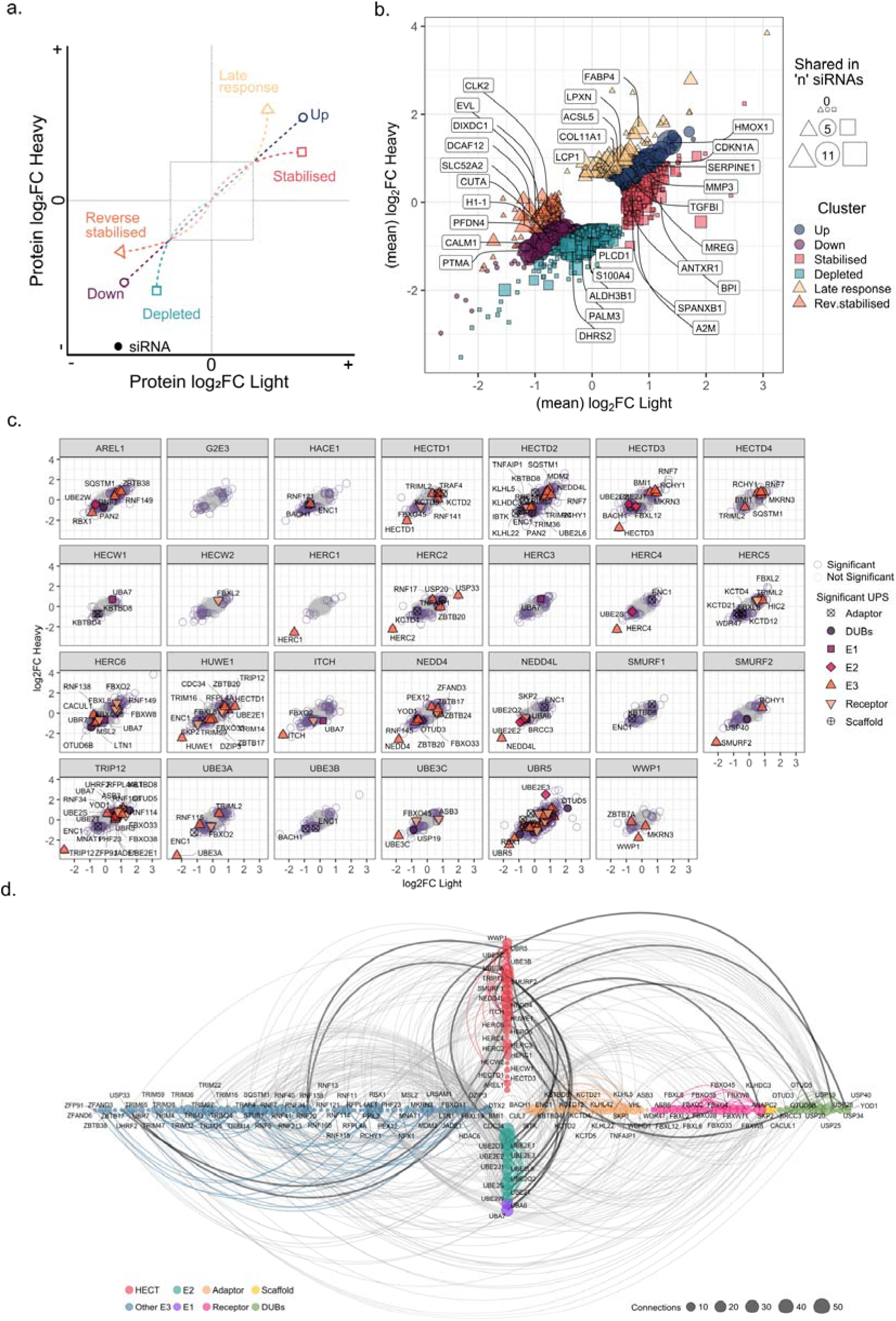
Many UPS-related processes are co-regulated by HECT E3s. a. Graphical representation of protein distribution into six clusters based on log_2_FC of light and heavy channel along x and y axis respectively. b. distribution of significant proteins (cut off log_2_FC ±0.58 and q value < 0.05) amongst six clusters as defined in Fig.2a and color and shape coded. Only proteins quantified in both SILAC channels were considered and difference in log_2_FC heavy and light were calculated to classify the protein into up, down, stabilized, depleted, late response and reverse stabilized clusters. For proteins falling in same cluster across multiple KDs, mean log_2_FC is plotted with size of the shape representing the number of shared KDs. c. Differentially expressed UPS-related proteins (adaptors, DUBs, E1, E2, E3, receptors, scaffold) with significance threshold of log_2_FC ±0.58 and q value < 0.05 used for assigning color and shape across KDs. d. All inter-(grey) and intra-class (colored) STRING annotations for differentially expressed UPS components from Fig.2c. Size of the circle represents the total number of connections annotated at scoring threshold of 400 (STRINGDB version 12.0) and thick black lines indicate STRING connections with observed significant foldchanges in Fig.2c.

Motivated by both substantial overlaps in identified DEPs, over-representation of folding, sorting and degradation related processes in shared DEPs^60^ (Fig.S2a-b), and prior evidence of redundancy and co-dependence within the UPS^10,61–63^, we examined individual UPS components – E3s, E2s, E1s, adaptors, receptors, scaffold, and deubiquitinases (DUBs) (based on annotation from Dutta et al.^64^ and Xue et al.^65^). Strikingly, multiple HECT and other E3s showed a co-dependency and changed significantly in response to the KD of individual HECT E3s (Fig.2c). Specifically, within HECT E3s, HECTD1 and TRIP12 increased in HUWE1 KD, NEDD4L increased in HECTD2 and UBR5 KDs, and UBR5 increased in TRIP12 KD suggesting either de-repression due to KD or redundant compensatory mechanism within HECT E3s. This is consistent with the evidence of co-regulation between HUWE1-TRIP12^66,67^ and TRIP12-UBR5^68,69^. Interestingly, non-HECT E3s, predominantly from the RING finger family (e.g., RCHY1, RNF168, MDM2, TRIML2, RBX1), E1 (UBA7), E2s (UBE2T, UBE2L6, UBE2E2, UBE2E1) and DUBs (PAN2, OTUD6B, OTUD5) either increased or decreased across multiple KDs (Fig.2c), showing a tight link between different components of the ubiquitin system. The observed increase in the mTOR regulatory DUB OTUD5^70^ in TRIP12 KD and UBR5 KD is coherent with similar observations by Morita et al^61^. Adaptors such as ENC1 and BACH1 also follow a similar expression pattern across multiple KDs as observed for E2s and E3s above, supporting the notion of UPS-wide crosstalk. Incorporating STRINGDB^71^ annotations onto these specific UPS related DEPs and HECT E3s shows an extensive network of intra- and inter-class connections within the UPS (Fig.2d) and reveal a highly coordinated regulation of interconnected UPS components.

As anticipated, we also observed significant accumulation or reduction of many annotated interactors and substrates of HECT E3s (based on BioGRID^20^, Ubibrower^59^ v2.0 and DeepUSI^22^) (Fig.S2c-d, Table.S3). However, a majority of the predicted and annotated HECT E3 substrates remained below the fold change threshold, further concretizing the evidence of compensatory mechanism within the UPS. This compensatory phenomenon reflects the redundancy that benefits the broader UPS plasticity, although an appreciable amount of exclusivity is maintained. Overall, our data suggests that cellular proteostasis is ensured largely through regulation of many overlapping UPS components by different HECT E3s, either individually or in concert.

### Robust statistics and machine learning resolves the exclusivity in HECT E3-substrate relationships

To address the substantial overlap in DEPs across HECT E3 knockdowns and resolve exclusivity in the HECT E3 functional landscape (i.e., reveal functional niches of individual HECT E3s), we chose to incorporate robust z statistics and machine learning (ML) onto the log_2_FC data. We applied four robust z-estimators^36^ across all KDs to allow independent, SILAC channel specific detection of anomalous proteins across KDs. The four z-estimators showed 57% overlap globally (Fig.S3a), and filtering for significance (DEPs) further increased the overlap to 85% with maximum outliers detected by MAD (Fig.3a). These outliers showed 3-to-23-fold enrichment of annotated substrates and interactors (from Fig.S2c-d) within the top 25-to-1% of the z scores respectively, supporting their validity (Fig.3b). The identified outliers for light and heavy (Fig.3c, Fig.S3b) were largely unique to each HECT E3s rather than shared, with UBR5 comprising the highest number of statistically significant outliers as represented in the light proteome (Fig.3d) and HERC1 being the lowest. Although the direction of regulation for the outliers is heterogenous across HECT E3s subfamilies (namely HERC, NEDD4 and others^72^), most KDs showed outlier accumulation in the light proteome (Fig.3d), consistent with the anticipated substrate behavior.

**Fig. 3:**
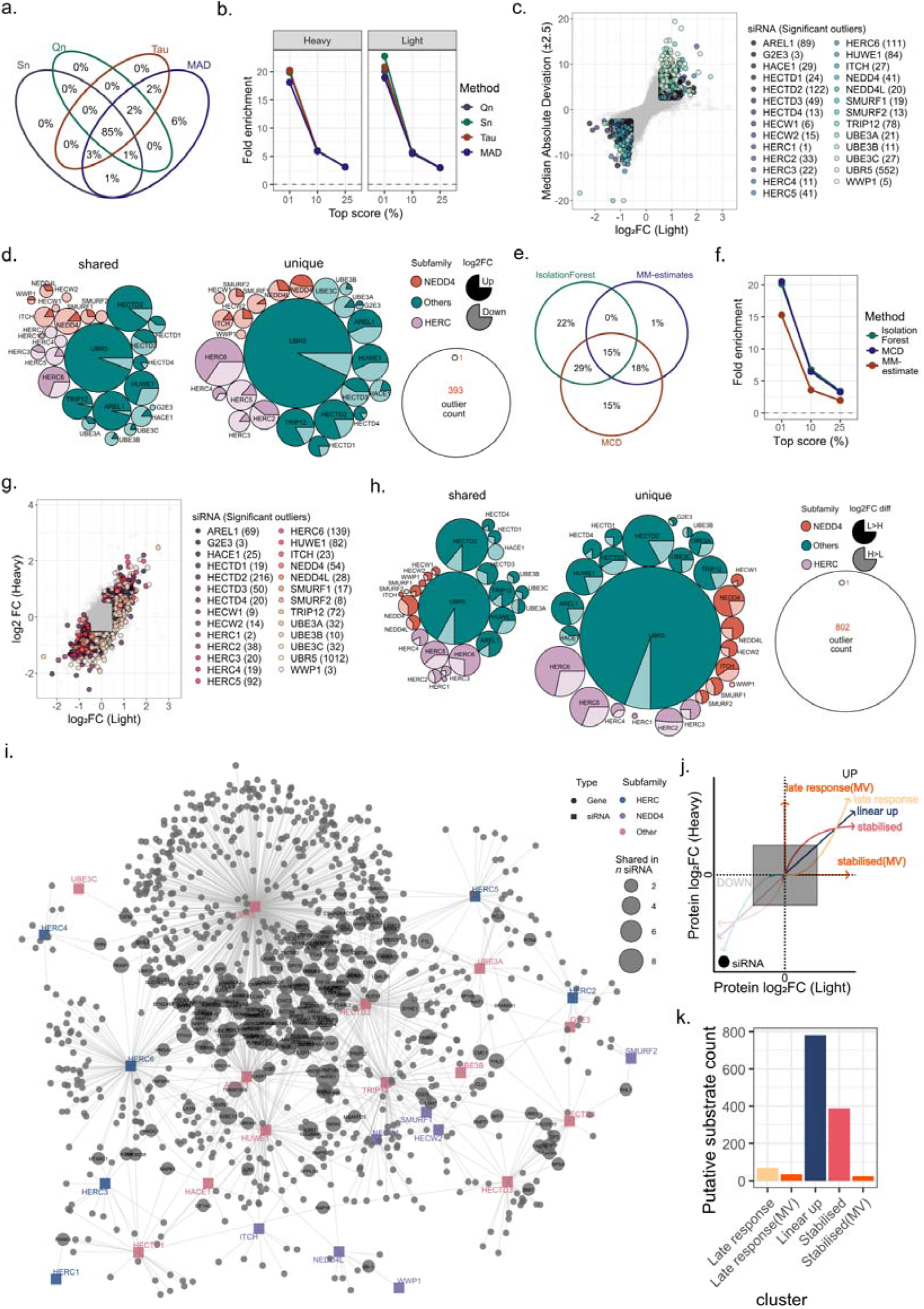
Robust statistics and machine learning resolves exclusivity in HECT E3-substrate relationships. a. Overlap between four z statistical methods (i.e. Sn, Qn, Tau and MAD) for filtering outliers from significant DEPs per SILAC channel. The percentages are rounded off. b. Enrichment of annotated interactors and substrates (from Fig.S2c-d) of HECT E3 ligases in top 1,10 and 25% of the outlier score per SILAC channel. c. Distribution of significant DEPs (log_2_FC ±0.58 q value < 0.05) for light proteome on x-axis versus corresponding MAD score on y-axis. Dots colored for individual siRNA and above log_2_FC, q value and z threshold. Legend indicates the number of outliers per knockdown. d. Number of shared and unique outliers based on MAD (light) for each HECT E3. The size of the circle corresponds to the number of outliers, and pie corresponding to the direction (upregulated = dark and downregulated = light), colored by HECT E3 subfamily. e. Overlap between MM-estimates, Isolation Forest and MCD based machine learning algorithms in identifying outliers from significant DEPs in light and heavy proteome. f. Enrichment per algorithm for annotation substrates in top 1,10 and 25% scoring outliers. g. Distribution of light and heavy log_2_FC of outliers identified by machine learning with number of significant (DEP) outliers/siRNA in legend. h. Number of machine learning based shared or unique outliers with circles indicating the total number per siRNA, pie indicating whether log_2_fc light > heavy (dark) or light < heavy (light), colored by HECT E3 subfamily. i. Network plot of upregulated outliers (dot) (n =1,300) per HECT E3 siRNA (square, colored by subfamily). The size of the dot indicates whether the protein is a shared outlier between HECT E3s or unique. Shared outliers are labeled by their gene name. j. Illustration of putative substrates clusters along log_2_FC light-heavy on x and y axis respectively. (MV) indicates missing value in one of the two SILAC channels. k. Corresponding putative substrate count.

To better define the exclusivity in HECT E3 functional landscape and capture proteins behaving dynamically over the duration of a HECT E3 KD, we relied on semi-supervised (MM-estimates^37^) and unsupervised (Isolation Forest^38^ and Minimum covariance determinant^40^ [MCD]) ML algorithms. MM-estimates attempts to identify off-trend datapoints along a regression, whereas MCD identifies points that are furthest from the robust centre and isolation forest identifies points in sparse regions across the two-dimensional space, allowing the three approaches to capture the diversity of outliers expected from the cumulative dataset. Biologically, these dynamic outliers signify adaptation in translational rates in response to the KD, with the difference in magnitude nearly exclusive to individual HECT E3. The threshold for outlier scores for each algorithm was set to ∼98% specificity (corresponding to 2% false discovery rate) based on ROC curves of all proteins against the pre-annotated ‘ground truth’ outliers (Fig.S3d). As anticipated, the overlap between the three algorithms was low (Fig.S3c, Fig.3e) with 99% of the significant dynamic outliers captured by either MCD or IsolationForest, whereas MM-estimates identified an additional of 1% outliers with poor sensitivity/specificity performance (Fig.3e, Fig.S3d). MCD and isolation forest showed 3-to-20-fold enrichment of known HECT E3 interactors and substrates in top 1 to 25% to flagged outliers, whereas MM-estimates showed a slightly lower, 2-to-15-fold enrichment (Fig.3f). Our data shows that unsupervised, non-regression-based algorithms may be better suited to capture the dynamics in proteome response as opposed to supervised, regression-based MM-estimates. Together, the three ML algorithms allowed us to incorporate the temporal resolution obtained from both quantified (light and heavy) SILAC channels within a single framework and identify protein dynamics under KD conditions (Fig.3g). As with *z* statistics, the ML-based outliers were also predominantly unique to each HECT E3 and increased in the light proteome as compared to heavy (Fig.3h), with 66% overlap between the two approaches (Fig.S3e). The two approaches in conjunction led to filtering of 56% of DEPs shared amongst majority KDs and identification of individual HECT E3 niches amounting to 3,077 putative targets (Fig.S3f). Crucially, z statistics showed a bimodal distribution, influenced by the extremes in light and heavy log_2_FC, whereas ML approaches resolved subtle differences in dynamic protein abundance and may be leveraged for measuring changes in smaller dynamic range below log_2_FC cutoffs (Fig.S3g). Of the 3,077 log_2_FC filtered targets, UBR5 accommodated the highest (1,012) and HERC1 the lowest (2) number of dynamic outliers (Fig.3g-h). Collectively, these outlier candidates summarize the distinct functional landscape of each HECT E3, with the least promiscuous substrates, partners, adaptors and overall proteasomal influence within the cell.

Given that we anticipated an increase in HECT E3 substrate stability upon KD, we looked specifically at upregulated outliers and mapped the points of overlap amongst HECT E3s. In total, 1,300 proteins were upregulated outliers across all HECT E3s, with 571 belonging to UBR5 alone, including known UBR5 substrates, such as MAFF^73^, MYC^74^, ATAD2^75^, and KATNA1^76^. UBR5 also accounted for the majority of shared outliers across E3s (*n =* 160), indicating a cross-subfamily synergism (Fig.3i). Reclassifying the upregulated outliers into putative substrate clusters (same as Fig.2a but also including proteins quantified only in one SILAC channel) (Fig.3j) showed that the majority were either stabilized or exponentially upregulated (Fig.3k). Remarkably, the number of shared candidates in stabilized clusters were reduced by 70% based on outlier scoring (Fig.S3h) when compared to fold change threshold-based DEPs (Fig.2b). Incorporation of STRING annotation onto the upregulated outliers for each HECT E3 siRNA provides a layer of association and delineates secondary effects from actual substrate accumulation, further reducing the putative substrate list (example Fig.S3i, supplementary file S1, Table.S2). In summary, we provide a *bona fide* list of putative HECT E3 substrates and associated proteins. Additionally, we extend this analysis to characterize unique HECT E3 dependencies from the down regulated pool of DEPs (Fig.S3j), with STRING annotation for causal resolution (supplementary file S2, Table.S2). Our data uncovers an extensive crosstalk within HECT E3 family that suggests an underlying signaling hierarchy and compensation across HECT E3s. We also provide an interactive tool that allows exploration of the data, protein-wise outlier trends and STRING network for both up and down-regulated outliers together (zenodo ID: 23885440).

### Combinatorial knockdown reveals crosstalk along the HUWE1-TRIP12-HECTD1 axis

Knockdown of HUWE1 (also known as mule/ARF-BP1) increased HECTD1 and TRIP12 levels (Fig.2c, Fig.S3i), amongst other known interactors and substrates such as TGFBR2^77^ (Table.S3, Fig.S4a). However, many prominent HUWE1 substrates, including the mitophagy and apoptosis driver MCL1^78,79^, showed no accumulation upon HUWE1 KD (Fig.S4a). Similar observations were made by Monda et al^80^., wherein a 16-fold HUWE1 depletion was insufficient to induce accumulation of known substrates in 293T and HAP1 cells^80^. The total (Fig.S4b) and phosphorylated MCL1 (Fig.S4c), a prerequisite for MCL1 degradation during mitophagy^78^, remained at the same abundance in HUWE1 KD when compared to control KD, indicating that MCL1 degradation may be controlled by E3s other than HUWE1. TRIP12 and HUWE1 are co-expressed, share a WWE domain, and collaborate in substrate degradation^66,67,71^.

We therefore asked whether the increase in HECTD1 and TRIP12 upon HUWE1 KD compensated partially towards HUWE1 substrate clearance, especially MCL1. To that end, we performed a comprehensive transcriptome, proteome and phenotypic investigation of HUWE1, HECTD1 and TRIP12 KD, individually and in combination (Fig.4a), to assess overall changes and particularly for MCL1 and its potential effects on mitophagy and mitochondria. Across whole cell and specifically the mitochondrial proteome, a significant decline in heavy-labelled proteins was observed in triple KD as compared to HUWE1 KD alone (Fig.4b), suggesting reduced translation. Focusing on the proteins and the transcripts synergistically regulated in HUWE1 combinatorial KDs (i.e., HUWE1+HECTD1, HUWE1+TRIP12, HUWE1+HECTD1+TRIP12; collectively referred to as HUWE1+), we identified 458 proteins in light (Fig.4c, Table.S4), 503 proteins in heavy (Fig.4d, Table.S5), and 472 transcripts (Fig.4e, Table.S6) to be differentially regulated specifically in HUWE1+ only. Notably, MCL1 abundance remained close to basal level in light proteome and transcript but significantly increased threshold when carrying out double (HUWE1+HECTD1; log_2_FC: 0.75) or triple (HUWE1+HECTD1+TRIP12; log_2_FC: 0.76) KD in the heavy proteome, contrasting the overall cellular and mitochondrial proteome response (Fig.4b, d). Interestingly, GSK3B, a kinase responsible for MCL1 phosphorylation and degradation^81,82^, showed a significant increase in the light proteome of double HUWE1+HECTD1 and triple HUWE1+HECTD1+TRIP12 KD (Fig.4c), consistent with persistent MCL1 turnover in early stages of combinatorial KD. We validated the GSK3B increase in light and MCL1 accumulation in heavy proteome by orthogonal parallel reaction monitoring (PRM), confirming concordant increase upon double and triple KD (Fig.S4d, Table.S7). Outside mitochondria, HAPSTR1, a cytoplasmic and nuclear cofactor for HUWE1 mediated control over stress signaling pathways^80,83,84^, also showed a similar trend (Fig.S4d, Table.S7), suggesting consistent compensatory HUWE1-HECTD1-TRIP12 cross-talk across sub-cellular locations.

**Fig. 4:**
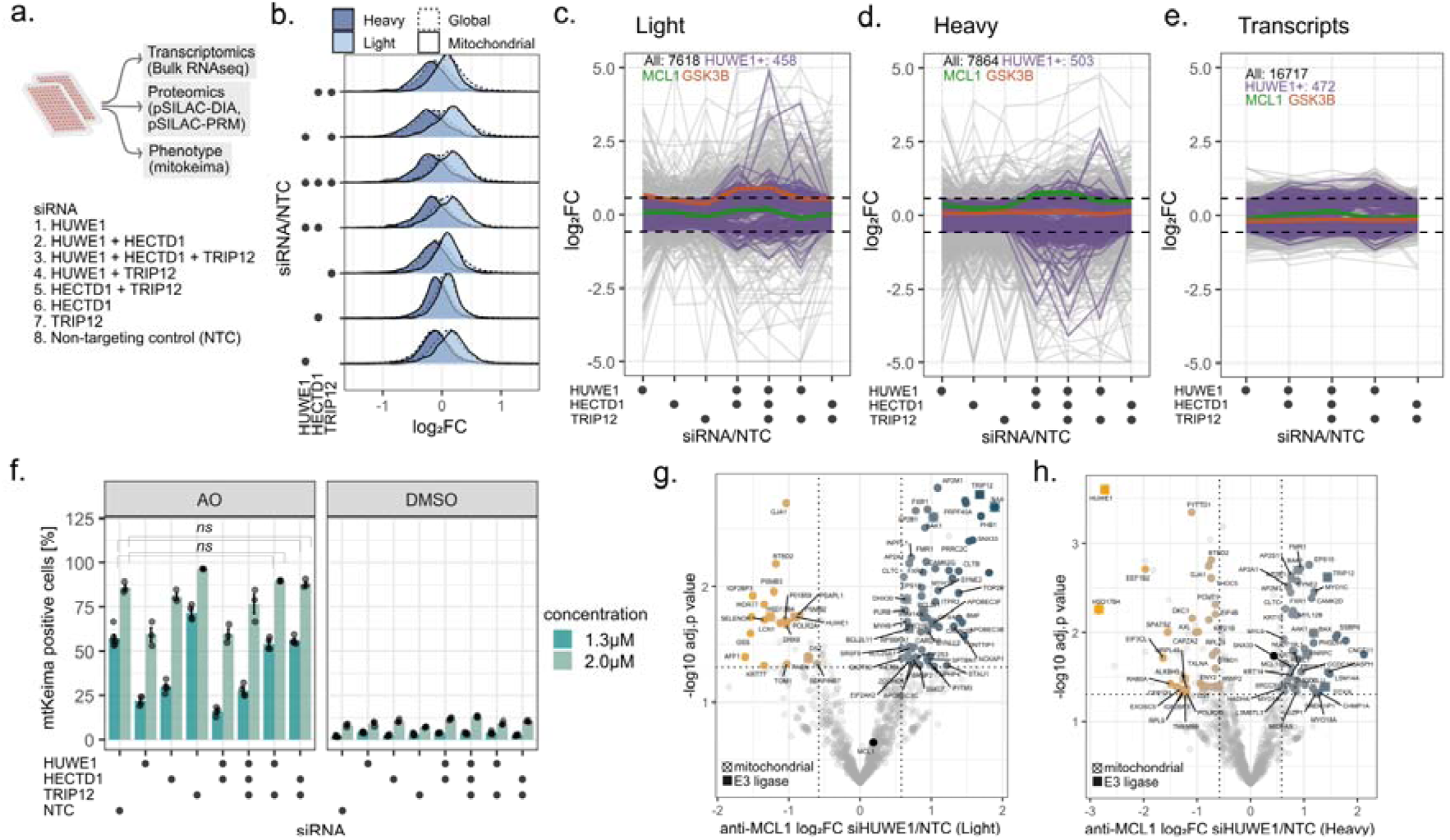
Combinatorial knockdown reveals crosstalk along the HUWE1-HECTD1-TRIP12 axis. a. Schematic of siRNA induced KD of individual or combination of HUWE1, HECTD1 and TRIP12, and experiments performed for combinatorial effect characterization. All the transfections were performed with the same timeline as Fig.1a, with 24 h silencing window in light media followed by 24 h heavy media culturing. b. Ridge plots of log_2_FC distribution in light (light blue) and heavy (dark blue) proteome for KD vs non-targeting control (NTC) in whole cell proteome (dotted) and mitochondrial subset (solid) (mitocarta 3.0) respectively. (Kolmogorov-Smirnov distance = 0.07 and 0.23, and FDR corrected p value: 4.6×10^-18^ and 8.5×10^-20^ for whole cell and mitochondrial subset respectively). c-e. Spaghetti plot with log_2_FC across comparisons in light and heavy proteome, and transcriptome respectively. Grey represents all quantified proteins/transcripts, purple represents differentially regulated proteins/transcripts specifically in HUWE1 combined with HECTD1, TRIP12 or both (denoted as HUWE1+) (log_2_FC >±0.58 q value < 0.05 in HUWE1 combinations and log_2_FC <±0.58 in individual KDs or HECTD1 + TRIP12) along with MCL1 (green) and GSK3B (orange). f. mtKeima positive HeLa-PARKIN cells with combination of KDs and mitophagy induction by antimycin-oligomycin (AO) treatment at the concentration of 1.3 or 2 µM each, DMSO as control at corresponding AO concentration, measured with flow cytometry. Shown are mean values with standard deviation, significance is derived by Dunnett’s test (n=3 biological replicates for 2 μM, n=4 biological replicates for 1.3 μM, n=6 biological replicates for NTC) and only non-significant conditions (p value >0.05) are labelled as ‘ns’. g-h. MCL1 interactome in HUWE1 KD cells compared to NTC in light and heavy isotopic label respectively. Only proteins significant (log_2_FC ±0.58, q value <0.05) in anti-MCL1 and not in anti-IgG pulldown are colored. E3 ligases and mitochondrial proteins are shaped square and cross respectively, MCL1 as black.

The differentially expressed HUWE1+ proteins and transcripts showed a poor overlap (Fig.S4e) and both light and heavy proteome showed poor correlation against transcriptome, indicating that the observed effects are predominantly post-transcriptional (Fig.S4f). Most E3s barring TRIML2 and the siRNA-induced KD itself, also followed a post-transcriptional regulation, including HUWE1-HECTD1-TRIP12 and TRIP12-UBR5 expression pattern observed in the initial TMT screen (Fig.3c, Fig.S4f). Co-regulated HUWE1+ proteins were enriched for mRNA splicing and cell cycle processes (Fig.S4g-i), which is in agreement with previous studies^79,85–88^, and the transcripts predominantly showed increased extra-cellular matrix organization related genes (Fig.S4l).

Considering the mitochondrial subset of proteins in HUWE1+ DEPs, we observed a contrasting trend in light and heavy proteins (Fig.S5a) whereas the transcripts remained stable below log_2_FC cutoff for all except HUWE1+TRIP12 KD (Fig.S5b). While the genes encoding the mitochondrial proteins and transcripts showed poor overlap (Fig.S5c), the overarching pathways were consistent across the proteome and transcriptome, showing increased mitochondrial protein translation in early KD and decreased protein and transcript abundance at the later stage (Fig.S5d-f). Located on mitochondrial outer membrane, MCL1 is known to undergo rapid turnover, essential for driving mitochondrial dynamics and apoptosis^89–94^. Downregulation of mitochondrial protein degradation, import and respiratory electron transport along with accumulation of MCL1 (Fig.4d, Fig.S4d, Fig.S5e) further hints towards impaired MCL1 clearance in HUWE1+ depleted cells with consequences on mitochondrial quality control.

To investigate the functional consequences of impaired MCL1 clearance on mitophagy regulation, we treated HeLa mt-KEIMA Parkin expressing cells with antimycin-oligomycin (AO) to induce mitochondrial dysfunction and promote mitophagy. Notably, we observed reduced mitophagy induction in HUWE1 and HECTD1 KD (Fig.4f). Although MCL-1 turnover was not significantly altered in single KDs under basal conditions (Fig.4c, d), insufficient turnover of MCL1 under stress conditions due to the absence of either E3 ligase could contribute to impaired mitophagy induction. In line with this, the combinatorial HUWE1+HECTD1 KD further reduced mitophagy induction (Fig.4f), highlighting the compensatory axis of HUWE1 and HECTD1 through accumulation of MCL1. Strikingly, TRIP12 single knockdown elevated AO-induced mitophagy, which was however restored to control levels with combinatorial knockdowns of TRIP12 with HUWE1 and HECTD1 (Fig.4f). This is in line with insignificant MCL1 accumulation under TRIP12 single or combined KD (Fig.4c,d) and indicates additional changes that influence mitophagy induction upon TRIP12 KD.

To confirm whether other E3s, especially HECTD1 and TRIP12 interacted with MCL1 under HUWE1 KD, we performed MCL1 co-immunoprecipitation (coIP). Interestingly, MCL1 showed a 2.6-to-3-fold increased interaction with TRIP12 under HUWE1 KD (Fig.4g-h, Table.S8). Moreover, other HECT E3s, namely WWP2, UBR5 and ITCH, were also enriched in MCL1 pulldown compared to IgG control, along with other non-HECT E3s (Fig.S5g-h) suggesting a multi-faceted E3 control over MCL1. Relative to IgG, the MCL1 enrichment in NTC and HUWE1 KD were comparable (Fig.S5g-h) and in line with our previous observations (Fig.4b-c, Fig.S4a), there was no substantial increase in MCL1 enrichment upon HUWE1 KD (Fig.4g-h). Additionally, ubiquitin enrichment in NTC and HUWE1 KD was also consistent in anti-MCL1 pull-down, suggesting unperturbed MCL1 ubiquitination under HUWE1 KD via other E3s (Fig.S5g-h). As we did not detect HECTD1 in MCL1 coIP under HUWE1 KD, this may reflect a highly transient or a TRIP12-dependent interaction between MCL1 and HECTD1 (Fig.S5i). HUWE1 KD instead led to increased MCL1 interaction with pro-apoptotic mitochondrial proteins BAK1/BAX and other apoptosis factors such as BCL2L1, BCL2L11, DYNLL2 and BMF (Fig.4g-h, Fig.S5i). In contrast, we did not observe any notable accumulation of these apoptotic markers in whole cell data (DIA, TMT or PRM), suggesting increased anti-apoptotic activity of MCL1 under HUWE1 KD. Amongst the differentially enriched MCL-1 interactors in HUWE1 KD (Fig.S5i), clathrin-mediated endocytosis and BCL2/MCL1-mediated apoptosis were the key pathways linked to TRIP12-HUWE1 cascade.

Together, our data implicates TRIP12 and HECTD1, in addition to previously reported HUWE1 involvement^78^ in MCL1 degradation, mitophagy induction and apoptosis. Crucially, the hierarchy of take-over remains to be resolved, as HUWE1+HECTD1 KD showed increased propensity for MCL1 accumulation (Fig.4d, Fig.S4d) with evident loss of mitophagy (Fig.4f), while TRIP12 appeared to interact with MCL1 in HUWE1-dependent manner (Fig.4g-h). The role of TRIP12 is further confounded by increased UBR5 expression (Fig.2c) and mitophagy upon TRIP12 KD (Fig.4f), in-line with UBR5 enrichment in MCL1 co-IP (Fig.S5g-h) and suggests that UBR5 may be the key arbiter of the observed phenotypes (Fig.4f, Fig.S5i). Unsurprisingly, MCL1 (Fig.S5j-k) and phosphorylated-MCL1 (Fig.S5l-m) abundance remained consistent under HUWE1 and HUWE1+TRIP12 KD, confirming unimpaired turnover of MCL1 with TRIP12 KD. Collectively, using combinatorial KDs and orthogonal assays we explored the compensatory axis of MCL1 degradation in HUWE1 knockdown, highlighting a multi-faceted control over MCL1 turnover. Importantly, our observations corroborate that proteostasis is maintained through overlapping niches of E3s.

## DISCUSSION

Investigation of E3 ligases has largely focused on resolving substrate specificity. This is crucial, especially in the targeted protein degradation (TPD) field, wherein specificity determines druggability and therapeutic scope. However, only over 2% of the 600+ E3 repertoire is currently utilized in TPD based therapeutic strategies^95^. This limited utilization is partly due to the fact that the existing experimental^12–18^ and computational^21,22,96,97^ approaches predominantly capture steady-state protein-protein interaction, undermining the redundancy in E3-substrate pairs and dynamic nature of protein-protein interactions. As an orthogonal approach to conventional PPI based annotation, we here focused on characterizing dynamic proteome-wide influence of E3s, specifically the HECT E3 family.

Our setup allowed a turnover-informed KD screen and measurement of proteome synthesis-degradation dynamics following HECT E3 KD (Fig.1a). This supported temporal resolution of protein accumulation, relative to each individual HECT E3. Moreover, multiplexing with TMTpro allowed for intra-plex experimental controls and minimization of batch effects or missingness. The quantitative precision and depth of peptide and protein coverage were afforded by intelligent data acquisition^28,29^ and can be further reconciled with increasing pSILAC time points for more resolved turnover dynamics^98^, and TMT35plex^99^ for higher throughput. Alternatively, as exemplified in combinatorial KDs (Fig.4b-c, Fig.S4f), ultra-fast mass spectrometer assisted pSILAC-DIA may be leveraged to collate KD induced proteome dynamics with increased longitudinal and conditional coverage^100–103^. Crucially, our approach allowed us to quantify the proteome accumulation and translation affected by individual HECT E3 KD, and to link HECT E3 essentiality and cellular viability with proteome response (Fig.1d-e). HECT E3s are increasingly being adopted in the TPD field^104,105^ and our dataset may aid define the tolerance and expanse of particular E3 enzymes’ engagement for protein degradation.

Our findings corroborate the widespread redundancy within the UPS^63,95,106^ (Fig.2c-d) and identify functional influence of individual HECT E3s in the U2OS proteome, characterized by distinct protein translation/accumulation dynamics (Fig.2b). The substantial overlap of temporally resolved DEPs across multiple HECT E3s (Fig.2b), together with crosstalk across the UPS (Fig.2c-d), suggests that the UPS achieves robustness through dispersal of ligases activity. Shared substrates have been reported previously, for instance, within the NEDD4 subfamily^107^, and between HUWE1 and TRIP12^67^. Our approach further extends this notion by looking at proteome niches of individual HECT E3s, including where they overlap and diverge (Fig.3i, Fig.S3j).

One possible explanation for the overlap is KD-induced transcriptional adaptation^108^. However, we observed poor correlation in our comparison of proteomes and transcriptomes (Fig.S4f), suggesting that observed effects may be specific to individual KDs and are largely post-transcriptional. This is unsurprising, as mRNA and protein abundances seldom correlate^109,110^. TRIML2 was an exception (Fig.S4f) and may be an interesting E3 candidate towards understanding the fundamentals of UPS adaptation transcriptionally. Another possible cause of apparent overlap in DEPs could be siRNA transfection off-targets or statistical noise in the data rather than true co-regulation. By incorporating robust z statistics and machine learning supported by ground-truth annotations, we mitigate precisely this statistical noise and delineate the exclusivity from redundancy within HECT E3 influence network (Fig.3b-c, f-g). Thus, our approach defines high-confidence functional niches for individual HECT E3s and highlights their essentiality in proteostasis. Leveraging STRING interaction annotations, we further resolve the upregulated targets and downregulated dependencies to summarize the interaction cascade of each HECT E3 KD (Supplementary file S2-S3). These interaction networks allowed us to separate primary response from downstream effects and selectively isolate unique, overlapping proteins that support compensatory dispersal of HECT E3 activity. The networks can be explored in an interactive format through a local .html tool available on zenodo (ID: 23885440).

Our data shows UBR5 at the core of HECT E3 essentiality, with many shared modalities with other HECT E3s (Fig.3i). Thus, UBR5 may serve as an ideal, broad footprint E3 ligase for TPD and pharmacology. Notably, there is increasing interest in UBR5 as a therapeutic target for regulation of key transcription factors^111,112^. Conversely, among the HECT E3s with a particularly narrow influence range identified in our dataset (Fig.3g), HERC4 has been leveraged for small molecule induced degradation of STING^113^.

Mechanistically, we focused on HUWE1 and observed a compensatory, rheostat function of TRIP12 and HECTD1 upon HUWE1 KD, thereby ensuring cellular proteostasis (Fig.4b-d). Specifically, mitochondrial protein translation, import, respiratory electron transport and mitophagy appear to be regulated through a functional relay between HUWE1, TRIP12 and HECTD1 (Fig.S5d-f). We found that MCL1, a key regulator of mitophagy and apoptosis, is regulated through a multifaceted network and interacts with other E3 ligases under HUWE1 KD. Moreover, we suspect that the compensation is not restricted to mitochondria, with similar expression pattern observed for HAPSTR1^80,83,84^, a cytosolic/nuclear target of HUWE1. Other HECT E3 KDs with increased alternative E3 ligase may exhibit similar compensation mechanism (Fig.2c) and are promising candidates for exploring redundancy and multiplicity within the UPS.

In conclusion, our data provides a bird’s-eye view of HECT influence landscape as well as individual protein dynamics within the U2OS context, and highlights a compensatory mechanism among E3 ligases, exemplified by HUWE1-TRIP12-HECTD1 axis. The extent and dynamics of redundancy and exclusivity within the UPS remain to be thoroughly understood and our approach attempts to resolve this by dynamic proteome measurement under KD. Beyond physical interaction, our statistical and machine learning approach allowed identification of targets that may depend exclusively on specific HECT E3s or catalyzed by multiple E3s. Our results pave the way for similar applications of dynamic proteo-genomic screening in cataloguing E3s-substrate relationships and identifying exclusive or overlapping functional niches for accelerated progress in the TPD field.

## ACKNOWLEDGEMENTS

We thank the Center for Functional Proteomics (Goethe University Frankfurt, DFG registered research infrastructure RI_ 00611) for support with mass spectrometry measurements, particularly Martin Adrian All-good and Doha Boutguetait for their technical assistance, and Suleyman Bozkurt for maintaining the analysis pipeline for pSILAC-TMT experiments. This work was supported by the German Federal Ministry of Education and Research (BMBF) within the Clusters4Future PROXIDRUGS program (FKZ03ZU2109FC, to C.M.), Deutsche Forschungs-gemeinschaft (DFG, German Research Foundation) projects 514894665 (TRR 387/1 UbiQancer, C.S-V, M.E. and C.M.), 456687919 (SFB1531 Stroma-vascular damage control, C.M.), under Germanýs Excellence Strategy – EXC 2026, Cardio-Pulmonary Institute (CPI, Project ID: 390649896 to C.M.) and EXC-3094, Cluster of Excellence SubCellular Architecture of Life (SCALE, Project ID: 533751785, to C.M.), the European Research Council under the European Union’s Horizon 2020 Research and Innovation Programme (Grant ERC CoG 101125845, C.M.), and the Leistungszentrum Innovative Therapeutics (TheraNova) funded by the Fraunhofer Society and the Hessian Ministry of Science and Arts. The mass spectrometery was supported by the European Fonds for regional development (REACT-EU, IWB-EFRE-program Hessen, 20008763, to C.M.) for the Vanquish Neo, Orbitrap Ascend LC-MS systems. C.M. also acknowledged support from the Hessian Ministry of Science and Research, Arts and Culture LOEWE Spitzenprofessorship, the Volkswagenstiftung, the Alfons and Gertrud Kassel-Stiftung, the Dr. Rolf M. Schwiete-Stiftung, and the Johanna Quandt-Universitäts-Stiftung (all to C.M.).

## MATERIAL AND METHODS

### Cell culture, siRNA knockdown and SILAC labelling for proteomic screen

96-well cell culture plates were seeded with 6,000 U2OS cells in 100 µL of SILAC Dulbecco’s Modified Eagle Medium (SILAC DMEM) supplemented with 10% fetal bovine serum and light lysine and arginine amino acids (K0R0, Silantes) and incubated overnight in a humidified incubator at 37°C with 5% CO_2_. Following day, cells were transfected with HECT E3 siRNA or non-targeting control (Dharmacon, onTarget plus smartpool) as per the following protocol. To prepare the transfection mix, 2.5 µL of 1 µM siRNA was pipetted into a PCR plate (Greiner bio-one, 652290, Sapphire Microplate) along with 7.5 µL of Opti-MEM (Gibco, 12559099) and incubated at room temperature for 5 mins. Lipofectamine RNAiMAX (Invitrogen, 10514953) transfection reagent was diluted 1:50 in Opti-MEM, incubated for 5 mins at room temperature and equal volume (10 µL) was added to each siRNA mix, followed by 20 min incubation at room temperature. 80 µL of SILAC light media was added to 20 µL of transfection mix. For double and triple knockdowns, same volumetric ratios were followed to make up 25 nM of total siRNA concentration per 100 µL of transfection media. Media in 96-well cell culture plates was aspirated and supplemented with transfection media, followed by 24h incubation at 37°C. Next day, the transfection media was decanted and replaced with SILAC light media for 24 h followed by 3X pre-warmed PBS wash & supplementing SILAC heavy media (SILAC DMEM, 10% fetal bovine serum and heavy lysine and arginine (K8R10, Silantes) for 24h. After a total of 72 h from transfection, the cells were washed 5x with pre-warmed PBS. 75 µL of prewarmed (95°C) lysis buffer (5% SDS, 5 mM TCEP, 20 mM CAA, 100 mM TEAB, 1x Protease inhibitor cocktail) was added per well, sealed with silicon mat and snap-frozen in liquid nitrogen. In parallel, U2OS cells were grown in DMEM light or heavy in 15cm plates and collected after 4 weeks (8 passages), washed with 3x PBS and snap frozen in lysis buffer, hereafter called ‘carrier lysate’.

For transcriptome analysis, immunoprecipitation and FACs experiments, the seeding, culturing and transfection volume was scaled to 4X in 24-well or 10X in 6-well plates relative to aforementioned 96-well protocol. For protein turnover experiment, U2OS cells were labelled with SILAC heavy media for 0, 1, 2, 4, 6, 24, 48, 72, and 96 h in 96-well format as described above.

### Cell lysis and sample preparation for mass spectrometry

The plates were processed as described in Parmar et al^28^. Briefly, cells were homogenized by heating 95°C followed by multi-horn probe sonication. Lysate was digested in 96-well STrap plates with 0.2 µg of Trypsin+Lys-C (1:1) (Promega V5111 and VA1170), eluted and dried in vacuum concentrator. The eluent was resuspended in 1% Acetonitrile, 0.1% Trifluoracetic acid and cleaned up with C18 plates (Affinisep-AttractSPE microelution μ96W-C18-M.T.1.1) and dried again. The peptides were resuspended in 2% Acetonitrile 0.1% Trifluoracetic acid for data-independent acquisition (DIA) at the concentration of 0.125 µg/µL or resuspended in 19µL of 100mM EPPS 5% Acetonitrile for TMTpro labelling. 300 µg of carrier lysates corresponding to light and heavy proteome were processed in the same fashion in STrap mini tube format, cleaned with C18 SepPak cartridges (tC18, 100 mg, Waters), resuspended at the concentration of 0.5µg/µL in 100mM EPPS 5% Acetonitrile and used within TMTpro 18-plex for the screen with channel 126 for as light carrier and channel 135N as heavy carrier. For pSILAC-DIA and PRM experiments, C18 eluents were resuspended in 2% Acetonitrile, 0.1% TFA and adjusted to 0.125 ng/µL and spiked with 0.2x of Biognosys iRT peptide mix.

### TMT-labelling and high pH fractionation

TMT label annotation for each plex is described in table.S1 and labelled as described in Parmar et al^28^. Briefly, 5mg dried TMTpro stocks were resuspended in 100% Acetonitrile and 1 µL was added to 19µL peptides resuspended in 100mM EPPS 5% Acetonitrile and incubated at 24°C for 1 h, followed by addition of 5 µL of 0.5% hydroxylamine and incubated at RT for 15 mins to quench the TMT-labelling. Labeling check was performed by pooling 1 µL of each TMTpro-channel for each of the 10-plexes, analysed by LC-MS and pooling volumes were adjusted to normalized median peptide intensities across all channels. The pooled TMTpro plexes were acidified and cleaned with C18 SepPak (tC18, 50 mg, Waters), dried and resuspended in 5% Acetonitrile 5mM Ammonium bicarbonate at the concentration of 1.5 µg/µL for high-pH fractionation. 45µg of the TMTpro labelled peptides were loaded on to C18 column (XSelect CSH, 1 mm × 150 mm, 3.5 µm particle size; Waters SKU186005251) on-line with Ultimate 3000 HPLC system (Thermo Fisher Scientific) and eluted with a multistep gradient from 3-60% of 5 mM ammonium bicarbonate in 80% acetonitrile over 65 min at a flow rate of 30 µL/min. Eluting peptides were collected every 43 seconds into a total of 96 fractions and concatenated into 24 fractions. Pooled fractions were dried in a vacuum concentrator and resuspended in 2% acetonitrile 0.1% trifluoroacetic acid for LC-MS analysis.

### Data acquisition

pSILAC-TMTpro data was acquired on Orbitrap Ascend online with Vanquish Neo (Thermo Fischer Scientific) in SPS-MS3 mode enabled by target mass difference node and real-time search. Briefly, 400 ng of each peptides fraction was separated over a 120 min gradient (7-25% Solvent B ramp in 90 min, followed by 40% in 30 min and 60% Solvent B in 5 min) on in-house packed 35 cm fused-silica column (75 µm inner diameter) packed with 1.9 µm Reprosil-Pur C18 particles (Dr. Maisch r119.b9.). MS1 scan was acquired in orbitrap at 120,000 resolution, 350-1400 m/z scan range and AGC of 4e5 with selection for charge states 2-5, dynamic exclusion of 90 seconds and selected for target mass difference for SILAC isotopes. MS2 was acquired in iontrap at Turbo setting with isolation window on 0.7 Th, AGC of 2.5e4 and collision induced dissociation fragmentation at 35%. Scan were then passed through real-time search^29^ node for triggering synchronous precursor selection with TMTpro on peptide N-terminus and lysine as fixed modifications and Methionine oxidation, Lysine (+8.0141) and Arginine (+10.0082) as variable modification, XCorr cut-off of 1 and deltaCn 0.1 at 10 ppm. Filtered precursors were selected for MS3 in orbitrap at 45,000 resolution at AGC 7.5e4 and high-energy collision energy of 50%.

pSILAC-DIA data was acquired on timsTOF HT mass spectrometer (Bruker Daltonics, Bremen, Germany) online with Evosep Eno LC system (Evosep, Odense, Denmark). Peptides (500 ng per sample) were loaded onto Evotips according to the manufacturer’s protocol and separated on an Evosep Eno using the standardized 30 samples-per-day (30 SPD) method (44-min gradient). Separation was performed on an Evosep Performance column (15 cm × 150 µm i.d., 1.5 µm C18 beads; EV1137) maintained at 40 °C and coupled to a 20 µm i.d. emitter (Bruker Daltonics). Peptides were ionized in positive mode using a CaptiveSpray source (capillary voltage 1600 V). They were analysed on timsTOF HT operated in dia-PASEF mode under timsControl version 8.

The TIMS ion mobility range was set to 1/K₀ = 0.65–1.45 V·s·cm⁻², with accumulation and ramp times of 100 ms each (100% duty cycle). MS1 and MS/MS spectra were recorded over m/z 100–1700. Each acquisition cycle consisted of one MS1 TIMS ramp followed by eight dia-PASEF MS/MS ramps. These ramps contained 24 isolation windows of 25 Th width with 1 Th overlap, covering m/z 400–1000 across the corresponding ion mobility range. The resulting cycle time was approximately 0.95 s. Collision energy was decreased linearly with decreasing ion mobility, from 59 eV at 1/K₀ = 1.60 V·s·cm⁻² to 20 eV at 1/K₀ = 0.60 V·s·cm⁻². Mass and ion mobility were calibrated before acquisition using three selected ions from the Agilent ESI-L Tuning Mix.

pSILAC-GPF, refinement and targeted PRM was acquired on Orbitrap Ascend online with Vanquish Neo with the same setup as before and all runs (pooled GPF and individual samples, n =6 per condition) were separated over 45mins gradient (5-38% Solvent B ramp in 40 min, 60% Solvent B in 5 min). GPF-DIA data was acquired over ten 65Th windows (350-1000 m/z) with a 50Th Adaptive RT^30^ iontrap scan executed after each orbitrap MS1 and DIA scan at 120K and 30K resolution respectively and high-collision energy of 30%. 584 precursors for target proteins were filtered from GPF library for further refinement via Skyline^31^ daily (version 26.0.9.054) with PRM conductor plugin for automated selection, retention window optimization, precursor list splitting (6 runs), method and .rtbin file generation. Eventually, 355 precursors with variable retention windows were selected for final PRM experiment and acquired with AdaptiveRT based dynamic retention time correction. The data was analyzed with skyline and 289 precursors corresponding to 69 proteins were normalized by total ion current, summarized with Tukey median polish, and MS2 level fold change were calculated with FDR correction. The data and associated skyline document can be found on https://panoramaweb.org/vHdHjQ.url

### pSILAC-TMTpro and -DIA data analysis

pSILAC-TMTpro data was analyzed with Proteome Discoverer (version 2.4). The data was searched against human proteome (Uniprot release 2023_05) and contaminants database (cRAPome) with precursors mass tolerance of 10ppm and MS2 mass tolerance of 0.6Da and MS3 reporter quantities were corrected for TMTpro batch correction factors from the manufacturer. Resulting PSMs files were filtered for high confidence, contaminants, median normalized and separated by isotope labelling with DynaTMT^32^. PSMs with <5 s/n mean ratio were removed and remaining psms rolled up to isotopically distinct peptides. Protein level fold changes were calculated for light and heavy proteins individually using PBLMM^33^. pSILAC-DIA data was searched with DIANN 2.2^34^, filtered for Q.value, Quality.Quantity, and channel.Q.value, and protein level fold changes were calculated based on empirical-Bayes modelling without imputation^35^. Cross-run normalization was activated for whole cell pSILAC-DIA in DIANN 2.2, and off for IP-MS experiment. Precursors were separated by channel and rolled up into proteins by summing the normalized intensities. For IP-MS, additional variance stabilizing and quantile normalization were applied. For gene enrichment analysis, ClusterProfiler (v3.18.1) was used to annotate Reactome pathways with total identified proteome from the experiment as background and compareCluster function. The STRINGDB annotations (v12.0, confidence threshold = 400) were incorporated using STRINGdb package (version 3.22) in R. The raw files and processed psm.txt files and report.parquet files are deposited on PRIDE under identification PXD087571 and PXD086047. R code for data processing is deposited on zenodo under DOI: 10.5281/zenodo.23885440

### Statistical analysis and machine learning

Z statistics: For each protein, log_2_FCs across all datasets (per SILAC channel) were used to calculate robust z statistics using Robustbase package (version 0.99-7) in R (version 4.5) as: 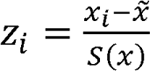 where *x~* is the median of protein log_2_FC across all KDs, x_i_ is the log_2_FC of knockdown *i* and S(x) is a robust scale estimator. Four alternative scale estimators were used namely Qn, Sn, Tau and Median Absolute Deviation (MAD)^36^. Significant outliers were filtered for z_i_ = >±2.5, log_2_FC >±0.58 and q value < 0.05. Machine learning (ML): SILAC pairs with fewer than 9 measurements were excluded from ML analysis. For each protein, log_2_FCs across all datasets and both channels were used for machine learning-based outlier detection based on MM-estimates, Minimum covariance determinant (MCD) and IsolationForest. For MM-estimation^37^, a regression model was fitted using lmrob function with an S-estimator used for initialization from Robustbase package (version 0.99-7). Each protein log_2_FC (x) across knockdowns (j) was fit as *x_Heavy,j_ = β_0_ + β_1_x_Light,j_ + ê_j_* with β_0_ and β_1_ as MM-regression intercept and slope respectively, and ê as residual. Iteratively, weighted least squares (*W_j_*) were estimated from the fitted model as *w_j_ = w(ê_j_/σ_s_c* and down-weighted proteins (< 0.1) were considered as outlier. The parameters for convergence stability, tuning constant (c) and estimation precision can be found in the R script. IsolationForest^38,39^ was applied through isotree package (version 0.6.1-5) with default settings on light and heavy log_2_FC two-dimensional data matrix to identify multivariate outliers. Outlier score was calculated as *s_j_ = 2^-E[h(x,j)]/c(n)^* where *E[h(x_j_)]* is the average path length required to isolate *x_j_* across randomly partitioned forest and c(n) as normalization constant. Proteins above *s_j_* > 0.7 were considered outliers. Lastly, robust mahalonobis distance 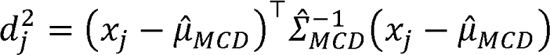 was computed based on location (μ^) and covariance estimates (Σ^) obtained by minimum covariance determinant (MCD) ^40,41^ using covMcd function in Robustbase with default settings. Observations with distance in top 0.0001^th^ percentile of the chi-squared distribution with 2 degrees of freedom were classified as outliers. Eventually, the outliers were further filtered for log_2_FC >±0.58 and q value < 0.05. The code and additional details on parameters are available on zenodo DOI: 10.5281/zenodo.23885440

### Protein turnover kinetics analysis

Protein turnover rates were determined from dynamic TMT-pulseSILAC experiment by modeling the fractional labeling ratio over time in a similar fashion as described by Zecha et al^42^. First, the TMT channels were normalized for total intensity and psms rolled up to peptides by proteotypicity and intensity summation. Only peptides with measurement in both light and heavy SILAC form were then selected for protein-level roll up with total intensity summation. For each protein, heavy (H) and light (L) channel TMT intensities were averaged across replicates (n = 2) at nine time points (0, 1, 2, 4, 6, 24, 48, 72, and 96 h) and arranged into protein-by-time-point matrices. The heavy-label fraction at each time point (*t*) was calculated as a ratio (*R*) of heavy to total.

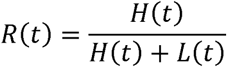

Proteins for which the heavy-label fraction did not increase between the first and last time point were excluded, as this is inconsistent with progressive isotope incorporation. For each remaining protein, the time-dependent (t) heavy-label fraction was fit to a single-exponential kinetic model that accounts for both protein degradation (k_deg_) and cell population growth (k_growth_) with doubling time of 32h.

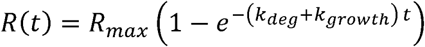

Nonlinear least-squares fitting was performed using the Levenberg-Marquardt algorithm (nlsLM, R package minpack.lm). Proteins for which the model failed to converge were excluded from downstream analysis. The code and additional details on parameters are available on zenodo DOI: 10.5281/zenodo.23885440

### Transcriptome sample preparation and sequencing

Cells in 24-well plates were lysed by freeze-thaw and RNA was extracted using NucleoSpin RNA isolation kit as per the manufacturer’s protocol. The quantity and quality of the RNA samples were assessed using Agilent 5400. Messenger RNA was purified from total RNA using poly-T oligo-attached magnetic beads. After fragmentation, the first strand cDNA was synthesized using random hexamer primers. Then the second strand cDNA was synthesized using dUTP, instead of dTTP. The directional library was ready after end repair, A-tailing, adapter ligation, size selection, USER enzyme digestion, amplification, and purification. The library was checked with Qubit and real-time PCR for quantification and bioanalyzer for size distribution detection. The qualified libraries were sequenced using Next Generation Sequencing (NGS) based on Illumina’s Sequencing Technology by Synthesis (SBS)-detection by fluorescence of the nucleotide added during the synthesis of the complementary chain - and in a parallelized and massive way. The Novaseq X Plus sequencing system was used to sequence the libraries. The strategy is paired-end 150bp (PE150).

### Transcriptome data analysis

Raw sequencing reads (paired-end) underwent quality control with FastQC (v0.12.1; Andrews, 2010) both before and after adapter trimming. Adapter and quality trimming was performed using Flexbar (v3.5.0) with paired-end mode and a Nextera adapter preset^43^. Trimmed reads were aligned to the human reference genome (GRCh38) using STAR (v2.7.11b) in paired-end mode with default parameters and BAM output sorted by coordinate^44^). Resulting BAM files were indexed using SAMtools (v1.19.2). Gene-level read counts were generated in R (v4.5.1) using the featureCounts function from the Rsubread package (doi: 10.1093/bioinformatics/btt656), using the Ensembl GRCh38.115 annotation (GFF3 format) as reference, with paired-end mode enabled and reads summarized at the gene feature level (GTF.featureType = “gene”, GTF.attrType = “ID”). Differential gene expression analysis was performed in R (v4.5.1) using DESeq2 (doi: 10.1186/s13059-014-0550-8). Genes with a total read count below 10 across all samples were excluded prior to model fitting. A DESeqDataSet was constructed from the raw count matrix with a single-factor design (∼ group), and the model was fit using the standard DESeq2 workflow (median-of-ratios normalization, negative binomial generalized linear model, and Wald testing). For each comparison, log_2_ fold changes were shrunk using the apeglm method (doi: 10.1093/bioinformatics/bty895) to improve effect-size estimation for genes with low counts or high dispersion. Genes were considered significantly differentially expressed at an adjusted p-value (Benjamini–Hochberg) < 0.05 and an absolute shrunken log_2_ fold change ≥ 0.58.

### mitoKeima assay

HeLa FlpIn mt-mKEIMA cells with inducible PRKN were pre-treated with 0.25 µg/ml doxycycline for 24h prior to flow cytometric measurement. In brief, after treatment with 1.3 or 2 µM Antimycin A (Sigma, A8674) /Oligomycin (MedChemExpress, HY-16589) or DMSO for 4 h, the cells were washed 3x with prewarmed HBSS. Flow-cytometry analysis for mKEIMA was performed on a BD FACSymphony A5 flow cytometry system (BD). Analysis of flow cytometer data was performed with FlowJo v.10. Viable and single cell populations were gated for dual excitation at 405 (pH 7) and 561 (pH 4) nm as described previously^45^. The percentage of lysosomal mt-mKEIMA was calculated using the ratio of mKEIMA excitation at 561 nm/405 nm.

### Western blot and PhosTAG gel

Cells were transfected with siRNA in 6-well as described above. Transfected cells were lysed in RIPA buffer containing protease inhibitor cocktail, PhosStop (Roche, 4906837001), and Benzonase (Santa Cruz Biotechnology, sc-202391). Cell debris was removed by centrifugation at 12.500 g for 15 min at 4°C. Protein concentration of the supernatant was quantified using BCA Assay (Thermo Fisher Scientific, 23225). Equal protein lysate was prepared in 1x reducing Laemmli buffer and boiled for 10 min at 95 °C. Proteins were separated by SDS-PAGE using 4–12% Bolt Bis-Tris Plus Gels (Thermo Fisher Scientific, NW04120BOX) or using SuperSep™ Phos-tag™ Precast Gels (Fujifilm, 199-18011) according to the manufacturer’s instructions. Proteins were transferred onto 0.2 µm nitrocellulose membrane (Bio-Rad, 1704270) using the Trans-Blot Turbo Transfer System (Bio-Rad). Phos-tag™ Precast Gels were washed for 10 min in running buffer with 10 mM EDTA and two times for 10 min in EDTA-free running buffer before membrane transfer. Immunoblots were blocked with Intercept Blocking Buffer (LI-COR Biosciences, 927-70001) overnight at 4°C. Primary and secondary antibodies were added to immunoblots for 1 h at RT. Primary and secondary antibodies used for the detection were α-MCL-1 (CST, 94296), α-Tubulin (CST, 3873), IRDye 800CW goat anti-rabbit IgG (H + L; Li-Cor catalogue no. 926–32211, 1:15,000), and IRDye 680RD donkey anti-mouse IgG (H + L; Li-Cor catalogue no. 926–68072, 1:15,000). Immunoblot analysis was performed with Image Studio v.6.0 (LI-COR).

### Co-immunoprecipitation

Cells transfected and pulsed as outlined above were lysed in IP lysis buffer (20 mM TRIS pH 8, 137 mM NaCl, 1% NP40, 2 mM EDTA, protease inhibitor cocktail) and incubated for 30 min at 4°C with overhead rotation. Cell debris was removed by centrifugation at 13.500 g for 20 min at 4°C and protein amount in the supernatant was determined using BCA assay. MCL-1 was immunoprecipitated with 0.255 µg α-MCL-1 antibody (CST, 94296) coupled to Pierce™ Protein A Magnetic Beads (Thermo Fischer Scientific, 88846) overnight at 4°C. After two washes with lysis buffer, three washes with PBS and a final wash with 50 mM Tris pH 7.8, bound proteins were eluted in two steps by incubating beads with 2% SDC in 50 mM Tris pH 7.8 at 95°C for 5 min. Eluted proteins were reduced and acetylated with 1 mM TCEP, 2 mM CAA at 95 °C for 5 min and digested overnight at 37°C with 500 ng trypsin and Lys C each. Digestion was stopped with isopropanol, and samples were acidified with TFA. Sample was desalted using SDB-RPS (3M Empore). Samples were dried using a vacuum concentrator and resuspended in 2% ACN, 0.1% TFA for LC-MS analysis.

