## Supplementary File 2 for "A dynamic proteomic HECT E3 ligase screen reveals substrate exclusivity and redundancies"

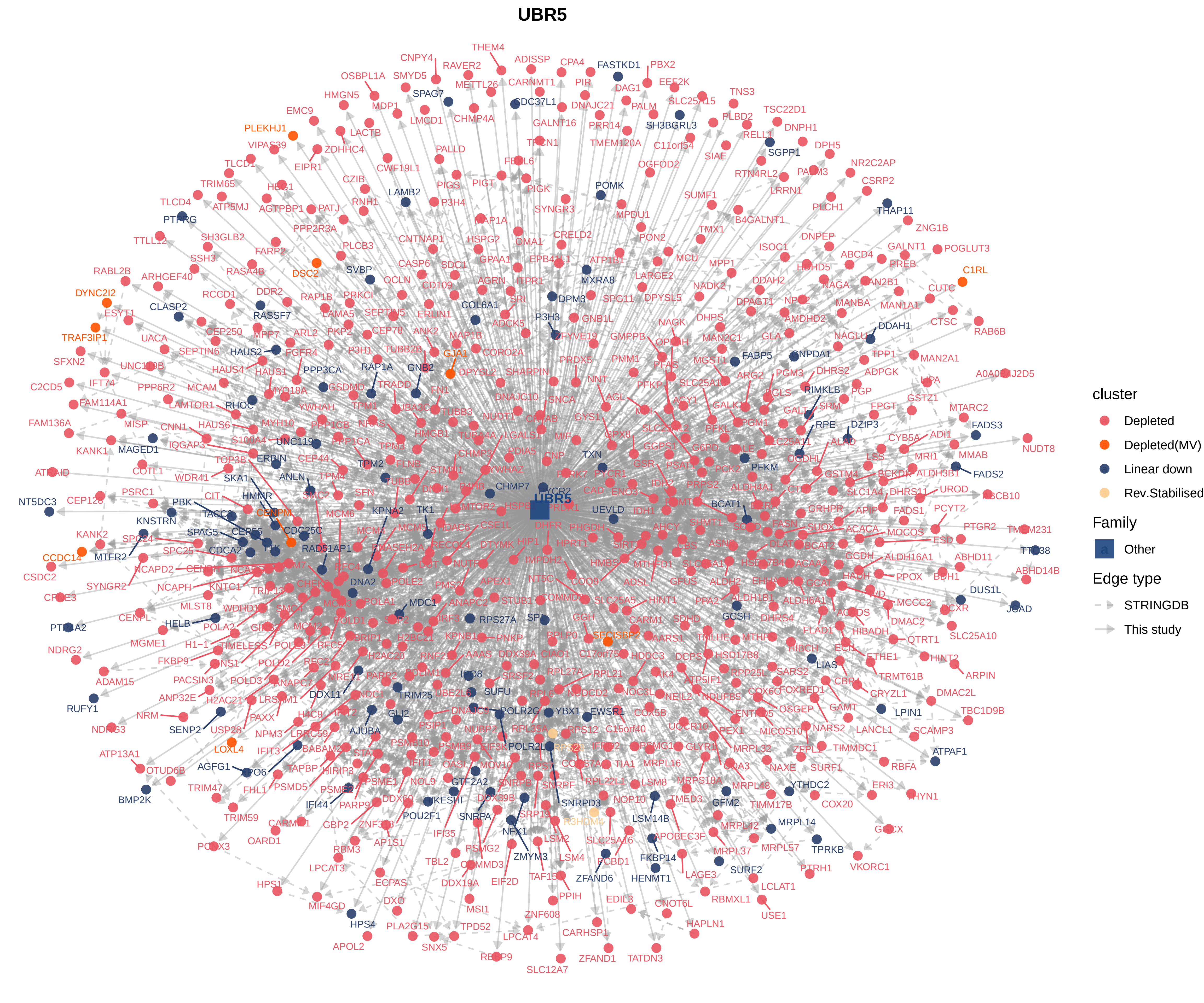
UBR5
cluster
Depleted
Depleted(MV)
Linear down
Rev.Stabilised
Family
Other
Edge type
STRINGDB
This study

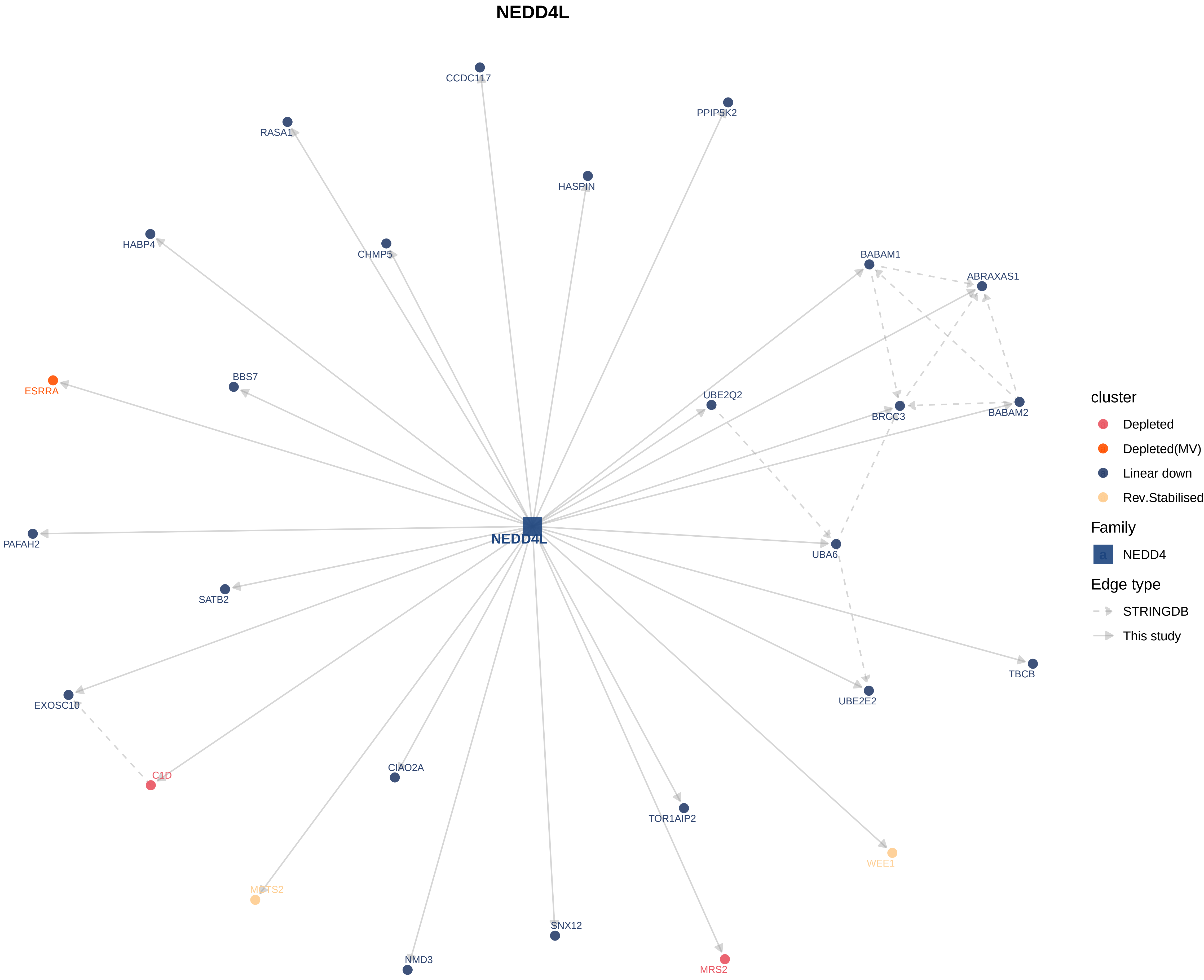
NEDD4L
CCDC117
PPIP5K2
RASA1
HASPIN
HABP4
CHMP5
BABAM1
ABRAXAS1
ESRRA
BBS7
UBE2Q2
BRCC3
BABAM2
PAFAH2
NEDD4L
UBA6
SATB2
TBCB
UBE2E2
EXOSC10
C1D
CIAO2A
TOR1AIP2
WEE1
SNX12
NMD3
MRS2
cluster
Depleted
Depleted(MV)
Linear down
Rev.Stabilised
Family
NEDD4
Edge type
STRINGDB
This study

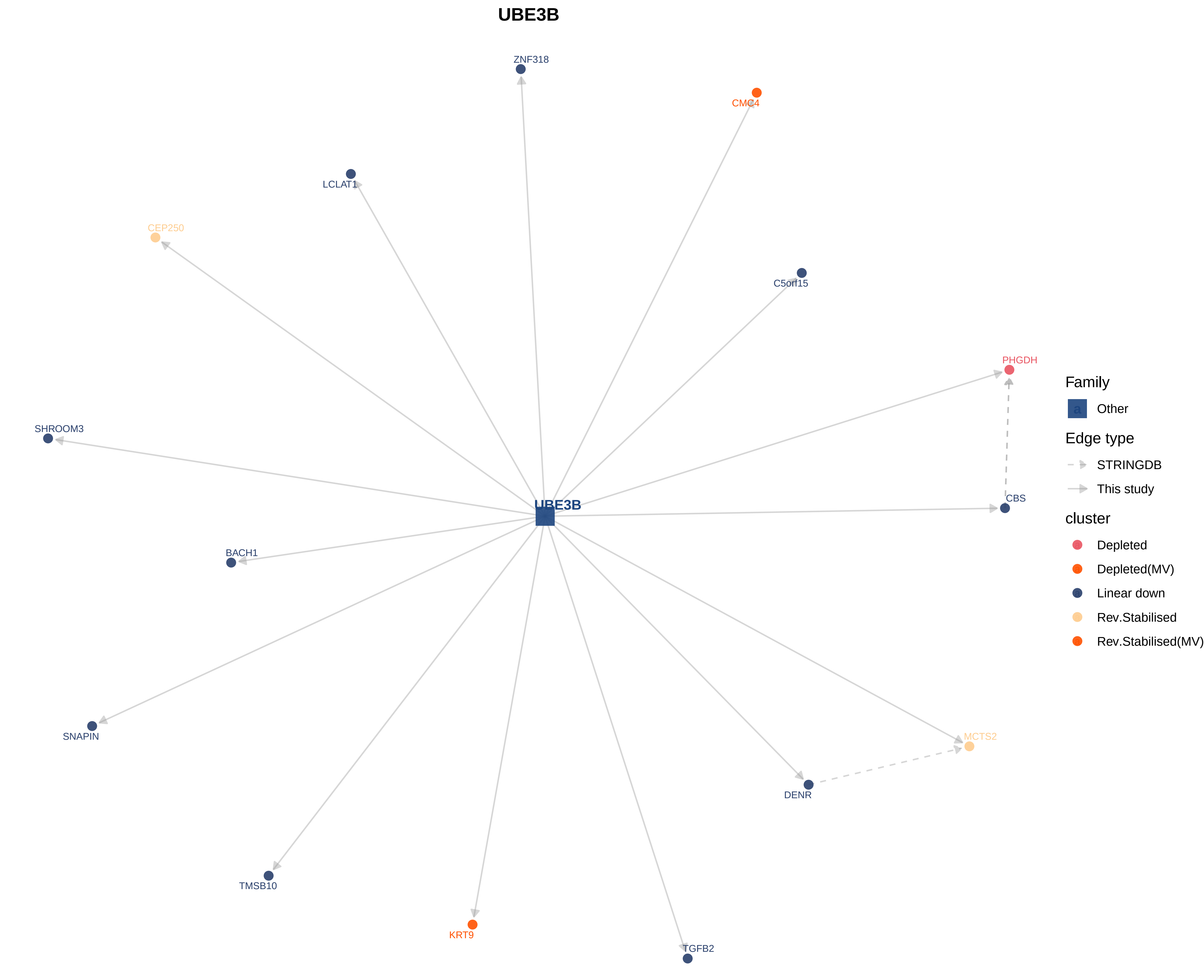
UBE3B
ZNF318
CMC4
LCLAT1
CEP250
C5orf15
PHGDH
SHROOM3
CBS
UBE3B
BACH1
SNAPIN
MCTS2
DENR
TMSB10
KRT9
TGFB2
Family
Other
Edge type
STRINGDB
This study
cluster
Depleted
Depleted(MV)
Linear down
Rev.Stabilised
Rev.Stabilised(MV)

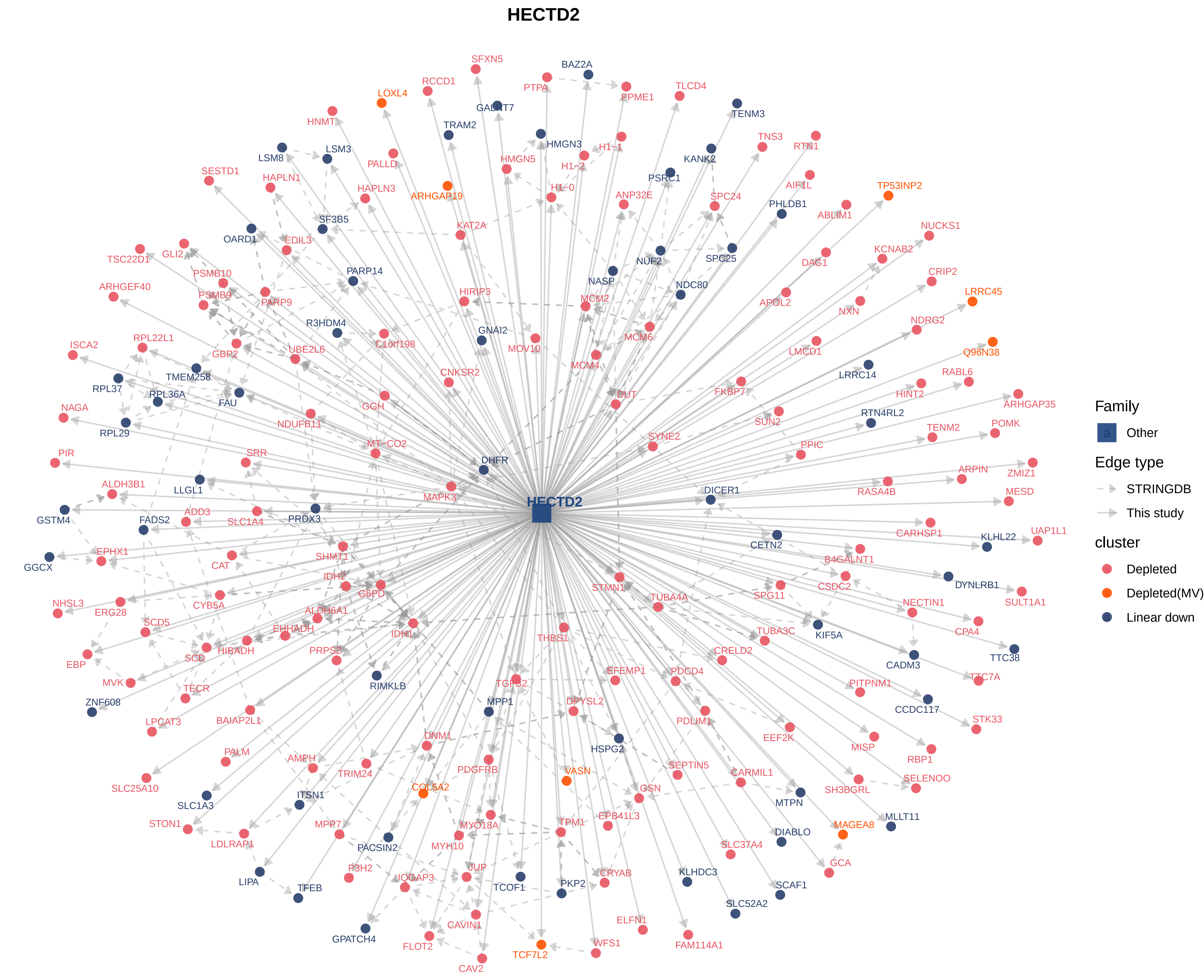

HECTD2
SFXN5
BAZ2A
RCCD1
PTPA
PPME1
TLCD4
LOXL4
GALNT7
TENM3
HNMT
TRAM2
HMGN3
H1-1
TNS3
RTN1
LSM8
LSM3
PALLD
HMGN5
H1-2
KANK2
SESTD1
HAPLN1
PSRC1
AIF1L
TP53INP2
HAPLN3
ARHGAP19
H1-0
ANP32E
SPC24
PHLDB1
ABLIM1
SF3B5
OARD1
KAT2A
EDIL3
NUCKS1
TSC22D1
GLI2
NUF2
SPC25
KCNAB2
DAG1
PSMB10
PARP14
NASP
CRIP2
ARHGEF40
NDC80
PSMB9
PARP9
HIRIP3
MCM2
LRRC45
APOL2
NXN
R3HDM4
GNAI2
NDRG2
ISCA2
RPL22L1
C1orf198
MOV10
MCM6
GBP2
UBE2L6
LMCD1
Q96N38
TMEM258
MCM4
LRRC14
RPL37
CNKSR2
RABL6
RPL36A
FAU
HINT2
DUT
FKBP7
NAGA
GGH
ARHGAP35
NDUFB11
RTN4RL2
RPL29
SUN2
TENM2
POMK
MT-CO2
SYNE2
PPIC
PIR
SRR
DHFR
ARPIN
ZMIZ1
ALDH3B1
LLGL1
MAPK3
RASA4B
DICER1
MESD
HECTD2
GSTM4
ADD3
PRDX3
FADS2
SLC1A4
CARHSP1
KLHL22
UAP1L1
CETN2
EPHX1
SHMT1
GGCX
CAT
B4GALNT1
IDH2
DYNLRB1
STMN1
CSDC2
SPG11
G6PD
TUBA4A
SULT1A1
NHSL3
CYB5A
NECTIN1
ERG28
ALDH6A1
KIF5A
CPA4
SCD5
EHHADH
IDH1
TUBA3C
THBS1
CADM3
TTC38
EBP
SCD
HIBADH
PRPS2
CRELD2
MVK
EFEMP1
PDCD4
TTC7A
RIMKLB
PITPNM1
TECR
TGFB2
CCDC117
ZNF608
MPP1
DPYSL2
STK33
LPCAT3
BAIAP2L1
PDLIM1
EEF2K
DNM1
HSPG2
MISP
RBP1
PALM
AMPH
TRIM24
PDGFRB
SEPTIN5
CARMIL1
SELENOO
SLC25A10
VASN
SH3BGRL
COL5A2
GSN
MTPN
SLC1A3
ITSN1
MLLT11
STON1
MPP7
MYO18A
TPM1
EPB41L3
MAGEA8
LDLRAP1
PACSIN2
MYH10
DIABLO
SLC37A4
GCA
P3H2
JUP
CRYAB
KLHDC3
LIPA
IQGAP3
TCOF1
PKP2
SCAF1
TFEB
SLC52A2
ELFN1
CAVIN1
GPATCH4
FLOT2
FAM114A1
WFS1
TCF7L2
CAV2
Family
Other
Edge type
STRINGDB
This study
cluster
Depleted
Depleted(MV)
Linear down

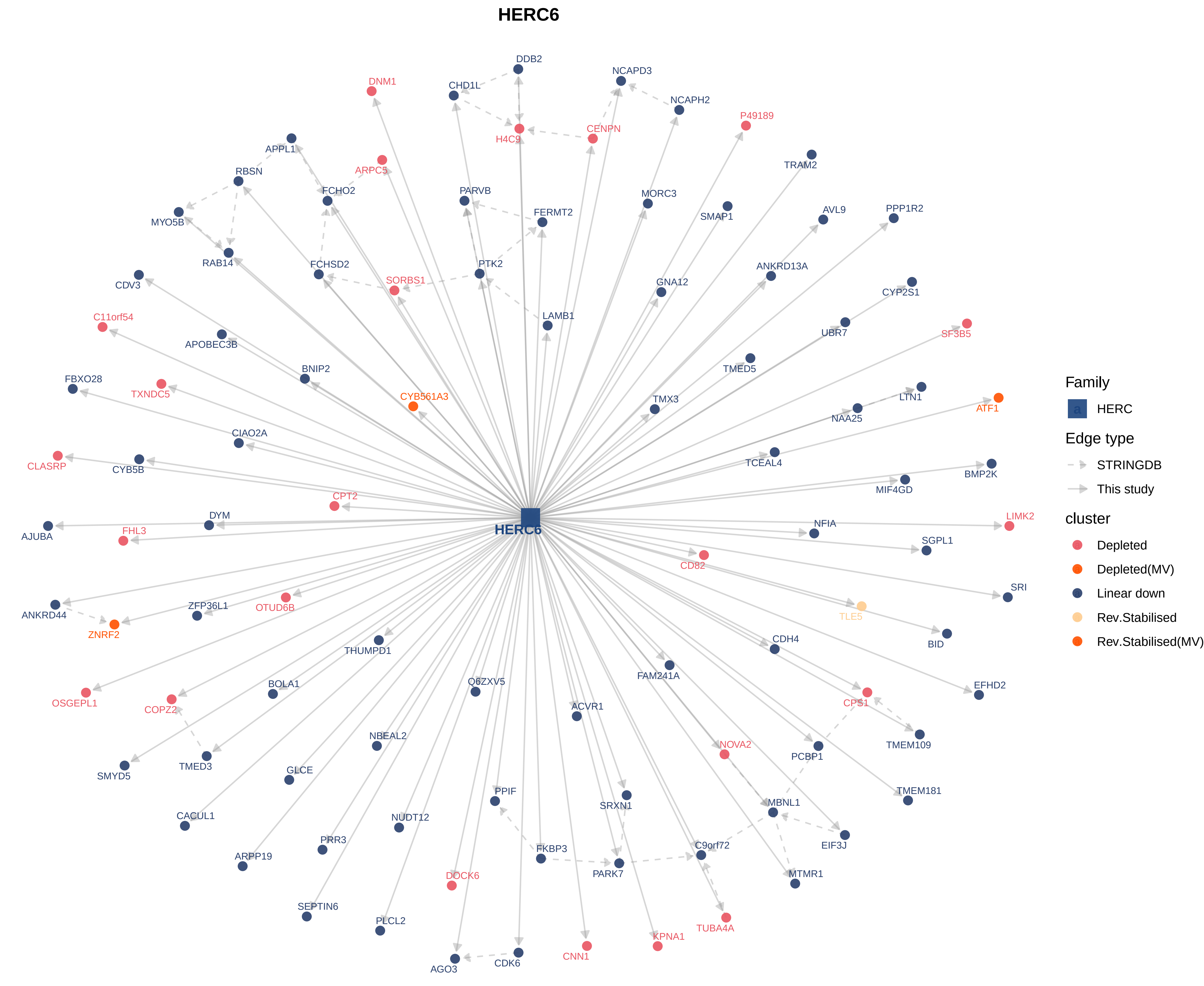
HERC6
DDB2
NCAPD3
DNM1
CHD1L
NCAPH2
P49189
CENPN
H4C9
APPL1
ARPC5
TRAM2
RBSN
FCHO2
PARVB
MORC3
SMAP1
AVL9
PPP1R2
MYO5B
FERMT2
RAB14
FCHSD2
PTK2
ANKRD13A
SORBS1
GNA12
CDV3
CYP2S1
LAMB1
C11orf54
UBR7
SF3B5
APOBEC3B
BNIP2
TMED5
FBXO28
TXNDC5
CYB561A3
TMX3
LTN1
ATF1
NAA25
CIAO2A
CLASRP
CYB5B
TCEAL4
BMP2K
MIF4GD
CPT2
DYM
AJUBA
FHL3
HERC6
NFIA
LIMK2
SGPL1
CD82
SRI
ANKRD44
ZFP36L1
OTUD6B
TLE5
ZNRF2
BID
CDH4
THUMPD1
FAM241A
EFHD2
OSGEPL1
BOLA1
Q6ZXV5
CPS1
COPZ2
ACVR1
TMEM109
NBEAL2
NOVA2
PCBP1
SMYD5
TMED3
GLCE
PPIF
SRXN1
MBNL1
TMEM181
CACUL1
NUDT12
EIF3J
PRR3
FKBP3
C9orf72
ARPP19
PARK7
MTMR1
DOCK6
SEPTIN6
TUBA4A
PLCL2
KPNA1
CNN1
AGO3
CDK6
Family
HERC
Edge type
STRINGDB
This study
cluster
Depleted
Depleted(MV)
Linear down
Rev.Stabilised
Rev.Stabilised(MV)

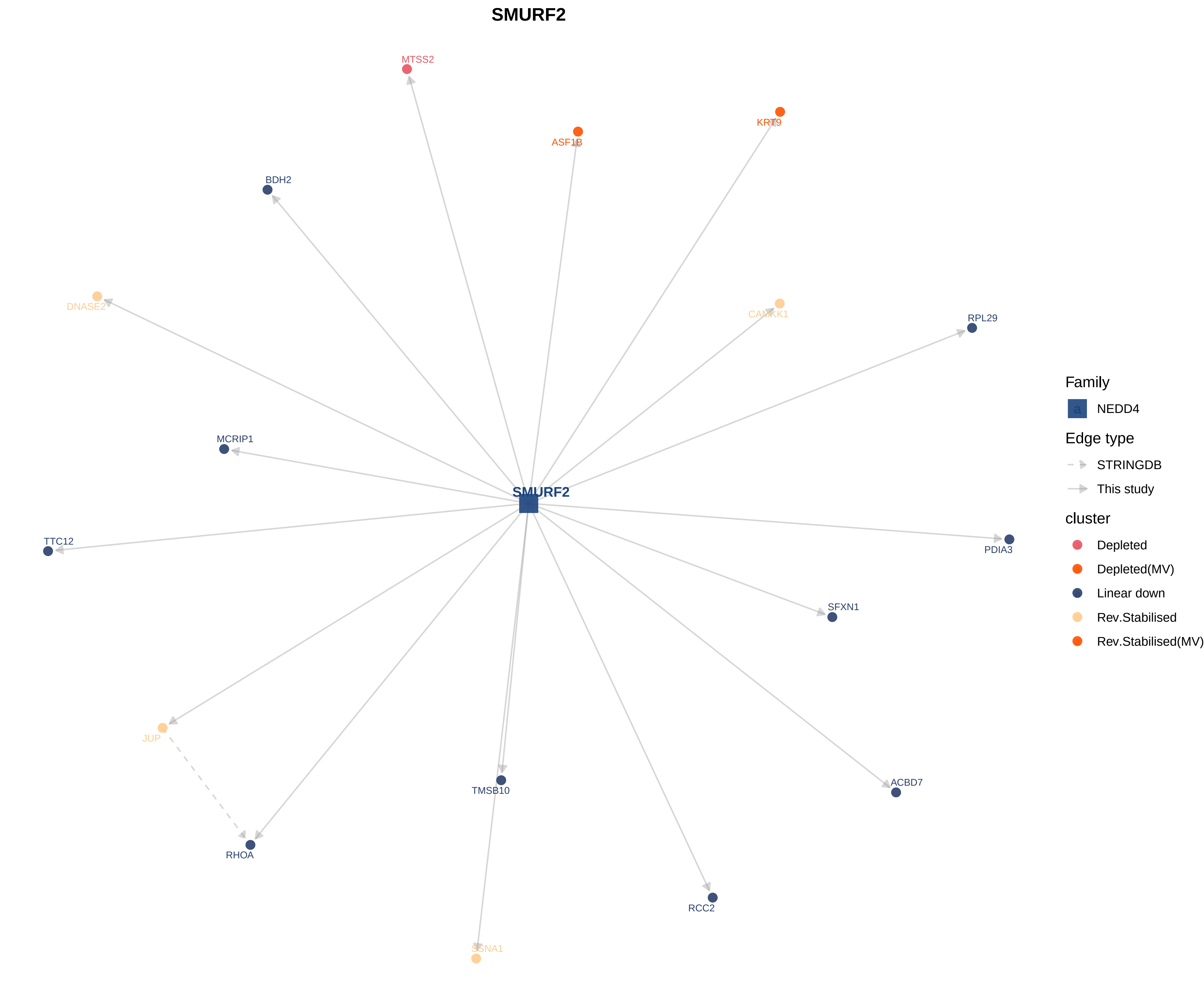
SMURF2
MTSS2
KRT9
ASF1B
BDH2
DNASE2
CAMKK1
RPL29
MCRIP1
SMURF2
TTC12
PDIA3
SFXN1
JUP
TMSB10
ACBD7
RHOA
RCC2
SSNA1
Family
NEDD4
Edge type
STRINGDB
This study
cluster
Depleted
Depleted(MV)
Linear down
Rev.Stabilised
Rev.Stabilised(MV)

### HACE1

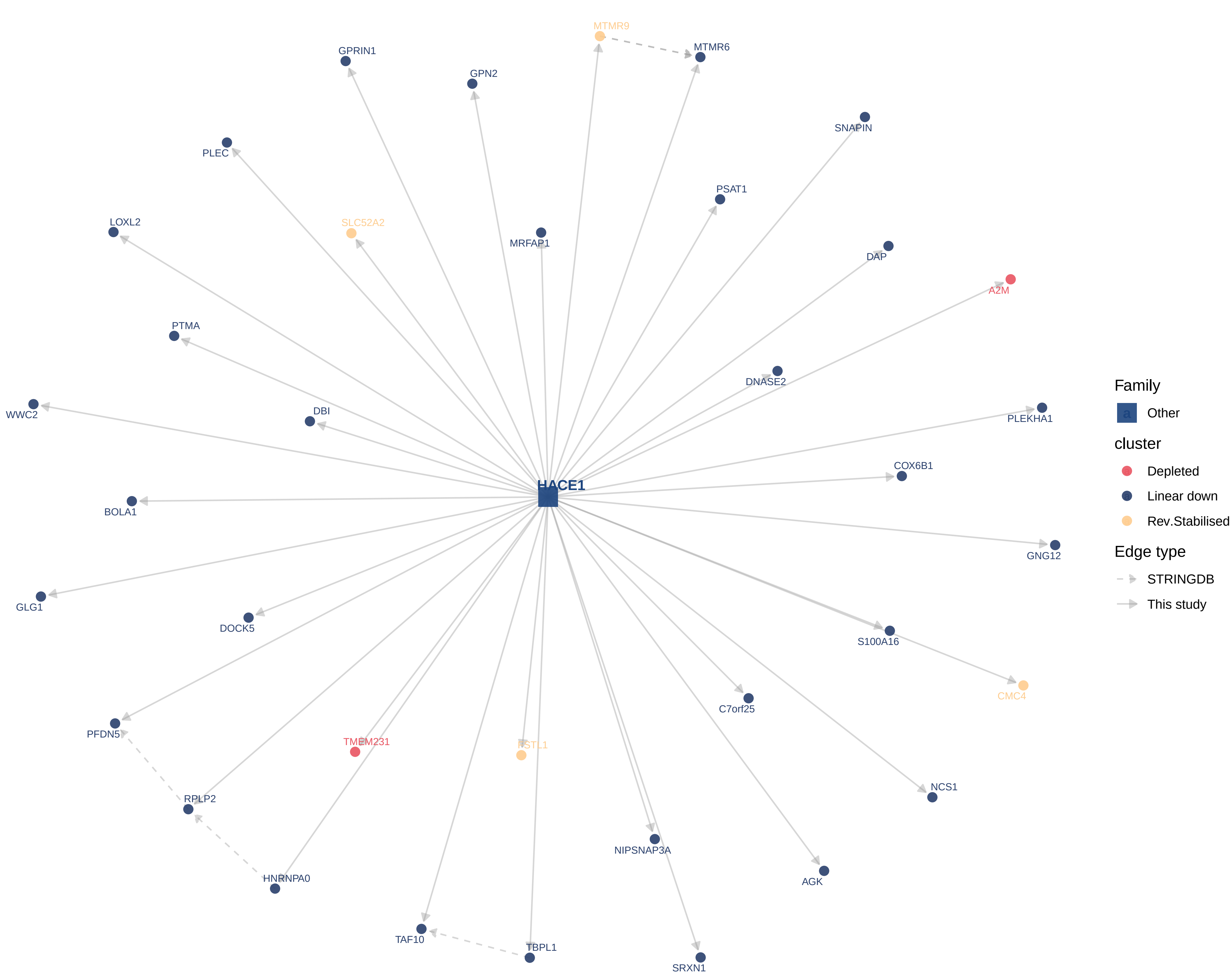

HACE1
MTMR9
MTMR6
GPRIN1
GPN2
SNAPIN
PLEC
PSAT1
LOXL2
SLC52A2
MRFAP1
DAP
A2M
PTMA
DNASE2
WWC2
DBI
PLEKHA1
COX6B1
BOLA1
GNG12
GLG1
DOCK5
S100A16
CMC4
C7orf25
PFDN5
TMEM231
FSTL1
NCS1
RPLP2
NIPSNAP3A
HNRNPA0
AGK
TAF10
TBPL1
SRXN1
Family
Other
cluster
Depleted
Linear down
Rev.Stabilised
Edge type
STRINGDB
This study

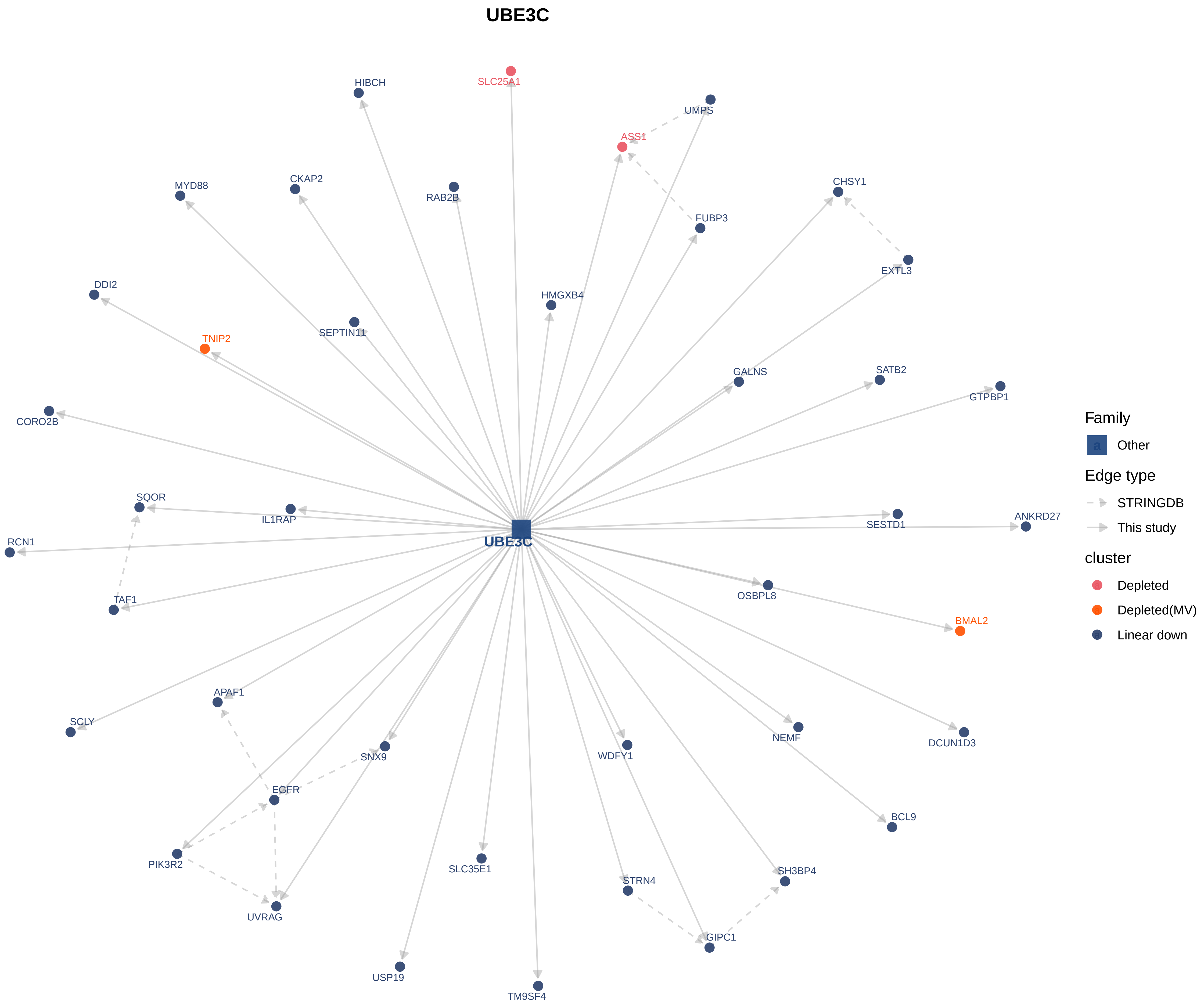

UBE3C
SLC25A1
HIBCH
UMPS
ASS1
MYD88
CKAP2
RAB2B
CHSY1
FUBP3
EXTL3
DDI2
HMGXB4
TNIP2
SEPTIN11
GALNS
SATB2
GTPBP1
CORO2B
Family
Other
Edge type
STRINGDB
This study
SQOR
IL1RAP
SESTD1
ANKRD27
RCN1
UBE3C
cluster
Depleted
Depleted(MV)
Linear down
OSBPL8
TAF1
BMAL2
APAF1
SCLY
NEMF
DCUN1D3
SNX9
WDFY1
EGFR
BCL9
PIK3R2
SLC35E1
SH3BP4
STRN4
UVRAG
GIPC1
USP19
TM9SF4

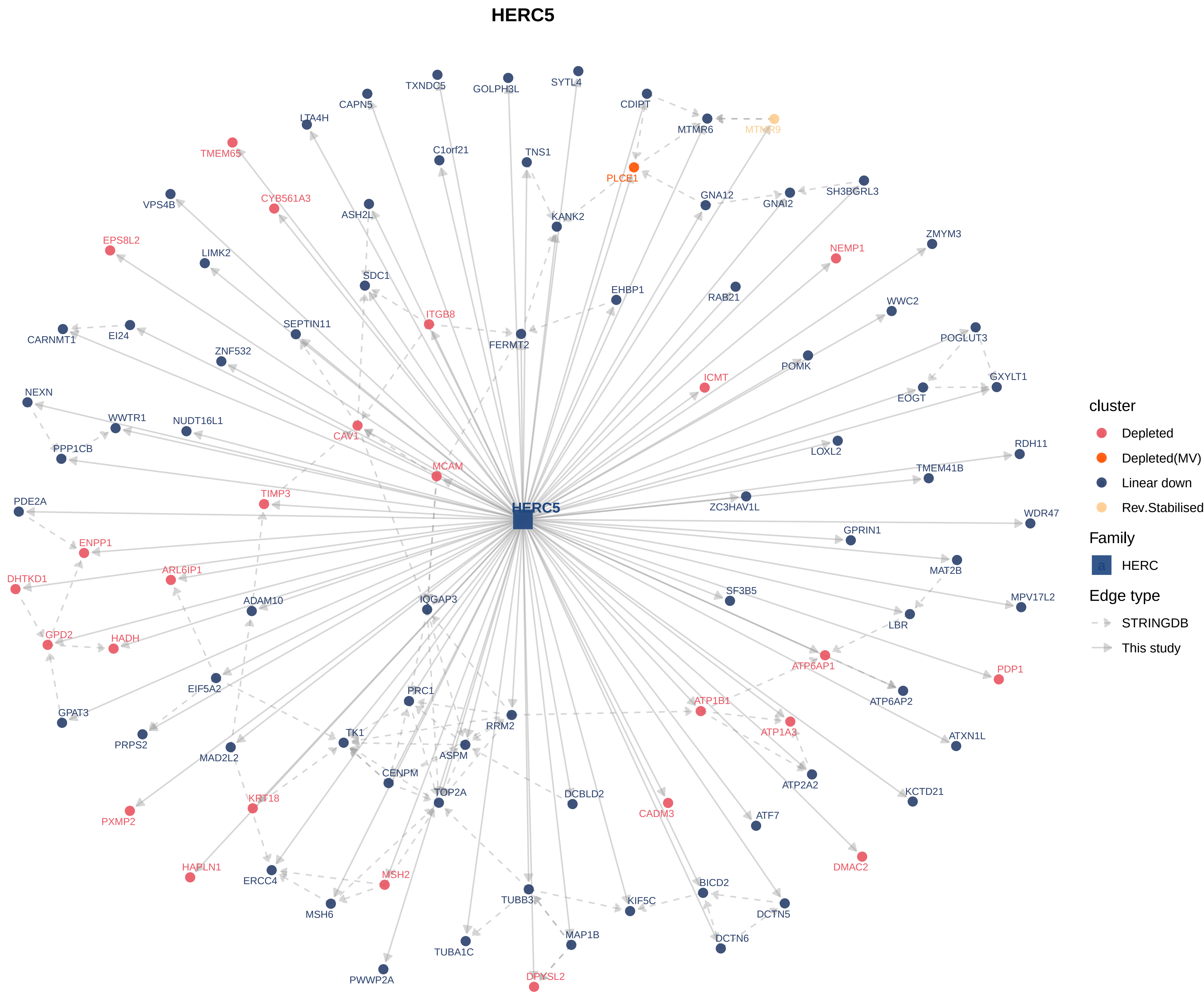
HERC5
TXNDC5
GOLPH3L
SYTL4
CAPN5
CDIPT
LTA4H
MTMR6
MTMR9
TMEM65
C1orf21
TNS1
PLCE1
VPS4B
CYB561A3
ASH2L
GNA12
GNAI2
SH3BGRL3
KANK2
EPS8L2
LIMK2
NEMP1
ZMYM3
SDC1
EHBP1
RAB21
WWC2
ITGB8
CARNMT1
EI24
SEPTIN11
FERMT2
POGLUT3
ZNF532
POMK
ICMT
GXYLT1
NEXN
EOGT
WWTR1
NUDT16L1
CAV1
PPP1CB
LOXL2
RDH11
MCAM
TMEM41B
TIMP3
PDE2A
HERC5
ZC3HAV1L
WDR47
GPRIN1
ENPP1
DHTKD1
ARL6IP1
MAT2B
ADAM10
IQGAP3
SF3B5
MPV17L2
LBR
GPD2
HADH
ATP6AP1
PDP1
EIF5A2
PRC1
ATP6AP2
GPAT3
ATP1B1
RRM2
ATP1A3
PRPS2
TK1
ATXN1L
MAD2L2
ASPM
CENPM
ATP2A2
KCTD21
KRT18
TOP2A
DCBLD2
CADM3
PXMP2
ATF7
HAPLN1
MSH2
DMAC2
ERCC4
BICD2
TUBB3
KIF5C
MSH6
DCTN5
MAP1B
DCTN6
TUBA1C
PWWP2A
DPYSL2
cluster
Depleted
Depleted(MV)
Linear down
Rev.Stabilised
Family
HERC
Edge type
STRINGDB
This study

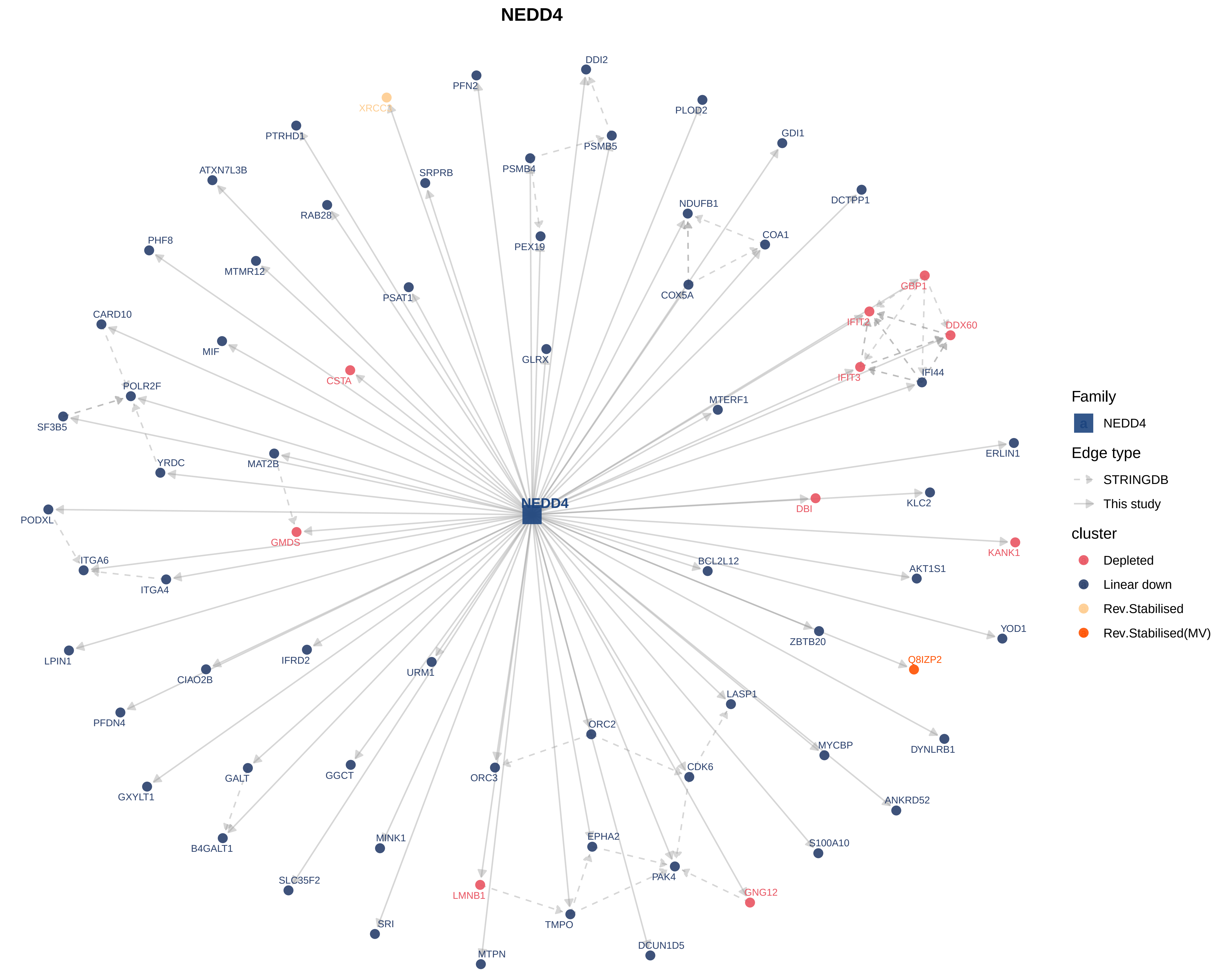

NEDD4
DDI2
PFN2
XRCC1
PLOD2
PTRHD1
PSMB5
GDI1
PSMB4
ATXN7L3B
SRPRB
NDUFB1
DCTPP1
RAB28
PEX19
COA1
PHF8
MTMR12
PSAT1
COX5A
GBP1
CARD10
IFIT2
DDX60
MIF
GLRX
CSTA
IFIT3
IFI44
POLR2F
MTERF1
SF3B5
YRDC
MAT2B
ERLIN1
NEDD4
DBI
KLC2
PODXL
GMDS
KANK1
ITGA6
BCL2L12
AKT1S1
ITGA4
YOD1
ZBTB20
LPIN1
IFRD2
Q8IZP2
CIAO2B
URM1
PFDN4
LASP1
ORC2
MYCBP
DYNLRB1
CDK6
GXYLT1
GALT
GGCT
ORC3
ANKRD52
B4GALT1
MINK1
EPHA2
S100A10
PAK4
SLC35F2
LMNB1
GNG12
TMPO
SRI
DCUN1D5
MTPN
Family
NEDD4
Edge type
STRINGDB
This study
cluster
Depleted
Linear down
Rev.Stabilised
Rev.Stabilised(MV)

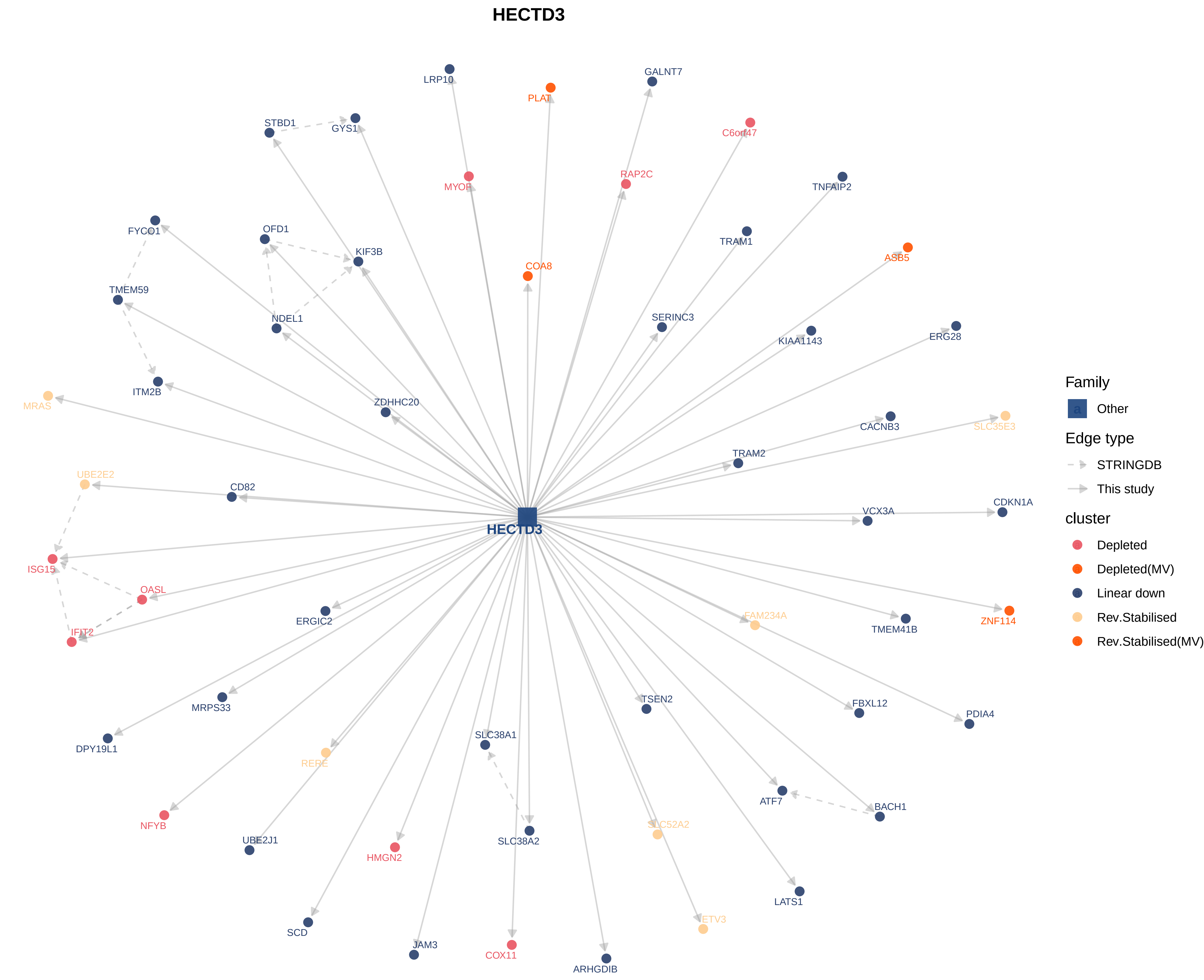

HECTD3
LRP10
PLAT
GALNT7
STBD1
GYS1
C6orf47
MYOF
RAP2C
TNFAIP2
FYCO1
OFD1
KIF3B
TRAM1
COA8
ASB5
TMEM59
NDEL1
SERINC3
KIAA1143
ERG28
ITM2B
MRAS
ZDHHC20
CACNB3
SLC35E3
TRAM2
UBE2E2
CD82
VCX3A
CDKN1A
ISG15
OASL
ERGIC2
FAM234A
TMEM41B
ZNF114
IFIT2
MRPS33
TSEN2
FBXL12
PDIA4
DPY19L1
SLC38A1
RERE
ATF7
BACH1
NFYB
UBE2J1
SLC38A2
SLC52A2
HMGN2
LATS1
ETV3
SCD
JAM3
COX11
ARHGDIB
Family
Other
Edge type
STRINGDB
This study
cluster
Depleted
Depleted(MV)
Linear down
Rev.Stabilised
Rev.Stabilised(MV)

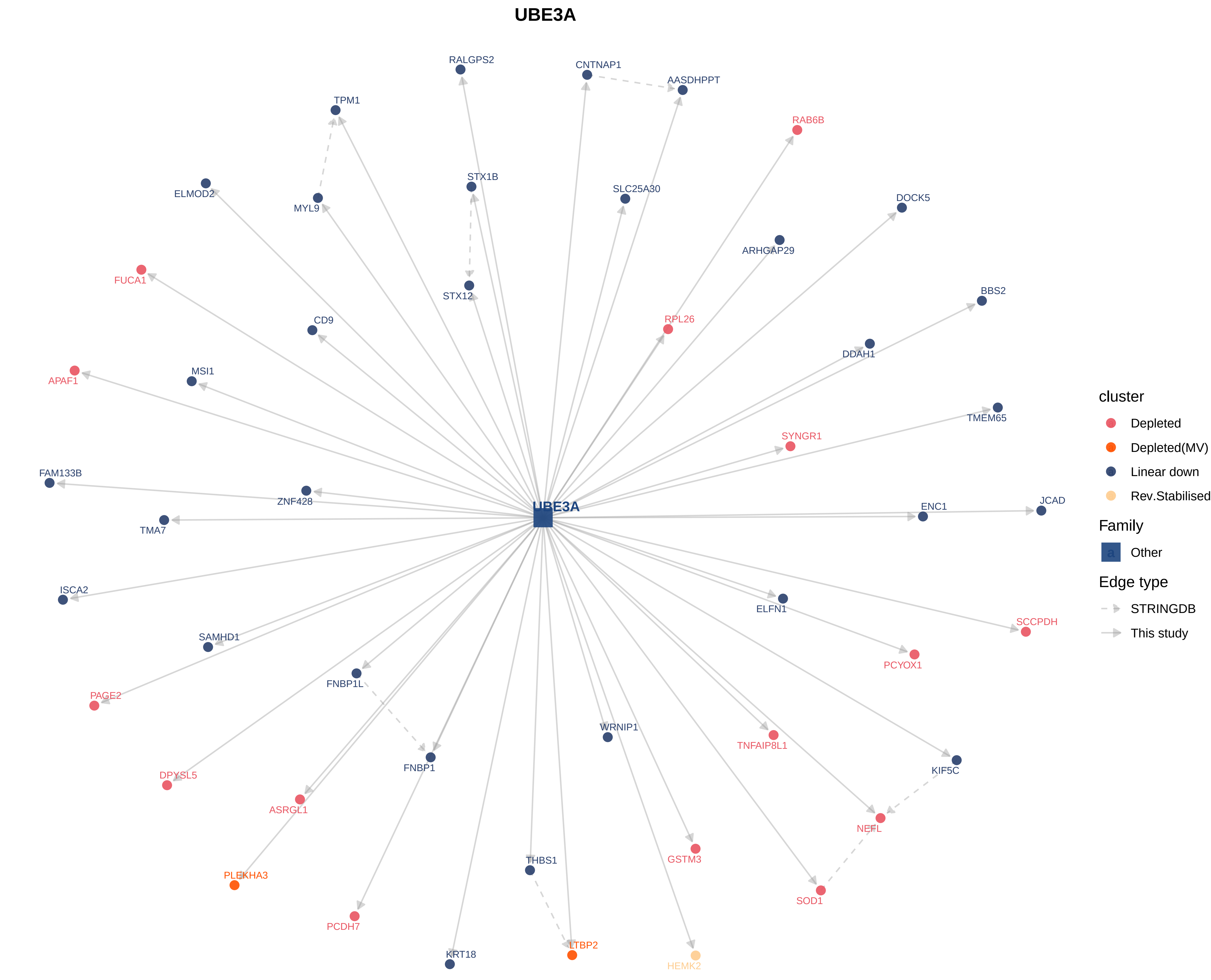

UBE3A
RALGPS2
CNTNAP1
AASDHPPT
TPM1
RAB6B
STX1B
SLC25A30
DOCK5
ELMOD2
MYL9
ARHGAP29
FUCA1
STX12
BBS2
CD9
RPL26
DDAH1
APAF1
MSI1
TMEM65
SYNGR1
FAM133B
ZNF428
TMA7
ENC1
JCAD
ISCA2
ELFN1
SCCPDH
SAMHD1
PCYOX1
PAGE2
FNBP1L
WRNIP1
TNFAIP8L1
KIF5C
DPYSL5
FNBP1
ASRGL1
NEFL
THBS1
GSTM3
PLEKHA3
SOD1
PCDH7
LTBP2
KRT18
HEMK2
cluster
Depleted
Depleted(MV)
Linear down
Rev.Stabilised
Family
Other
Edge type
STRINGDB
This study

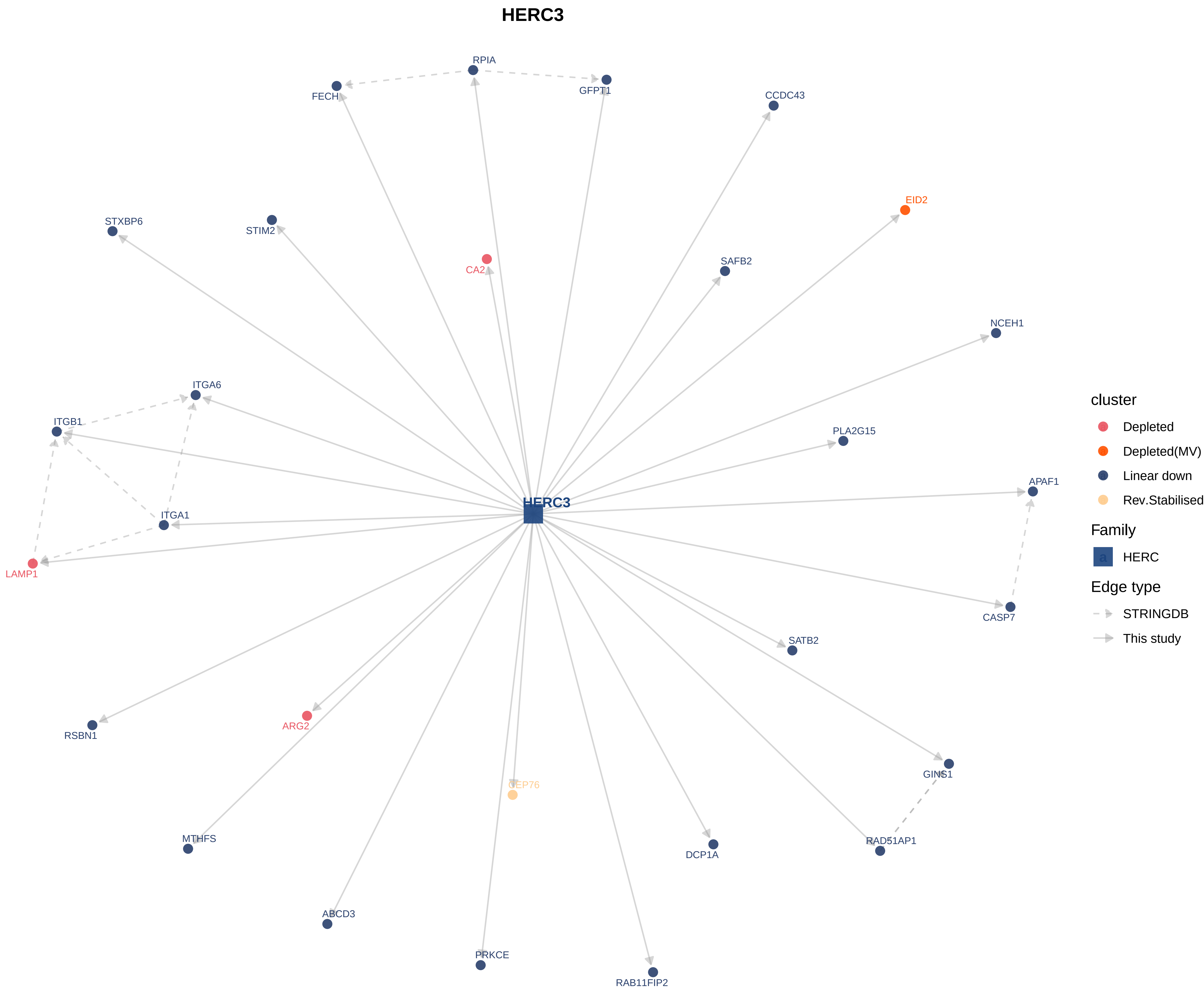

HERC3
RPIA
FECH
GFPT1
CCDC43
EID2
STXBP6
STIM2
CA2
SAFB2
NCEH1
ITGA6
ITGB1
PLA2G15
APAF1
HERC3
ITGA1
LAMP1
CASP7
SATB2
RSBN1
ARG2
GINS1
CEP76
MTHFS
RAD51AP1
DCP1A
ABCD3
PRKCE
RAB11FIP2
cluster
Depleted
Depleted(MV)
Linear down
Rev.Stabilised
Family
HERC
Edge type
STRINGDB
This study

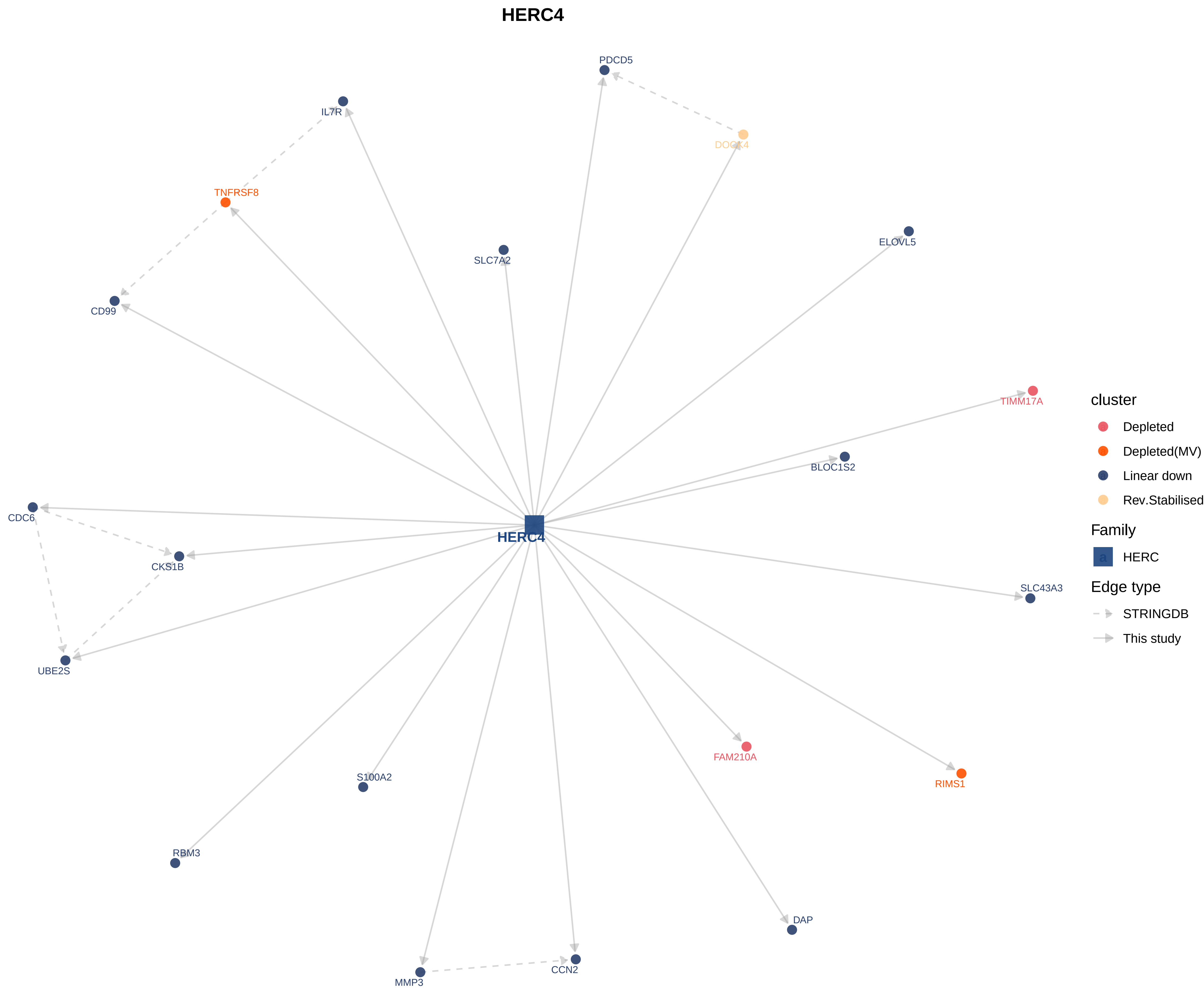
HERC4
PDCD5
IL7R
DOCK4
TNFRSF8
SLC7A2
ELOVL5
CD99
TIMM17A
BLOC1S2
CDC6
HERC4
CKS1B
SLC43A3
UBE2S
FAM210A
RIMS1
S100A2
RBM3
DAP
CCN2
MMP3
cluster
Depleted
Depleted(MV)
Linear down
Rev.Stabilised
Family
HERC
Edge type
STRINGDB
This study

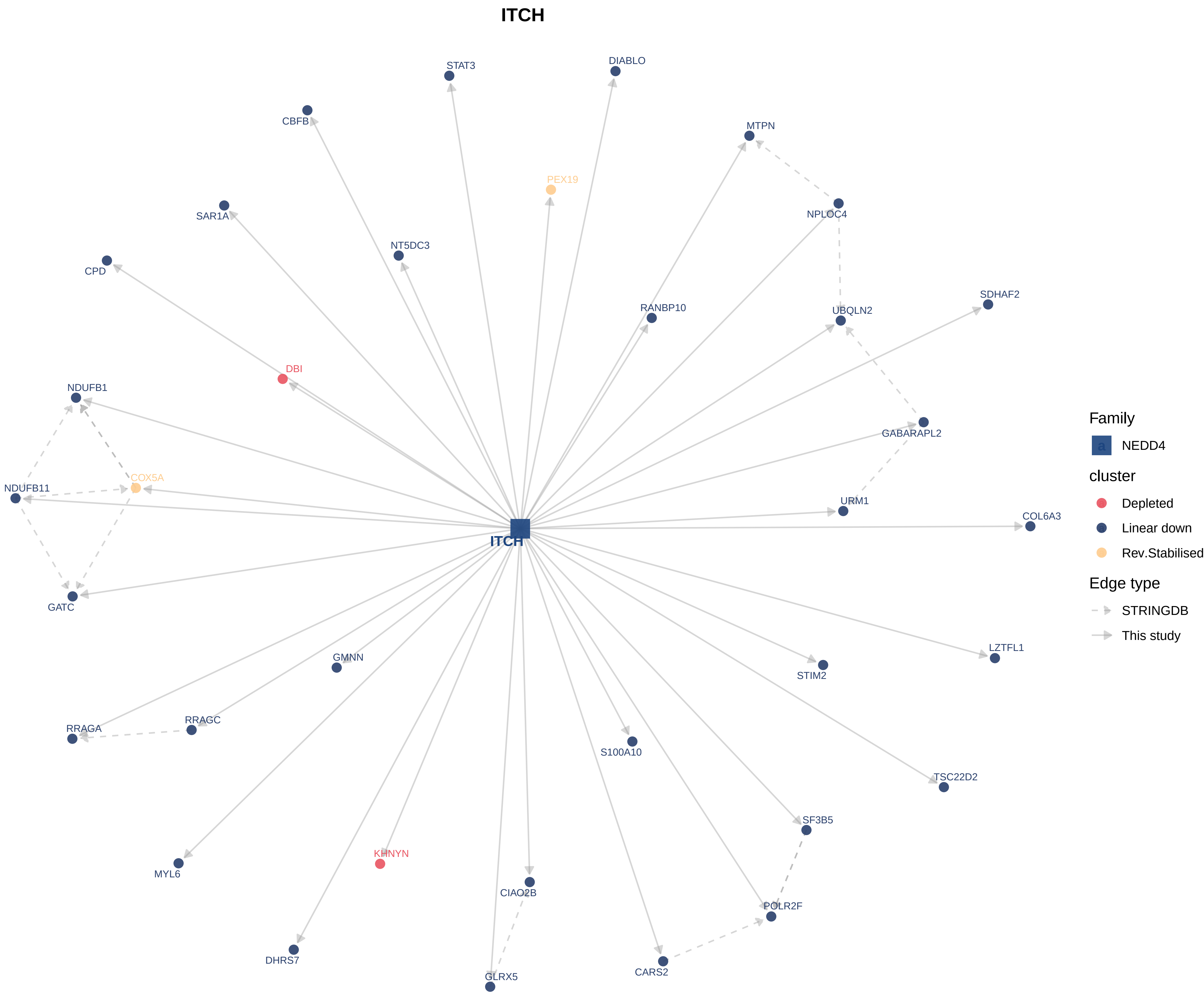

ITCH
STAT3
DIABLO
CBFB
MTPN
PEX19
SAR1A
NPLOC4
NT5DC3
CPD
RANBP10
UBQLN2
SDHAF2
DBI
NDUFB1
GABARAPL2
COX5A
NDUFB11
URM1
COL6A3
ITCH
GATC
GMNN
LZTFL1
STIM2
RRAGA
RRAGC
S100A10
TSC22D2
SF3B5
KHNYN
MYL6
CIAO2B
POLR2F
DHRS7
CARS2
GLRX5
Family
NEDD4
cluster
Depleted
Linear down
Rev.Stabilised
Edge type
STRINGDB
This study

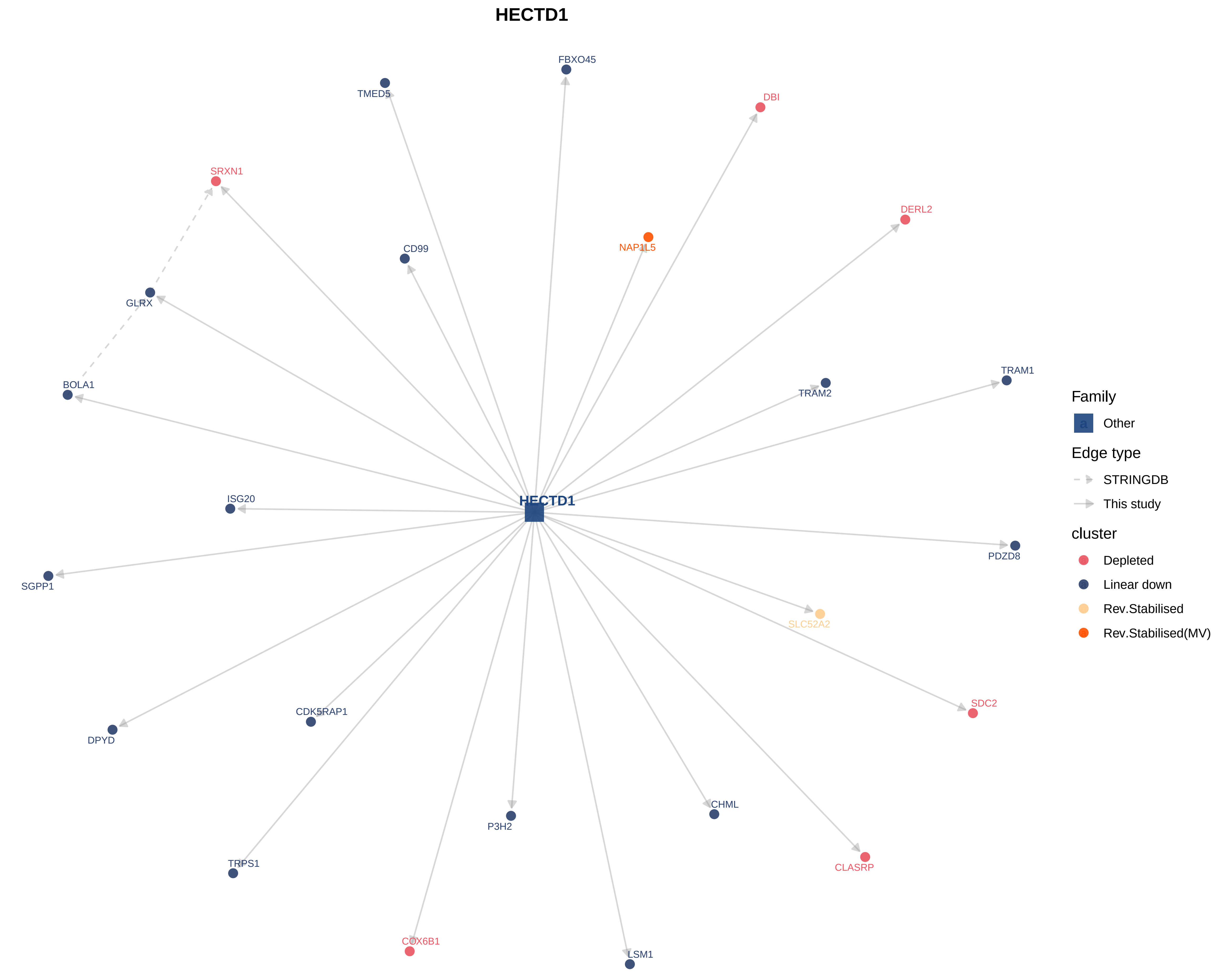

HECTD1
FBXO45
TMED5
DBI
SRXN1
DERL2
CD99
NAP1L5
GLRX
BOLA1
TRAM1
TRAM2
HECTD1
ISG20
PDZD8
SGPP1
SLC52A2
SDC2
CDK5RAP1
DPYD
CHML
P3H2
CLASRP
TRPS1
COX6B1
LSM1
Family
Other
Edge type
STRINGDB
This study
cluster
Depleted
Linear down
Rev.Stabilised
Rev.Stabilised(MV)

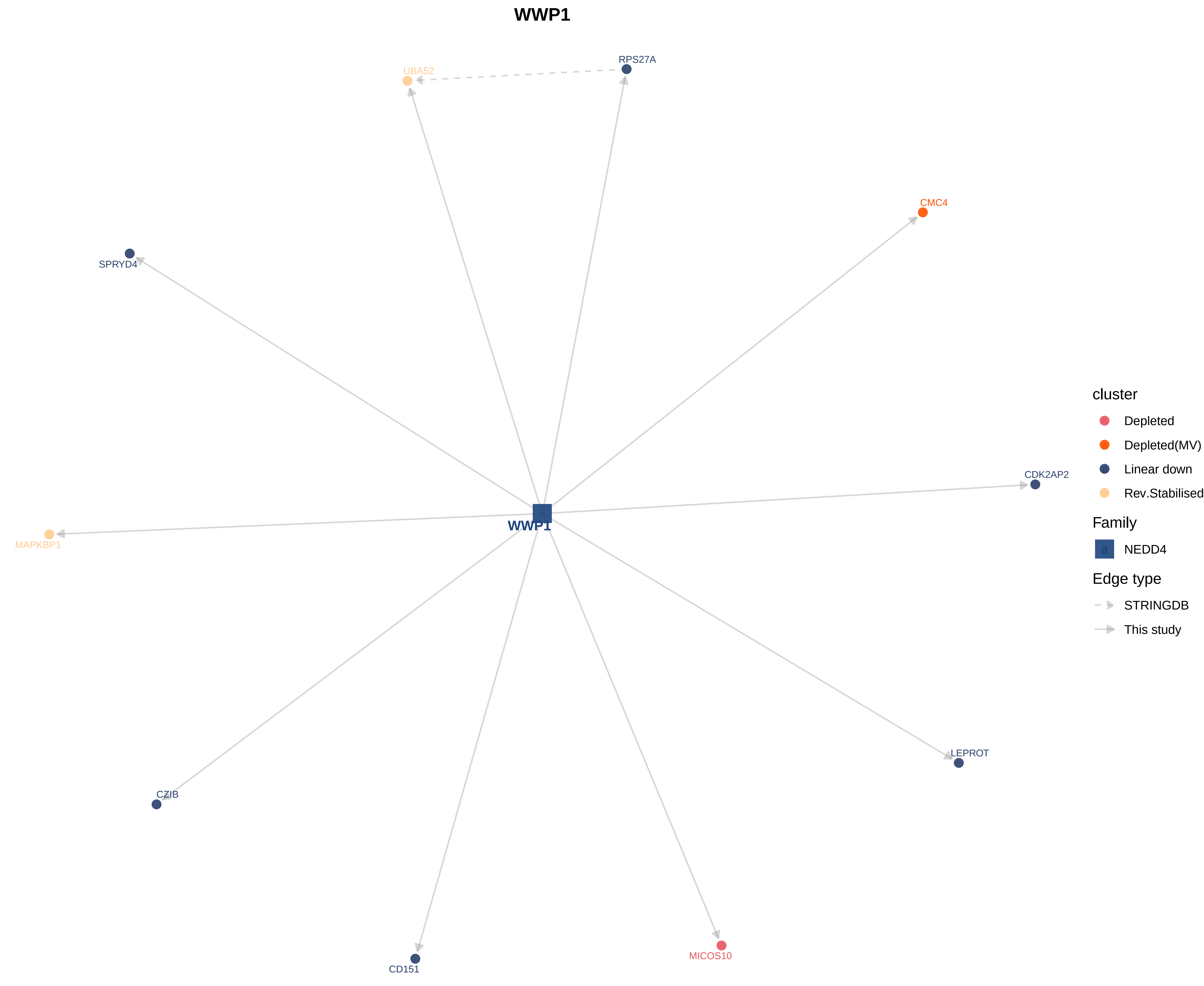

WWP1
RPS27A
UBA52
CMC4
SPRYD4
CDK2AP2
WWP1
MAPKBP1
LEPROT
CZIB
CD151
MICOS10
cluster
Depleted
Depleted(MV)
Linear down
Rev.Stabilised
Family
NEDD4
Edge type
STRINGDB
This study

#### AREL1

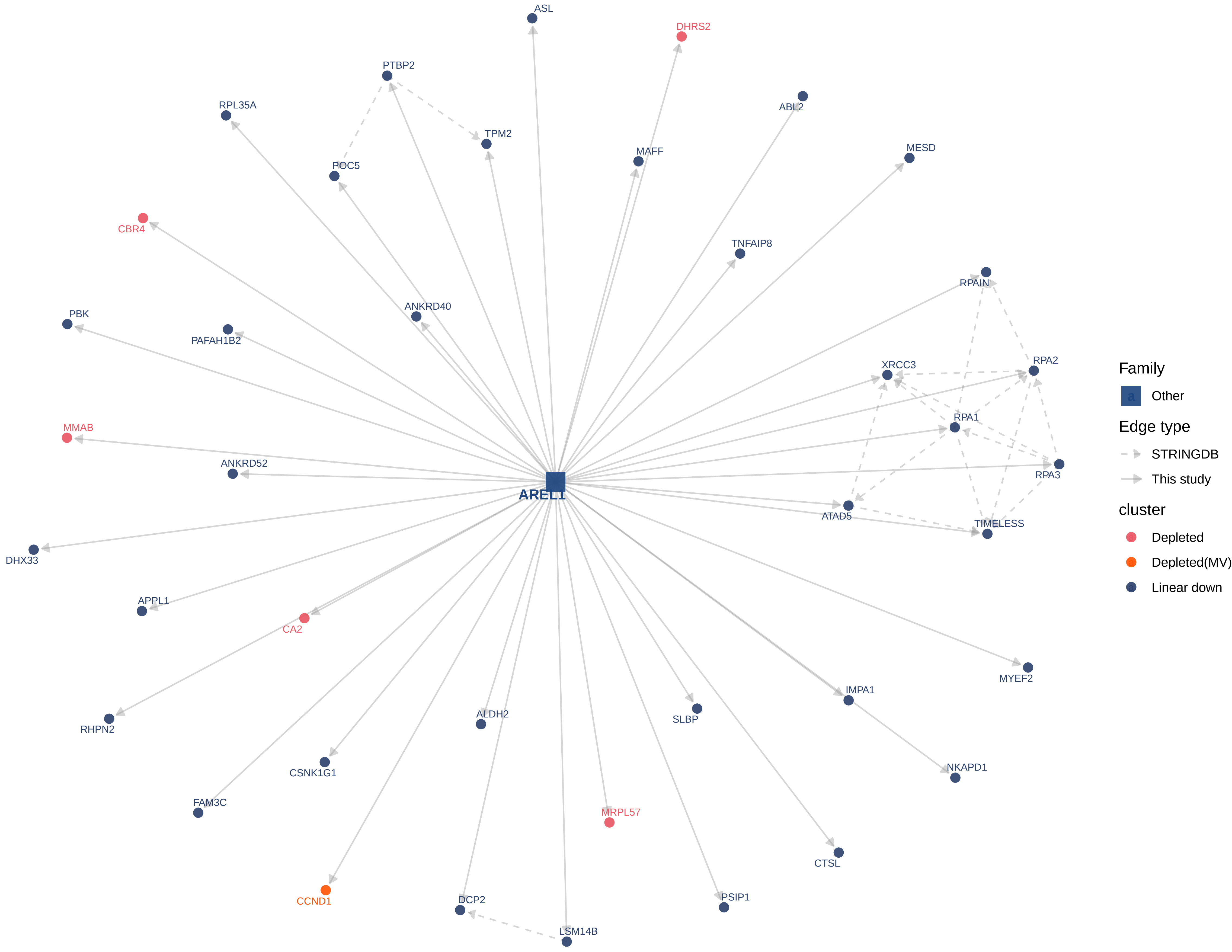
ASL
DHRS2
PTBP2
RPL35A
TPM2
ABL2
MESD
MAFF
POC5
CBR4
TNFAIP8
RPAIN
PBK
ANKRD40
PAFAH1B2
XRCC3
RPA2
RPA1
MMAB
ANKRD52
RPA3
AREL1
ATAD5
TIMELESS
DHX33
APPL1
CA2
MYEF2
IMPA1
RHPN2
ALDH2
SLBP
CSNK1G1
NKAPD1
FAM3C
MRPL57
CTSL
CCND1
DCP2
PSIP1
LSM14B
Family
Other
Edge type
STRINGDB
This study
cluster
Depleted
Depleted(MV)
Linear down

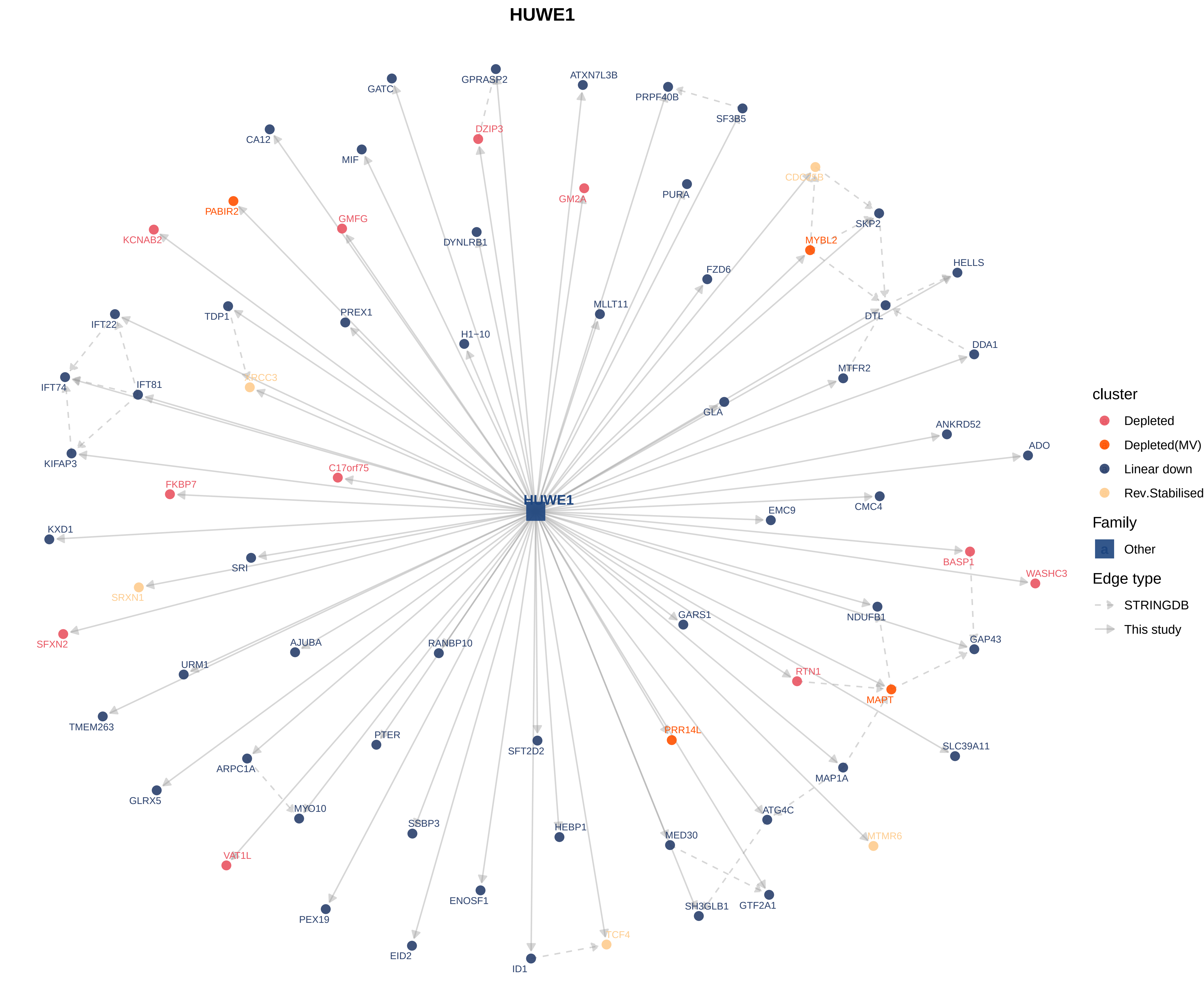
HUWE1
GATC
GPRASP2
ATXN7L3B
PRPF40B
SF3B5
CA12
DZIP3
MIF
GM2A
PURA
PABIR2
GMFG
KCNAB2
DYNLRB1
SKP2
MYBL2
HELLS
FZD6
DTL
IFT22
TDP1
PREX1
MLLT11
H1−10
DDA1
MTFR2
IFT74
IFT81
GLA
ANKRD52
ADO
KIFAP3
C17orf75
FKBP7
HUWE1
CMC4
EMC9
KXD1
BASP1
SRI
WASHC3
SRXN1
GARS1
NDUFB1
SFXN2
AJUBA
RANBP10
GAP43
URM1
RTN1
MAPT
TMEM263
PTER
PRR14L
SFT2D2
SLC39A11
ARPC1A
MAP1A
GLRX5
MYO10
ATG4C
SSBP3
HEBP1
MTMR6
MED30
VAT1L
ENOSF1
SH3GLB1
GTF2A1
PEX19
EID2
TCF4
ID1
cluster
Depleted
Depleted(MV)
Linear down
Rev.Stabilised
Family
Other
Edge type
STRINGDB
This study

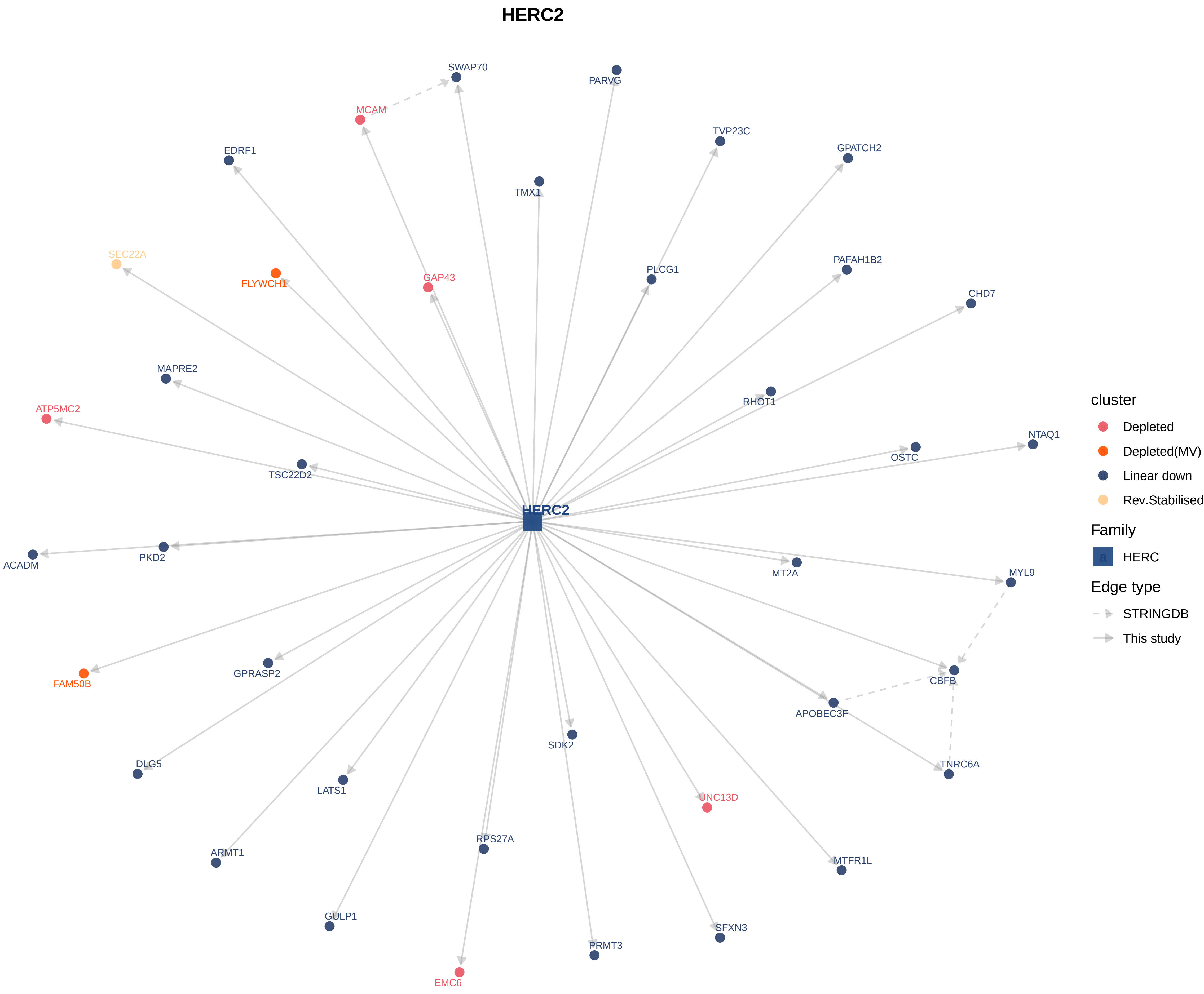
HERC2
SWAP70
PARVG
MCAM
TVP23C
GPATCH2
EDRF1
TMX1
SEC22A
FLYWCH1
GAP43
PLCG1
PAFAH1B2
CHD7
MAPRE2
RHOT1
ATP5MC2
TSC22D2
OSTC
NTAQ1
HERC2
ACADM
PKD2
MT2A
MYL9
GPRASP2
FAM50B
CBFB
APOBEC3F
SDK2
DLG5
TNRC6A
LATS1
UNC13D
RPS27A
ARMT1
MTFR1L
GULP1
SFXN3
PRMT3
EMC6
cluster
Depleted
Depleted(MV)
Linear down
Rev.Stabilised
Family
HERC
Edge type
STRINGDB
This study

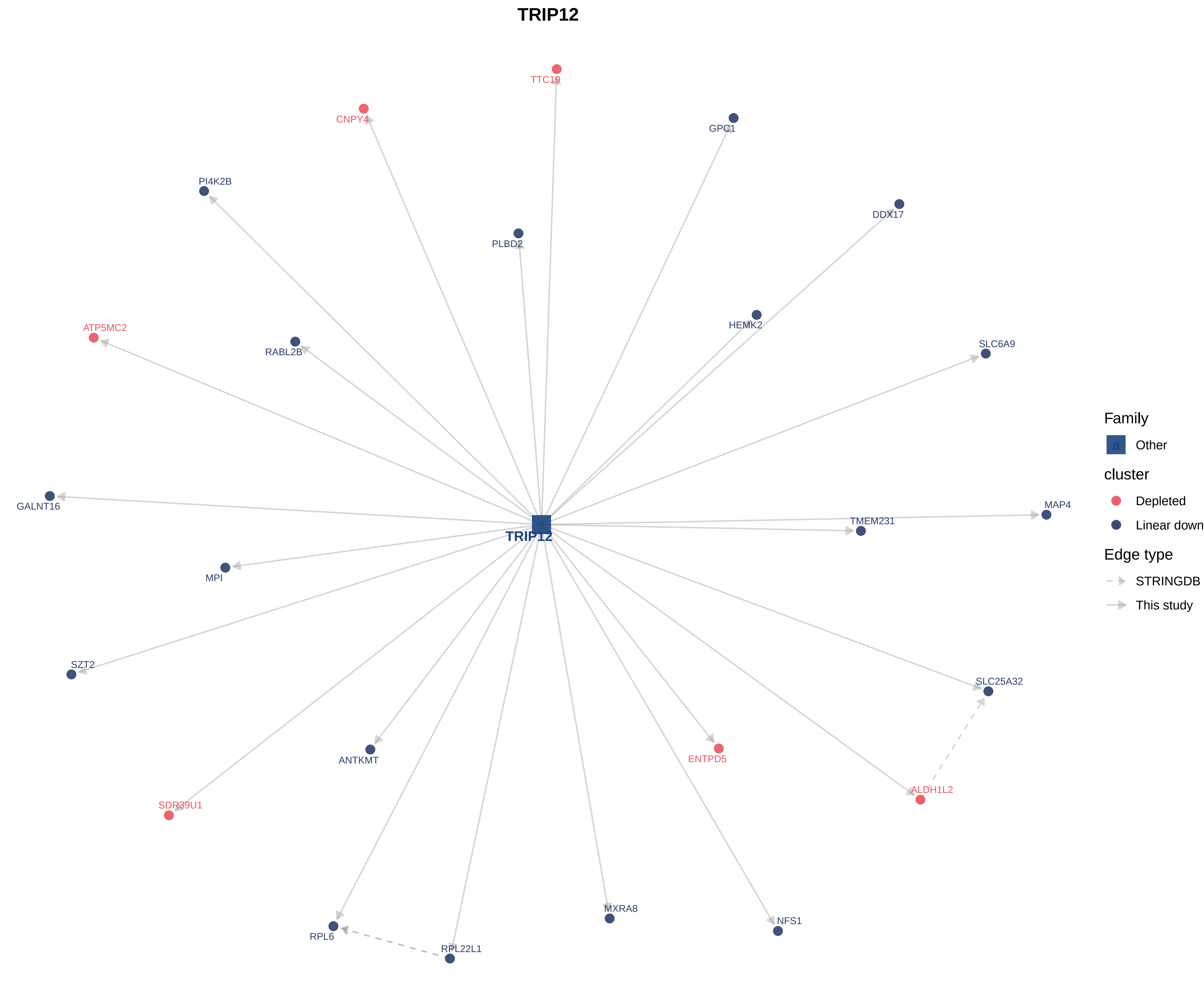
TRIP12
TTC19
CNPY4
GPC1
PI4K2B
DDX17
PLBD2
HEMK2
SLC6A9
ATP5MC2
RABL2B
GALNT16
MAP4
TMEM231
TRIP12
MPI
SZT2
SLC25A32
ANTKMT
ENTPD5
ALDH1L2
SDR39U1
MXRA8
NFS1
RPL6
RPL22L1
Family
Other
cluster
Depleted
Linear down
Edge type
STRINGDB
This study

HECW2

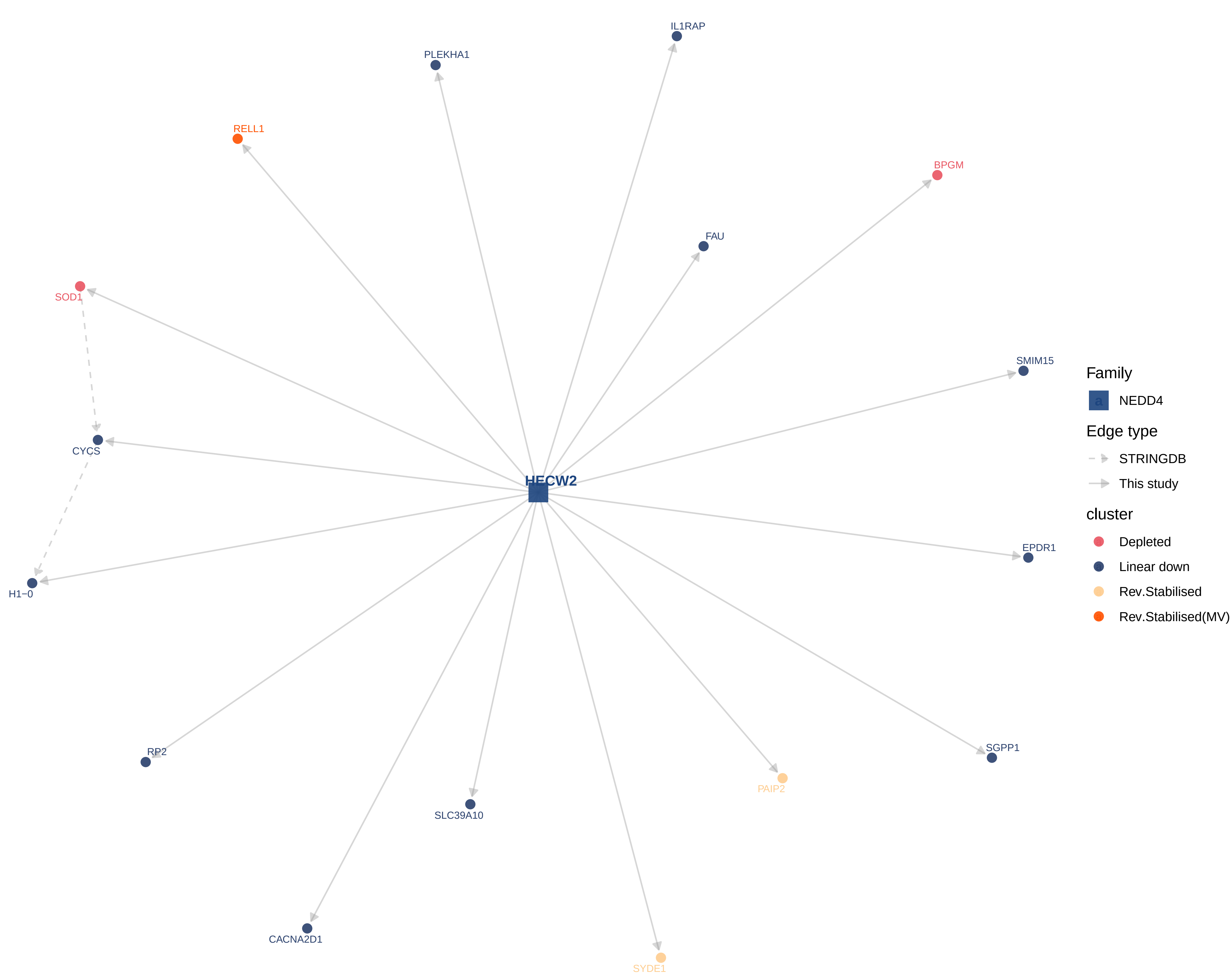

IL1RAP
PLEKHA1
RELL1
BPGM
FAU
SOD1
SMIM15
CYCS
HECW2
EPDR1
H1−0
RP2
SGPP1
PAIP2
SLC39A10
CACNA2D1
SYDE1
Family
NEDD4
Edge type
STRINGDB
This study
cluster
Depleted
Linear down
Rev.Stabilised
Rev.Stabilised(MV)

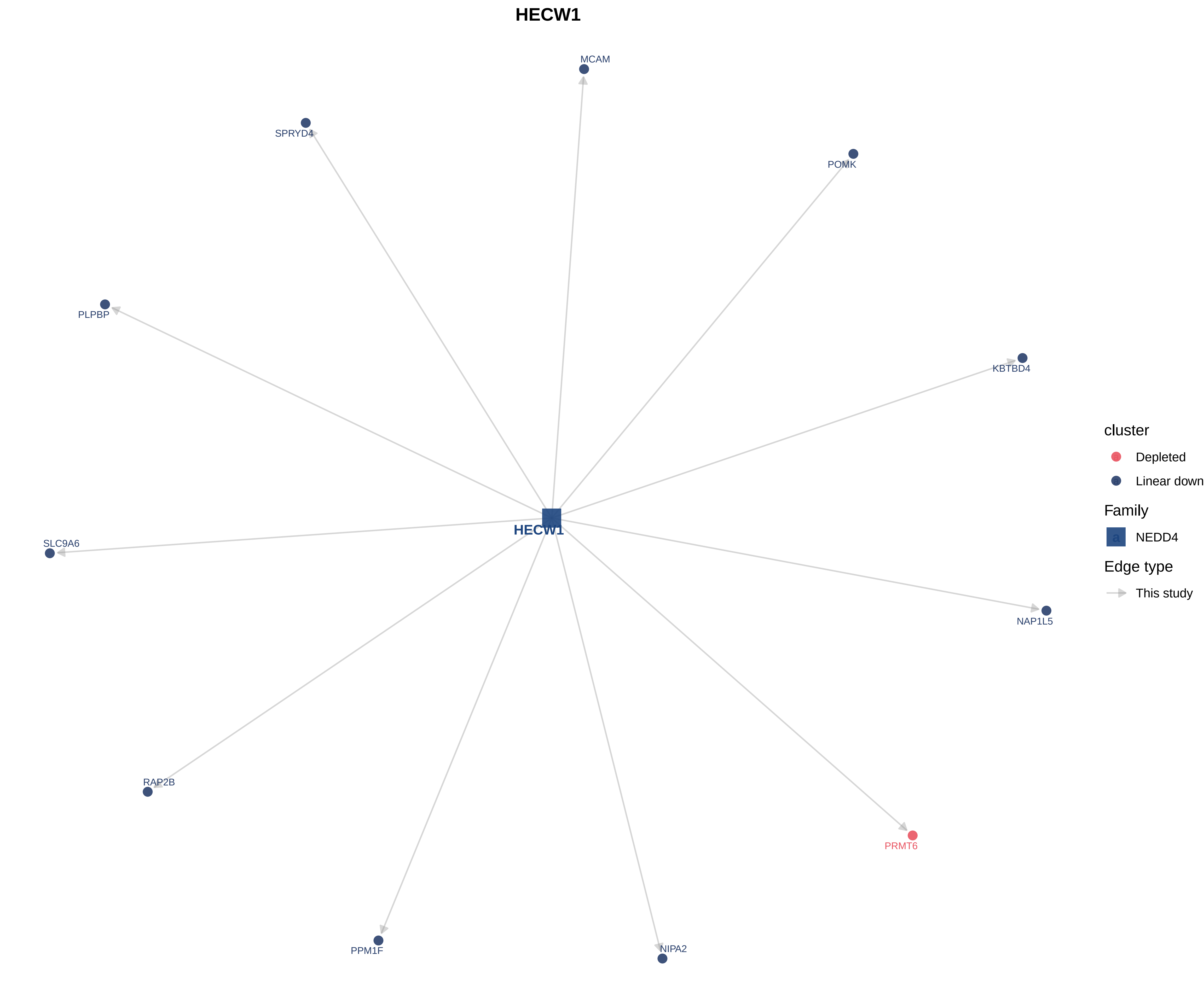
HECW1
MCAM
SPRYD4
POMK
PLPBP
KBTBD4
HECW1
SLC9A6
NAP1L5
RAP2B
PRMT6
PPM1F
NIPA2
cluster
Depleted
Linear down
Family
NEDD4
Edge type
This study

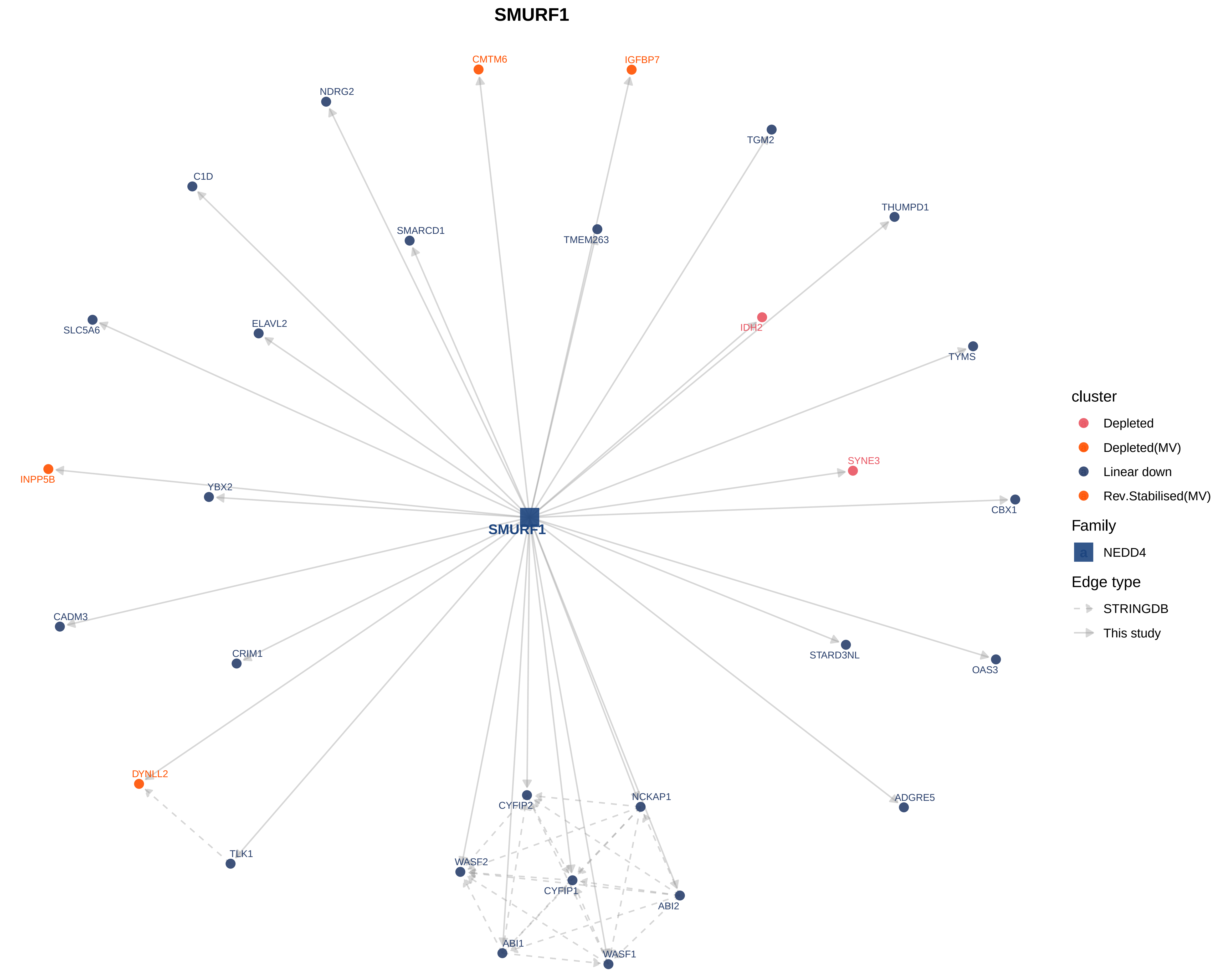

SMURF1
CMTM6
IGFBP7
NDRG2
TGM2
C1D
THUMPD1
SMARCD1
TMEM263
IDH2
SLC5A6
ELAVL2
TYMS
cluster
Depleted
Depleted(MV)
Linear down
Rev.Stabilised(MV)
SYNE3
INPP5B
YBX2
CBX1
SMURF1
Family
NEDD4
Edge type
STRINGDB
This study
CADM3
CRIM1
STARD3NL
OAS3
DYNLL2
NCKAP1
ADGRE5
CYFIP2
TLK1
WASF2
CYFIP1
ABI2
ABI1
WASF1

### HECTD4

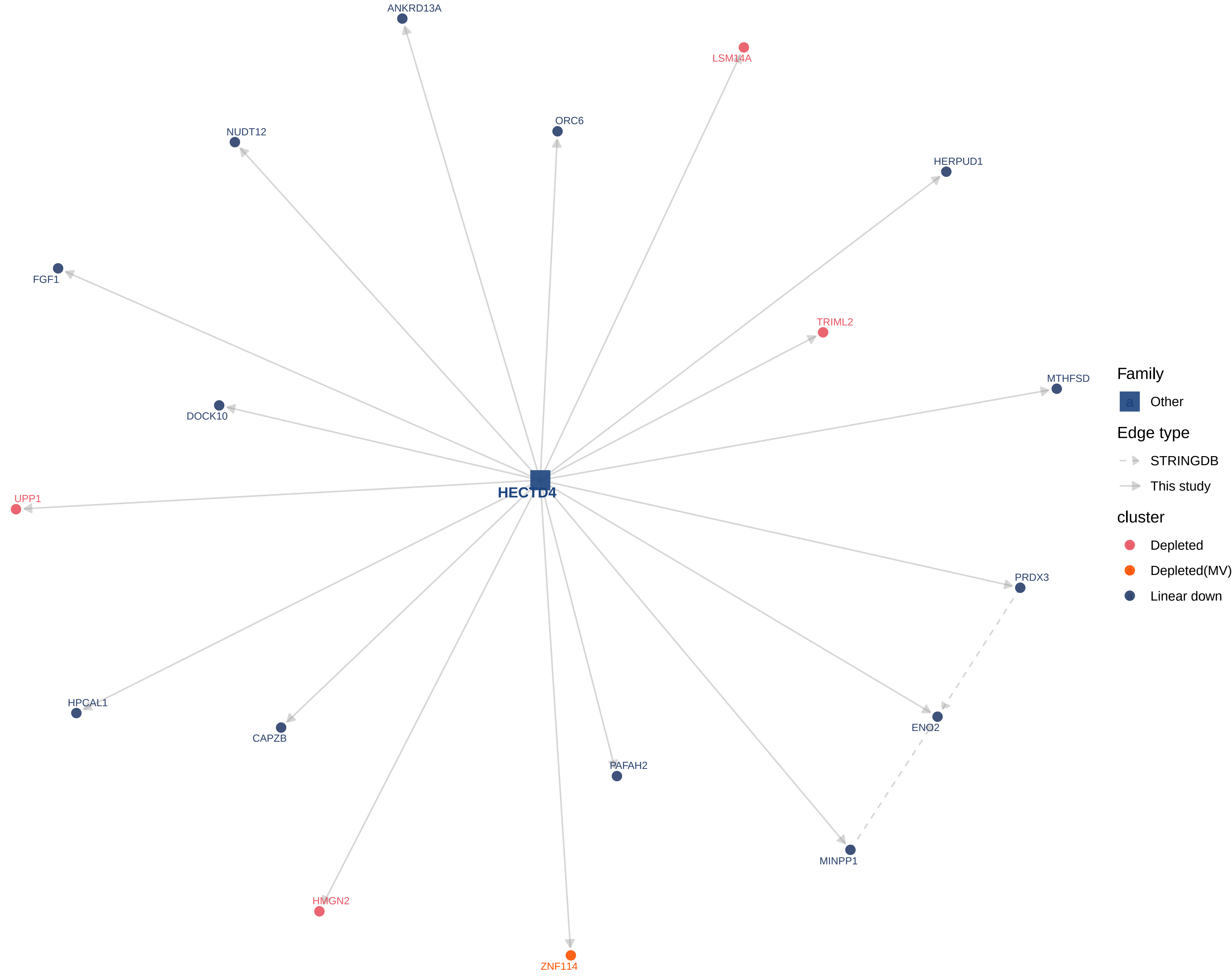
ANKRD13A
LSM14A
NUDT12
ORC6
HERPUD1
FGF1
TRIML2
MTHFSD
DOCK10
HECTD4
UPP1
PRDX3
HPCAL1
CAPZB
ENO2
PAFAH2
MINPP1
HMGN2
ZNF114
Family
Other
Edge type
STRINGDB
This study
cluster
Depleted
Depleted(MV)
Linear down

# G2E3

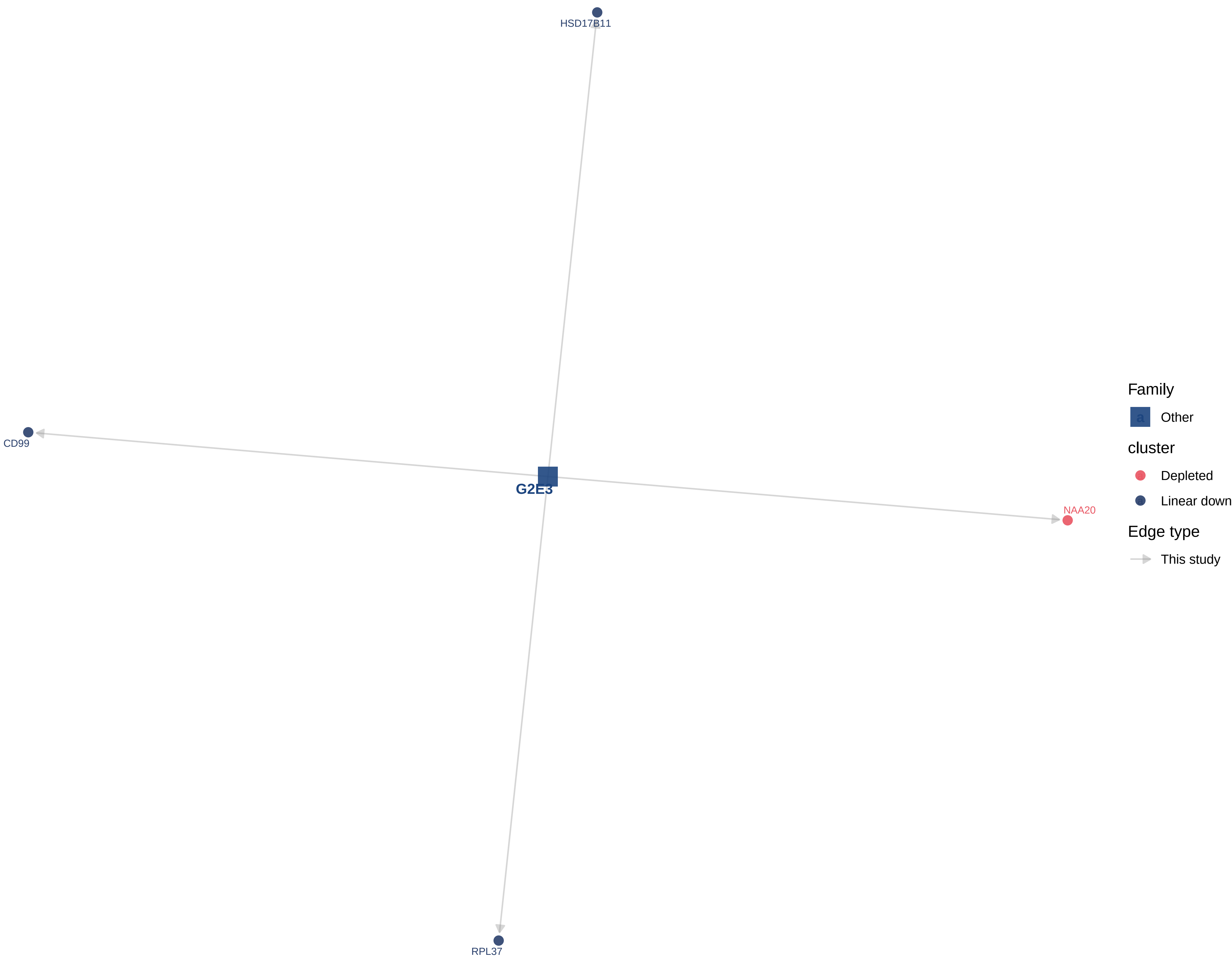

HSD17B11
CD99
G2E3
NAA20
RPL37
Family
Other
cluster
Depleted
Linear down
Edge type
This study

#### HERC1

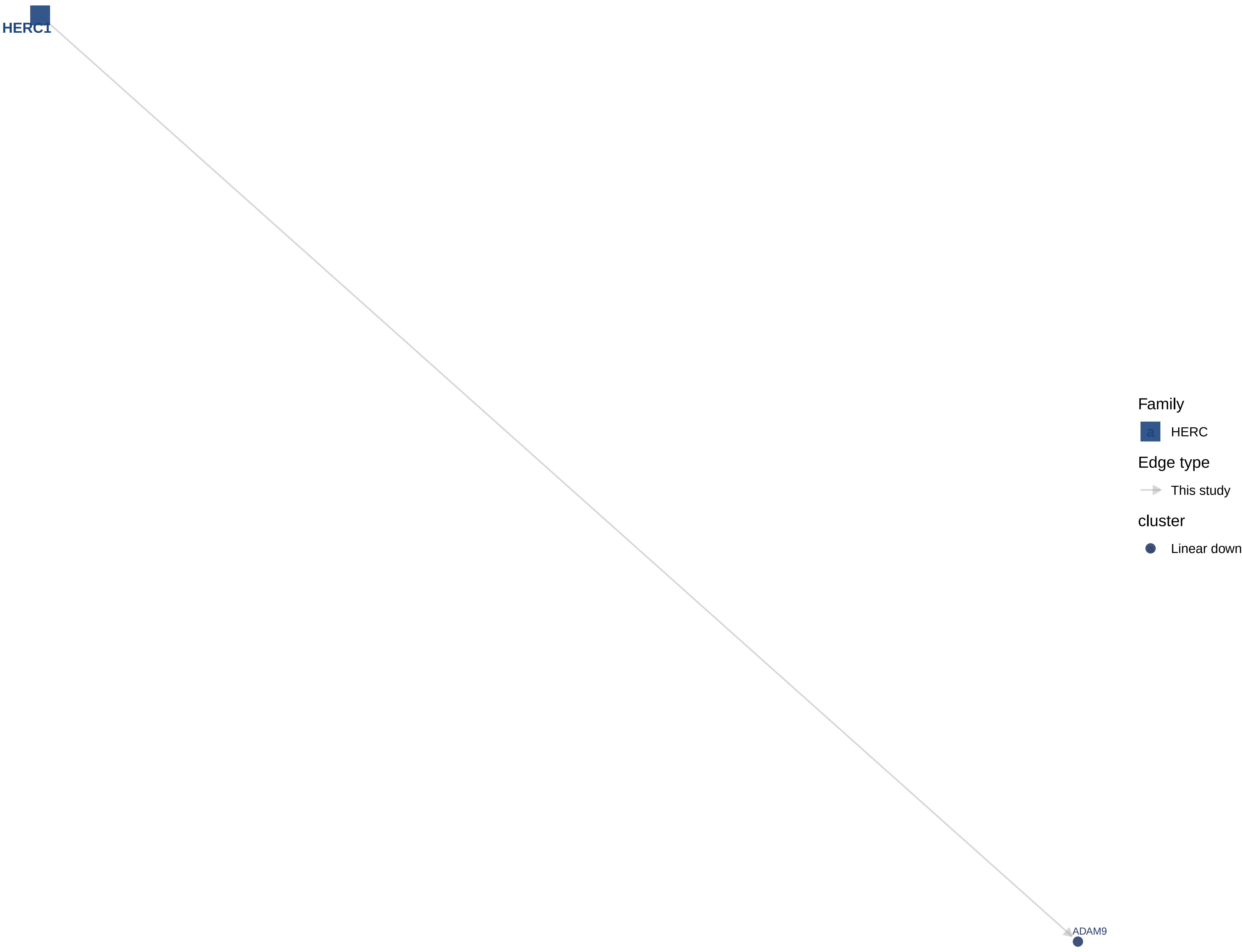

HERC1
ADAM9
Family
HERC
Edge type
This study
cluster
Linear down
