## Supplementary File 1 for "A dynamic proteomic HECT E3 ligase screen reveals substrate exclusivity and redundancies"

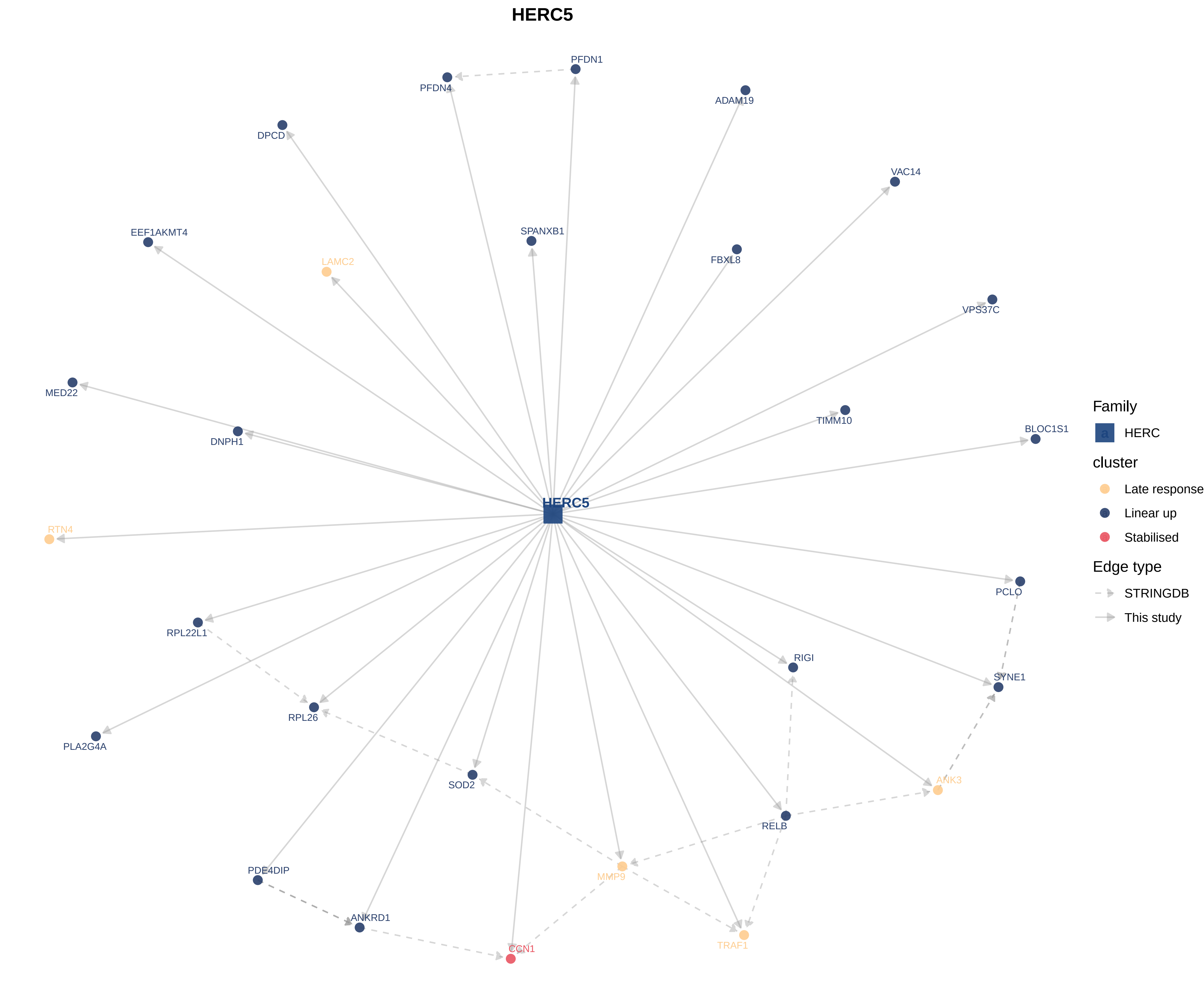
HERC5
PFDN1
PFDN4
ADAM19
DPCD
VAC14
EEF1AKMT4
SPANXB1
LAMC2
FBXL8
VPS37C
MED22
TIMM10
DNPH1
BLOC1S1
HERC5
RTN4
PCLO
RPL22L1
RIGI
SYNE1
RPL26
PLA2G4A
SOD2
ANK3
RELB
PDE4DIP
MMP9
ANKRD1
CCN1
TRAF1
Family
HERC
cluster
Late response
Linear up
Stabilised
Edge type
STRINGDB
This study

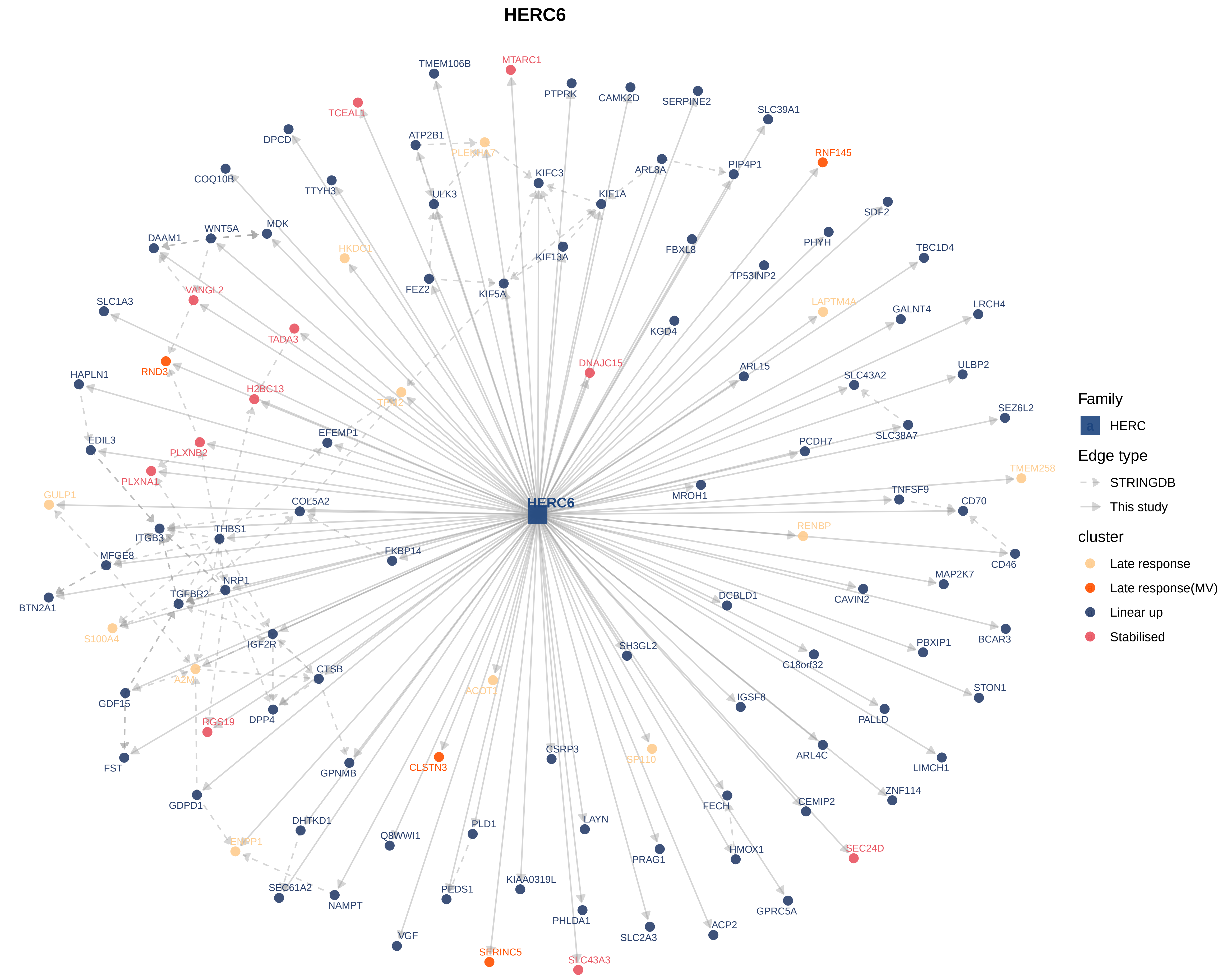

HERC6
TMEM106B
MTARC1
PTPRK
CAMK2D
SERPINE2
SLC39A1
TCEAL1
DPCD
ATP2B1
PLEKHA7
RNF145
COQ10B
TTYH3
ULK3
KIFC3
ARL8A
PIP4P1
KIF1A
SDF2
DAAM1
WNT5A
MDK
HKDC1
KIF13A
FBXL8
PHYH
TBC1D4
FEZ2
KIF5A
TP53INP2
SLC1A3
VANGL2
LAPTM4A
GALNT4
LRCH4
TADA3
KGD4
HAPLN1
RND3
H2BC13
TPM2
DNAJC15
ARL15
SLC43A2
ULBP2
EFEMP1
SEZ6L2
EDIL3
PLXNB2
PCDH7
SLC38A7
PLXNA1
TMEM258
GULP1
COL5A2
MROH1
TNFSF9
CD70
ITGB3
THBS1
RENBP
MFGE8
FKBP14
CD46
NRP1
MAP2K7
BTN2A1
TGFBR2
DCBLD1
CAVIN2
S100A4
IGF2R
BCAR3
SH3GL2
PBXIP1
C18orf32
A2M
CTSB
ACOT1
STON1
GDF15
IGSF8
PALLD
RGS19
DPP4
FST
GPNMB
CLSTN3
CSRP3
SP110
ARL4C
LIMCH1
ZNF114
GDPD1
FECH
CEMIP2
DHTKD1
LAYN
PLD1
Q8WWI1
ENPP1
HMOX1
SEC24D
PRAG1
SEC61A2
PEDS1
KIAA0319L
NAMPT
GPRC5A
PHLDA1
ACP2
SLC2A3
VGF
SERINC5
SLC43A3
Family
HERC
Edge type
STRINGDB
This study
cluster
Late response
Late response(MV)
Linear up
Stabilised

#### UBR5

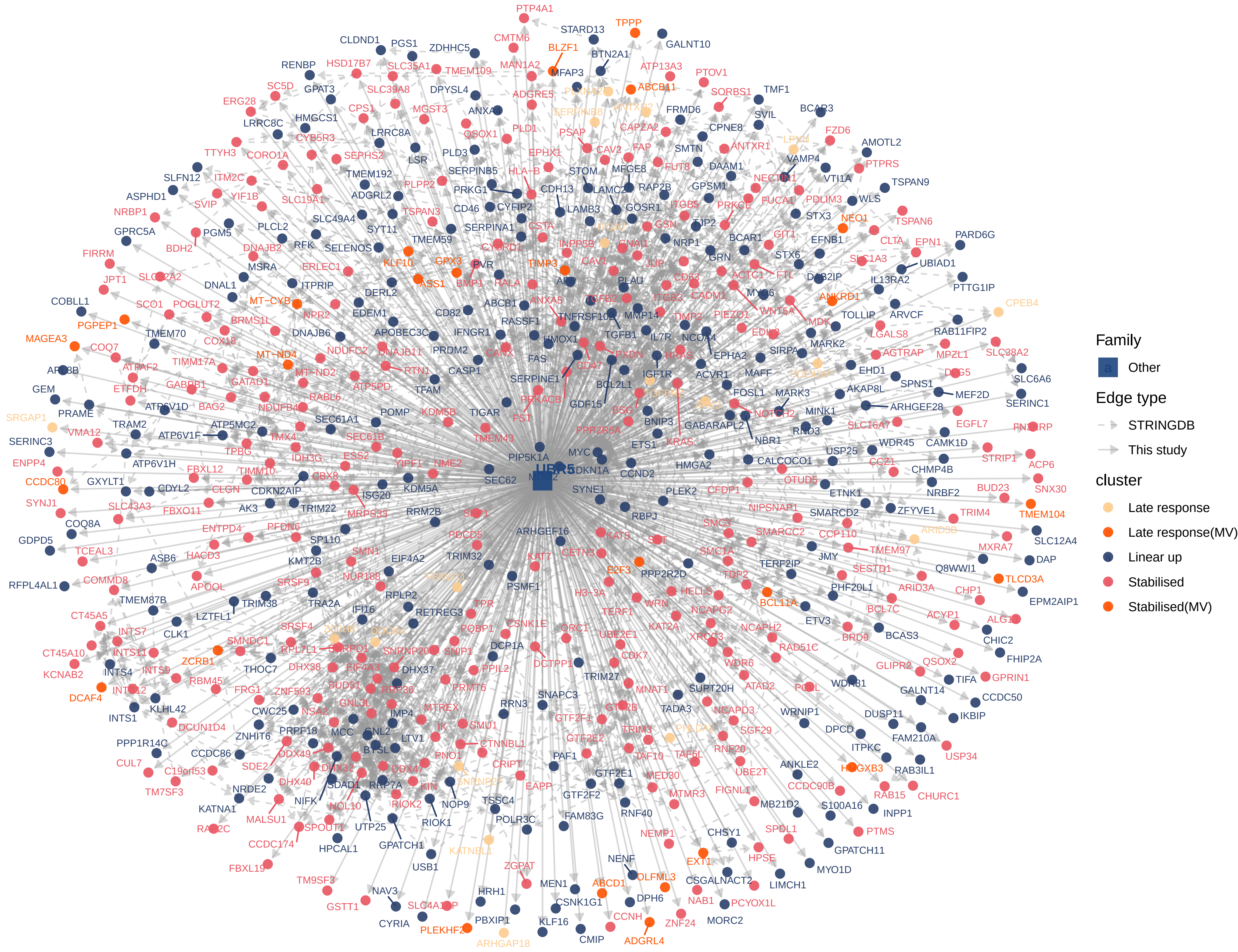

Family
Other
Edge type
STRINGDB
This study
cluster
Late response
Late response(MV)
Linear up
Stabilised
Stabilised(MV)
UBR5
PTP4A1
STARD13
TPPP
GALNT10
CLDND1
PGS1
ZDHHC5
CMTM6
BLZF1
BTN2A1
RENBP
HSD17B7
SLC35A1
TMEM109
MAN1A2
MFAP3
ATP13A3
PTOV1
SC5D
GPAT3
SLC39A8
DPYSL4
ADGRE5
PLXNB2
ABCB11
SORBS1
TMF1
ERG28
CPS1
MGST3
ANXA4
SERPINB8
ANTXR2
FRMD6
BCAP31
LRRC8C
HMGCS1
PLD1
PSAP
CAPZA2
SVIL
CYB5R3
LRRC8A
QSOX1
CPNE8
FZD6
TTYH3
CORO1A
SEPHS2
LSR
PLD3
EPHX1
CAV2
FAP
SMTN
ANTXR1
LPXN
AMOTL2
SLFN12
ITM2C
TMEM192
SERPINB5
HLA-B
STOM
MFGE8
FUT8
DAAM1
VAMP4
PTPRS
ASPHD1
YIF1B
SLC19A1
ADGRL2
PLPP2
PRKG1
CDH13
LAMC2
RAP2B
GPSM1
NECTIN1
VTI1A
TSPAN9
NRBP1
SVIP
CD46
CYFIP2
LAMB3
GOSR1
ITGB5
PRKCE
FUCA1
PDLIM3
WLS
GPRC5A
PGM5
SLC49A4
TSPAN3
SYT11
SERPINA1
CSTA
GSN
TJP2
STX3
NEO1
TSPAN6
BDH2
PLCL2
TMEM59
CYBRD1
INPP5B
GNAI1
NRP1
BCAR1
GIT1
EFNB1
CLTA
EPN1
PARD6G
FIRRM
DNAJB2
RFK
SELENOS
KLF10
GPX3
PVR
TIMP3
CAV1
JUP
GRN
STX6
SLC1A3
UBIAD1
JPT1
SLC2A2
MSRA
ERLEC1
ASS1
BMP1
RALA
ANPEP
PLAU
CD63
ACTC1
FTL
DAB2IP
IL13RA2
PTTG1IP
COBLL1
DNAL1
ITPRIP
DERL2
ABCB1
ANXA5
TGFB2
ITGB3
CADM1
MYO6
ANKRD1
CPEB4
SCO1
POGLUT2
MT-CYB
NPR2
EDEM1
CD82
RASSF1
TNFRSF10B
MMP14
TIMP2
PIEZO1
WNT5A
MDK
TOLLIP
ARVCF
PGPEP1
BRMS1L
DNAJB6
APOBEC3C
IFNGR1
HMOX1
TGFB1
IL7R
NCOA4
EDIL3
LGALS8
RAB11FIP2
TMEM70
MAGEA3
COQ7
COX18
NDUFC2
DNAJB11
PRDM2
CANX
FAS
PXDN
HRAS
EPHA2
SIRPA
MARK2
AGTRAP
MPZL1
SLC38A2
ARL8B
ATPAF2
TIMM17A
MT-ND4
RTN1
CASP1
CD47
IGF1R
ACVR1
MAFF
PODXL
EHD1
DCGG5
SLC6A6
MT-ND2
SERPINE1
BCL2L1
SPNS1
GEM
ETFDH
GABRB1
GATAD1
ATP5PD
TFAM
PRKACB
GDF15
TGFBR1
FOSL1
MARK3
AKAP8L
MEF2D
SERINC1
ATP6V1D
BAG2
RABL6
NDUFB4
PRAME
ARHGEF28
SRGAP1
POMP
KDM5B
TIGAR
FST
BSG
NOTCH2
MINK1
EGFL7
TRAM2
ATP5MC2
SEC61A1
BNIP3
GABARAPL2
SLC16A7
FNDC3B
VMA12
ATP6V1F
TMX4
SEC61B
TMEM43
PPP2R5A
KRAS
RND3
NBR1
WDR45
CAMK1D
SERINC3
TPBG
IDH3G
ESS2
PIP5K1A
MYC
ETS1
USP25
STRIP1
ACP6
ATP6V1H
FBXL12
TIMM10
YIPF1
NME2
CDKN1A
CCND2
HMGA2
CALCOCO1
CCZ1
CHMP4B
ENPP4
CBX8
SEC62
MDM2
OTUD5
CCDC80
GXYLT1
CDYL2
CLGN
CDKN2AIP
KDM5A
SYNE1
PLEK2
CFDP1
ETNK1
NRBF2
BUD23
SNX30
SYNJ1
ISG20
SLC43A3
FBXO11
AK3
TRIM22
MRPS33
RRM2B
SNRPB1
RBPJ
NIPSNAP1
SMARCD2
ZFYVE1
TRIM4
TMEM104
COQ8A
ENTPD4
PFDN6
SMC3
GDPD5
SP110
PDCD5
ARHGEF16
KAT5
SRIT
SMARCC2
CCP110
ARID3B
SLC12A4
TCEAL3
HACD3
SMN1
KMT2B
TRIM32
KAT7
CETN3
SMC1A
TERF2IP
JMY
TMEM97
MXRA7
ASB6
EIF4A2
DAP
COMMD8
SRSF9
NUP188
PRPF40A
PSMF1
E2F3
PPP2R2D
TDP2
SESTD1
Q8WWI1
TLCD3A
RFPL4AL1
APOOL
H3-3A
HELLS
BCL11A
PHF20L1
ARID3A
CHP1
TMEM87B
TRIM38
TRA2A
RPLP2
IFI16
TPR
RETREG3
WRN
EPM2AIP1
CT45A5
LZTFL1
TERF1
NCAPG2
BCL7C
ACYP1
ALG10
INTS7
CLK1
SRSF4
ZC3H8
DOCK8
PQBP1
CSNK1E
ORC1
UBE2E1
KAT2A
NCAPH2
ETV3
BRD9
BCAS3
CHIC2
CT45A10
INTS11
SMNDC1
RPL7L1
SNRPD1
SNRNP200
SNIP1
DCP1A
XRCC3
RAD51C
FHIP2A
KCNAB2
INTS4
INTS9
ZCRB1
THOC7
DHX38
EIF4A3
DHX37
PPIL2
DCTPP1
CDK7
WDR6
GLIPR2
QSOX2
TIFA
GPRIN1
RBM45
TRIM27
SUPT20H
DCAF4
INTS12
FRG1
ZNF593
BUD31
RRP36
PRMT6
SNAPC3
MNAT1
ATAD2
PCNL
WDR81
GALNT14
CCDC50
INTS1
KLHL42
CWC25
NSA2
GNL3L
IMP4
MTREX
RRN3
GTF2F1
GTF2B
TADA3
NCAPD3
WRNIP1
DUSP11
IKBIP
DCUN1D4
PRPF18
MCC
GNL2
SMU1
TRIM3
PHLDA2
SGF29
DPCD
FAM210A
PPP1R14C
ZNHIT6
LTV1
IK
GTF2E2
RNF20
ITPKC
USP34
CCDC86
DDX49
BYSL
CTNNBL1
PAF1
TAF10
TAF6L
CUL7
C19orf53
SDE2
DHX32
PNO1
CRIPT
UBE2T
ANKLE2
HOXB3
RAB3IL1
TM7SF3
NRDE2
DHX40
SDAD1
DDX47
RRP7A
SNRNP27
GTF2E1
MED30
CCDC90B
RAB15
CHURC1
KATNA1
NIFK
KIN
EAPP
FIGNL1
MALSU1
NOL10
RIOK2
NOP9
TSSC4
GTF2F2
MTMR3
MB21D2
S100A16
INPP1
RAB2C
SPOUT1
UTP25
RIOK1
POLR3C
FAM83G
RNF40
SPDL1
PTMS
CCDC174
HPCAL1
GPATCH1
KATNBL1
NEMP1
CHSY1
GPATCH11
FBXL19
USB1
NENF
EXT1
HPSE
MYO1D
ZGPAT
OLFML3
CSGALNACT2
LIMCH1
TM9SF3
NAV3
MEN1
ABCD1
HRH1
GSTT1
SLC4A1AP
CSNK1G1
DPH6
NAB1
PCYOX1L
CYRIA
PLEKHF2
PBXIP1
CCNH
KLF16
ZNF24
MORC2
ARHGAP18
CMIP
ADGRL4

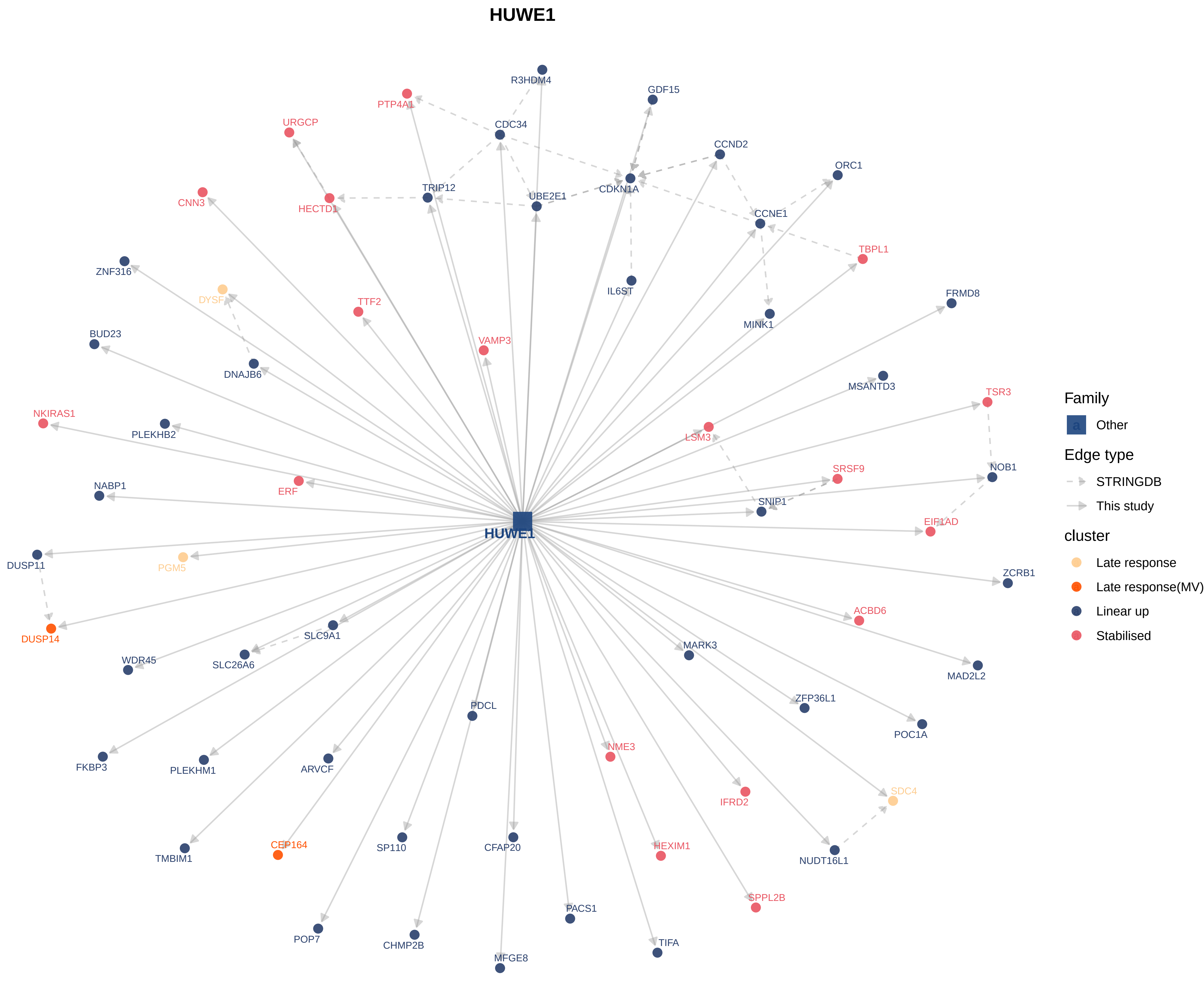
HUWE1
R3HDM4
GDF15
PTP4A1
CDC34
URGCP
CCND2
ORC1
TRIP12
UBE2E1
CDKN1A
CNN3
HECTD1
CCNE1
TBPL1
ZNF316
IL6ST
DYSF
TTF2
MINK1
FRMD8
BUD23
VAMP3
DNAJB6
MSANTD3
TSR3
NKIRAS1
PLEKHB2
LSM3
NOB1
SRSF9
NABP1
ERF
SNIP1
EIF1AD
HUWE1
DUSP11
PGM5
ZCRB1
ACBD6
DUSP14
SLC9A1
MARK3
WDR45
SLC26A6
MAD2L2
ZFP36L1
PDCL
POC1A
FKBP3
PLEKHM1
ARVCF
NME3
IFRD2
SDC4
CEP164
TMBIM1
SP110
CFAP20
HEXIM1
NUDT16L1
SPPL2B
PACS1
POP7
CHMP2B
TIFA
MFGE8
Family
Other
Edge type
STRINGDB
This study
cluster
Late response
Late response(MV)
Linear up
Stabilised

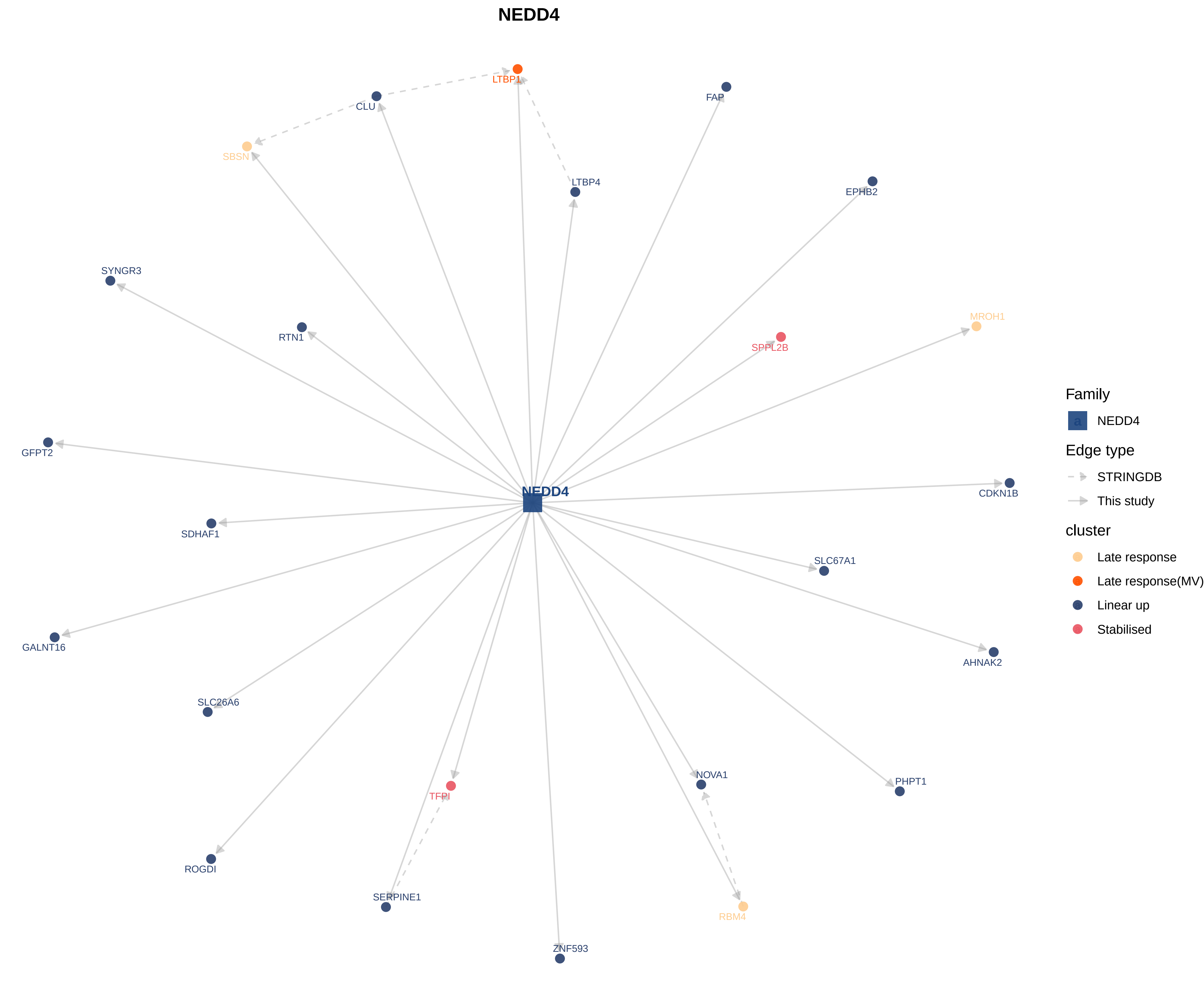
NEDD4
LTBP1
CLU
FAP
SBSN
LTBP4
EPHB2
SYNGR3
RTN1
MROH1
SPPL2B
GFPT2
NEDD4
CDKN1B
SDHAF1
SLC67A1
GALNT16
AHNAK2
SLC26A6
NOVA1
PHPT1
TFPI
ROGDI
SERPINE1
RBM4
ZNF593
Family
NEDD4
Edge type
STRINGDB
This study
cluster
Late response
Late response(MV)
Linear up
Stabilised

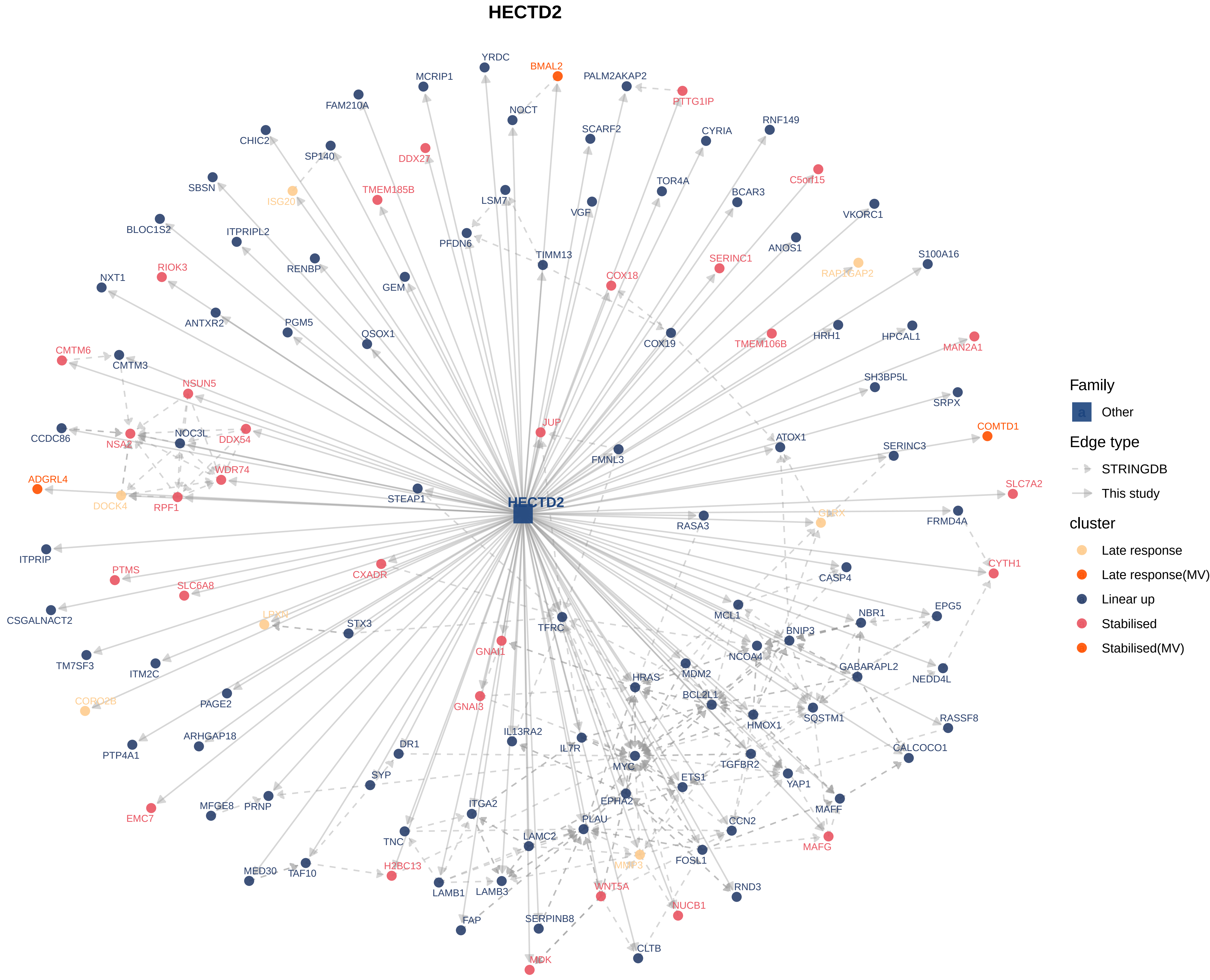
HECTD2
YRDC
BMAL2
MCRIP1
PALM2AKAP2
PTTG1IP
FAM210A
NOCT
CHIC2
SCARF2
CYRIA
RNF149
SP140
DDX27
SBSN
ISG20
TMEM185B
LSM7
TOR4A
BCAR3
C5orf15
VGF
VKORC1
BLOC1S2
ITPRIPL2
PFDN6
TIMM13
ANOS1
S100A16
RENBP
GEM
COX18
SERINC1
RAP1GAP2
NXT1
RIOK3
ANTXR2
PGM5
QSOX1
COX19
TMEM106B
HRH1
HPCAL1
MAN2A1
CMTM6
CMTM3
NSUN5
SH3BP5L
SRPX
CCDC86
NOC3L
DDX54
NSA2
JUP
FMNL3
ATOX1
COMTD1
SERINC3
WDR74
ADGRL4
DOCK4
RPF1
STEAP1
SLC7A2
RASA3
GLRX
FRMD4A
ITPRIP
CYTH1
PTMS
CXADR
SLC6A8
CASP4
CSGALNACT2
LPXN
STX3
TFRC
MCL1
NBR1
EPG5
BNIP3
TM7SF3
ITM2C
GNAI1
NCOA4
MDM2
HRAS
GABARAPL2
NEDD4L
CORO2B
PAGE2
GNAI3
BCL2L1
HMOX1
SQSTM1
RASSF8
PTP4A1
ARHGAP18
IL13RA2
IL7R
CALCOCO1
DR1
MYC
TGFBR2
SYP
ETS1
YAP1
EMC7
MFGE8
PRNP
ITGA2
EPHA2
MAFF
PLAU
CCN2
TNC
LAMC2
FOSL1
MAFG
MED30
TAF10
H2BC13
MMP3
LAMB1
LAMB3
WNT5A
RND3
NUCB1
FAP
SERPINB8
CLTB
MDK
Family
Other
Edge type
STRINGDB
This study
cluster
Late response
Late response(MV)
Linear up
Stabilised
Stabilised(MV)

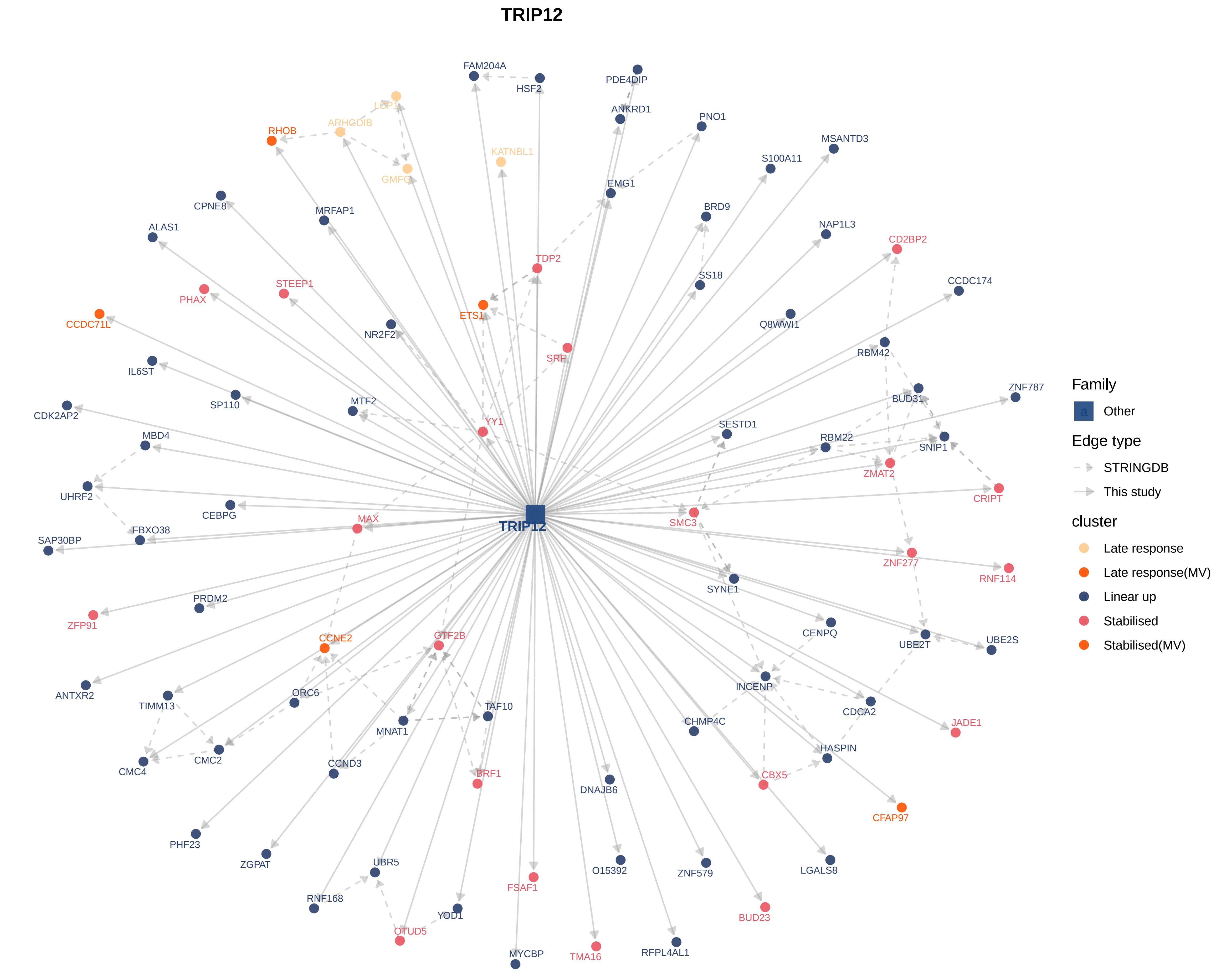
TRIP12
FAM204A
HSF2
PDE4DIP
ANKRD1
PNO1
MSANTD3
S100A11
LSP1
ARHGDIB
RHOB
KATNBL1
GMFG
EMG1
BRD9
NAP1L3
CD2BP2
CPNE8
MRFAP1
ALAS1
TDP2
SS18
CCDC174
PHAX
STEEP1
ETS1
CCDC71L
NR2F2
Q8WWI1
SRF
RBM42
IL6ST
SP110
MTF2
BUD31
ZNF787
CDK2AP2
YY1
SESTD1
RBM22
SNIP1
MBD4
ZMAT2
UHRF2
CRIPT
CEBPG
SMC3
MAX
TRIP12
FBXO38
SAP30BP
ZNF277
RNF114
SYNE1
PRDM2
ZFP91
CENPQ
UBE2T
UBE2S
CCNE2
GTF2B
ANTXR2
INCENP
TIMM13
ORC6
CDCA2
TAF10
JADE1
MNAT1
CHMP4C
HASPIN
CMC4
CMC2
CCND3
BRF1
CBX5
DNAJB6
CFAP97
PHF23
ZGPAT
O15392
ZNF579
LGALS8
UBR5
RNF168
FSAF1
YOD1
BUD23
OTUD5
MYCBP
TMA16
RFPL4AL1
Family
Other
Edge type
STRINGDB
This study
cluster
Late response
Late response(MV)
Linear up
Stabilised
Stabilised(MV)

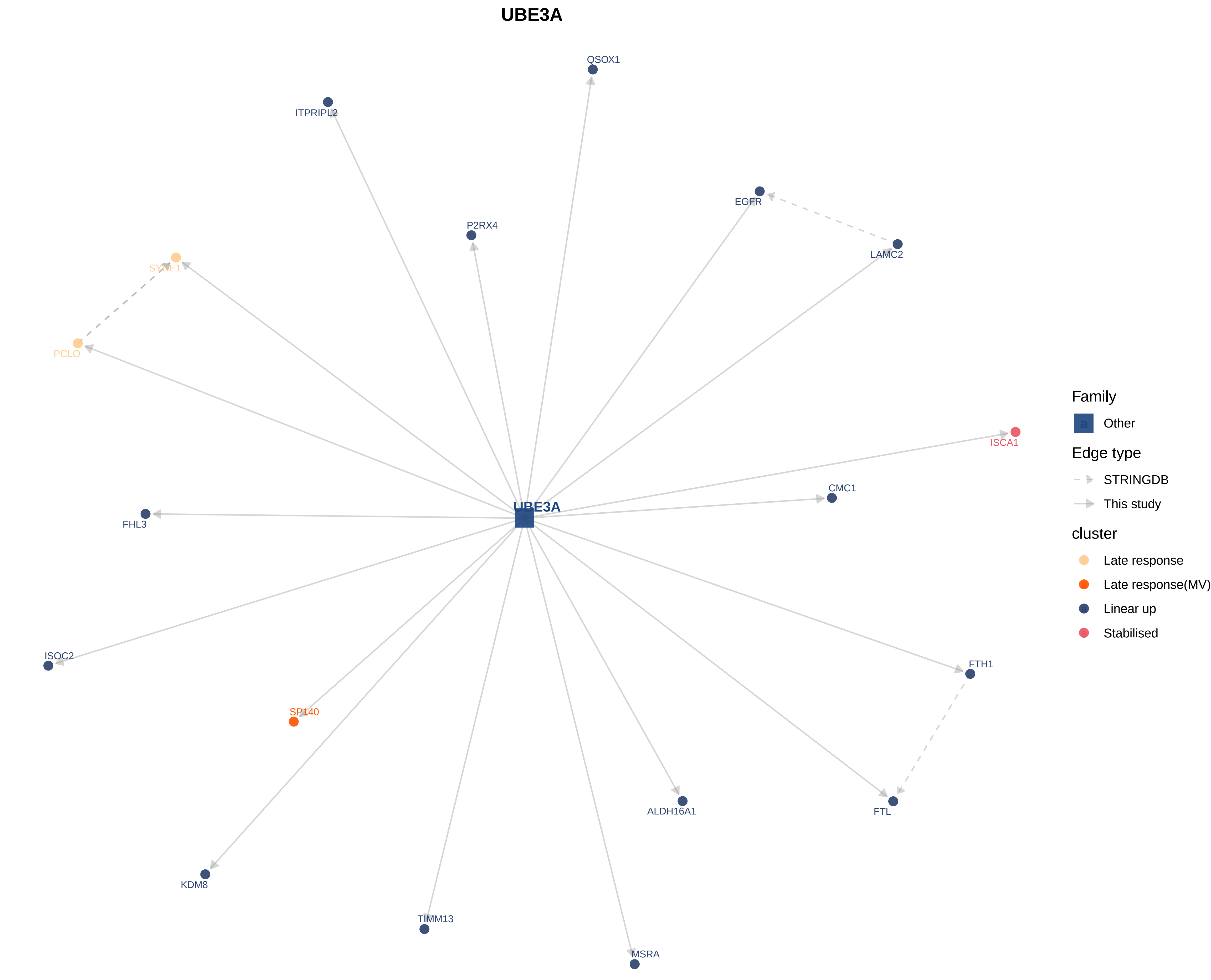

UBE3A
QSOX1
ITPRIPL2
EGFR
P2RX4
LAMC2
SYNE1
PCLO
ISCA1
CMC1
UBE3A
FHL3
ISOC2
SP140
FTH1
ALDH16A1
FTL
KDM8
TIMM13
MSRA
Family
Other
Edge type
STRINGDB
This study
cluster
Late response
Late response(MV)
Linear up
Stabilised

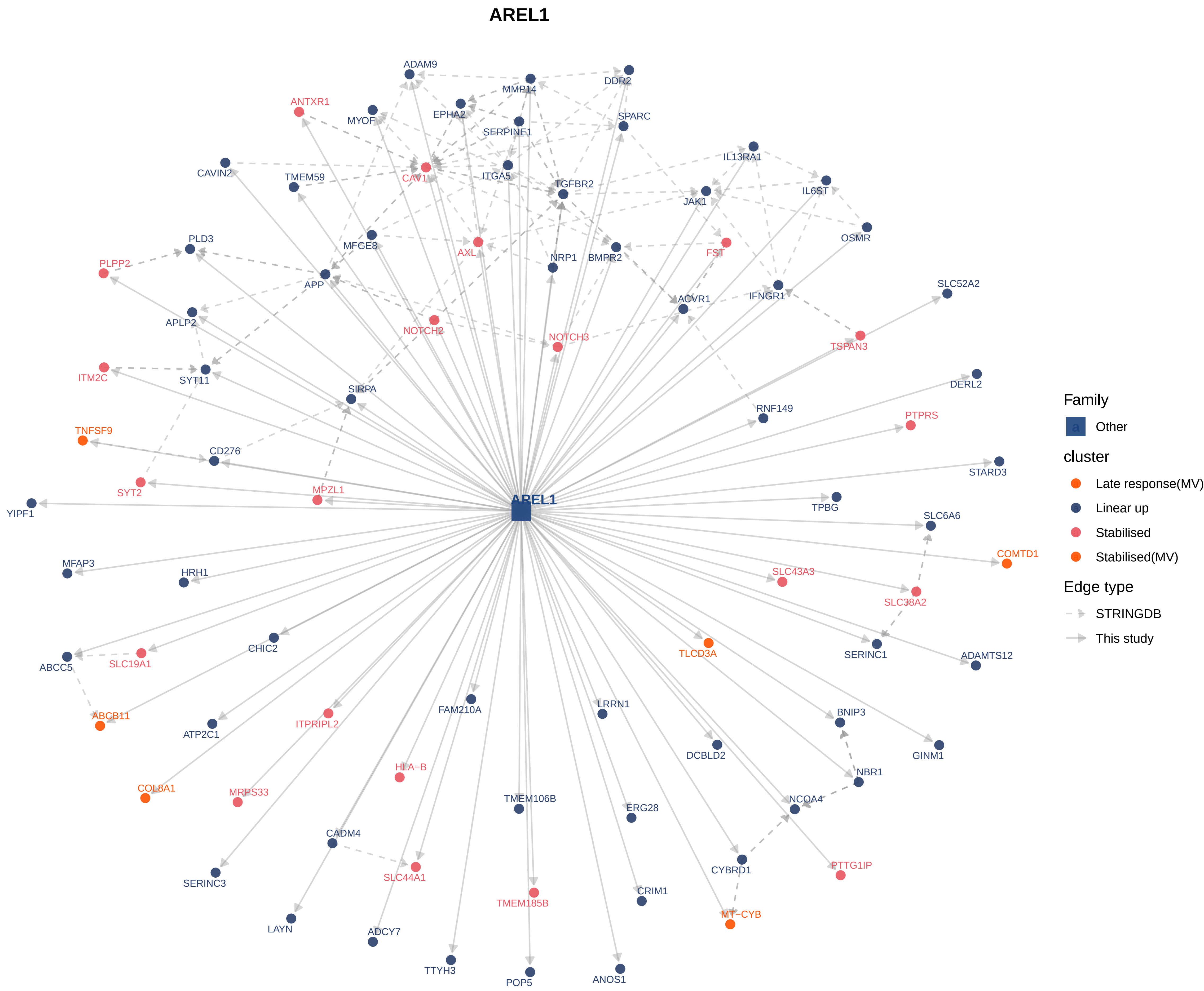

AREL1
ADAM9
MMP14
DDR2
ANTXR1
MYOF
EPHA2
SERPINE1
SPARC
CAVIN2
TMEM59
CAV1
ITGA5
TGFBR2
IL13RA1
JAK1
IL6ST
OSMR
PLD3
PLPP2
MFGE8
AXL
NRP1
BMPR2
FST
APP
ACVR1
IFNGR1
SLC52A2
APLP2
NOTCH2
NOTCH3
TSPAN3
ITM2C
SYT11
SIRPA
DERL2
RNF149
PTPRS
TNFSF9
CD276
STARD3
SYT2
MPZL1
YIPF1
TPBG
SLC6A6
COMTD1
MFAP3
HRH1
SLC43A3
SLC38A2
CHIC2
ABCC5
SLC19A1
TLCD3A
SERINC1
ADAMTS12
ABCB11
ATP2C1
ITPRIPL2
FAM210A
LRRN1
BNIP3
DCBLD2
GINM1
HLA−B
NBR1
COL8A1
MRPS33
TMEM106B
ERG28
NCOA4
CADM4
SLC44A1
SERINC3
TMEM185B
CRIM1
CYBRD1
PTTG1IP
MT−CYB
LAYN
ADCY7
TTYH3
POP5
ANOS1
Family
Other
cluster
Late response(MV)
Linear up
Stabilised
Stabilised(MV)
Edge type
STRINGDB
This study

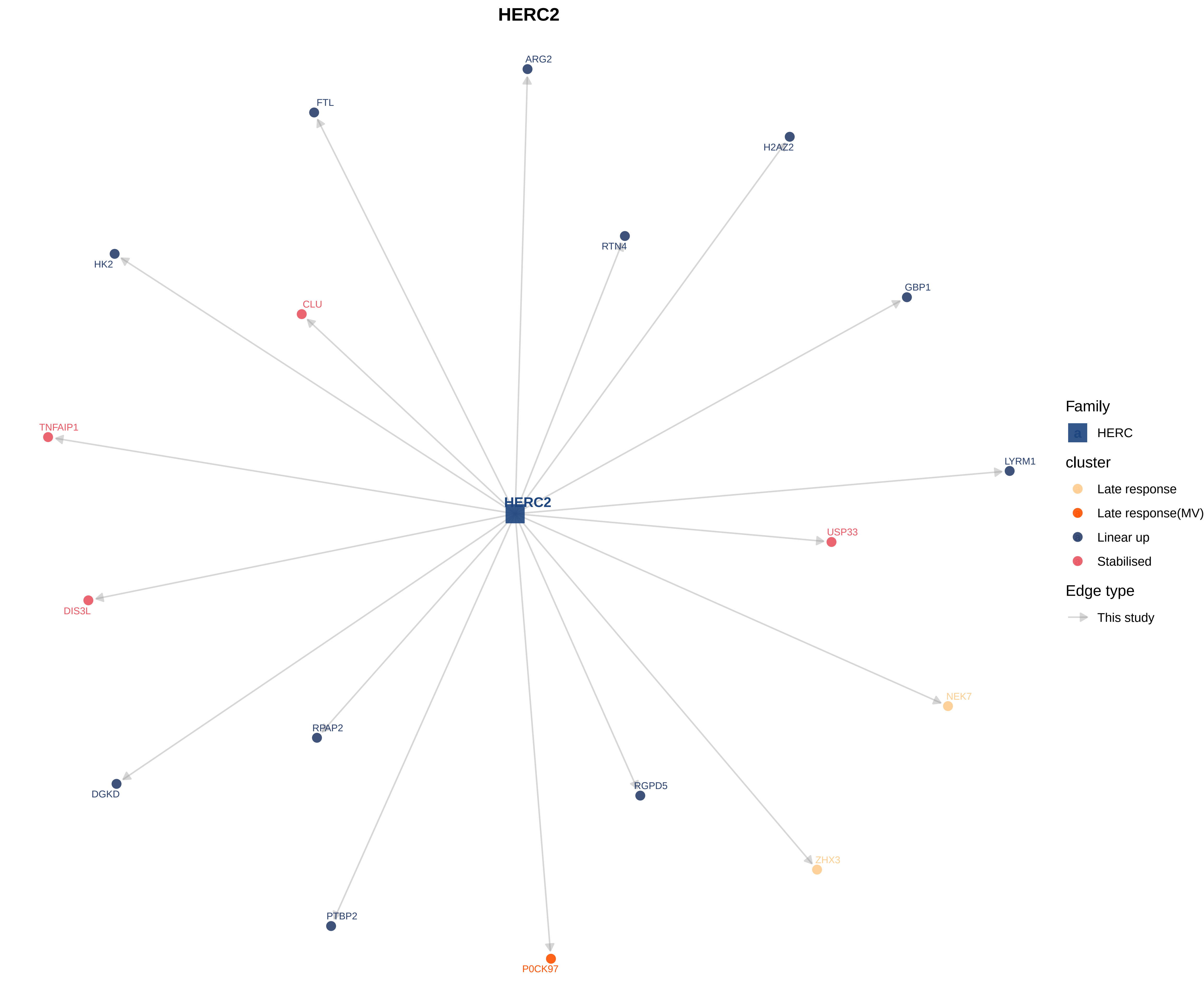

HERC2
ARG2
FTL
H2AZ2
RTN4
HK2
GBP1
CLU
TNFAIP1
LYRM1
HERC2
USP33
DIS3L
NEK7
RPAP2
DGKD
RGPD5
ZHX3
PTBP2
P0CK97
Family
HERC
cluster
Late response
Late response(MV)
Linear up
Stabilised
Edge type
This study

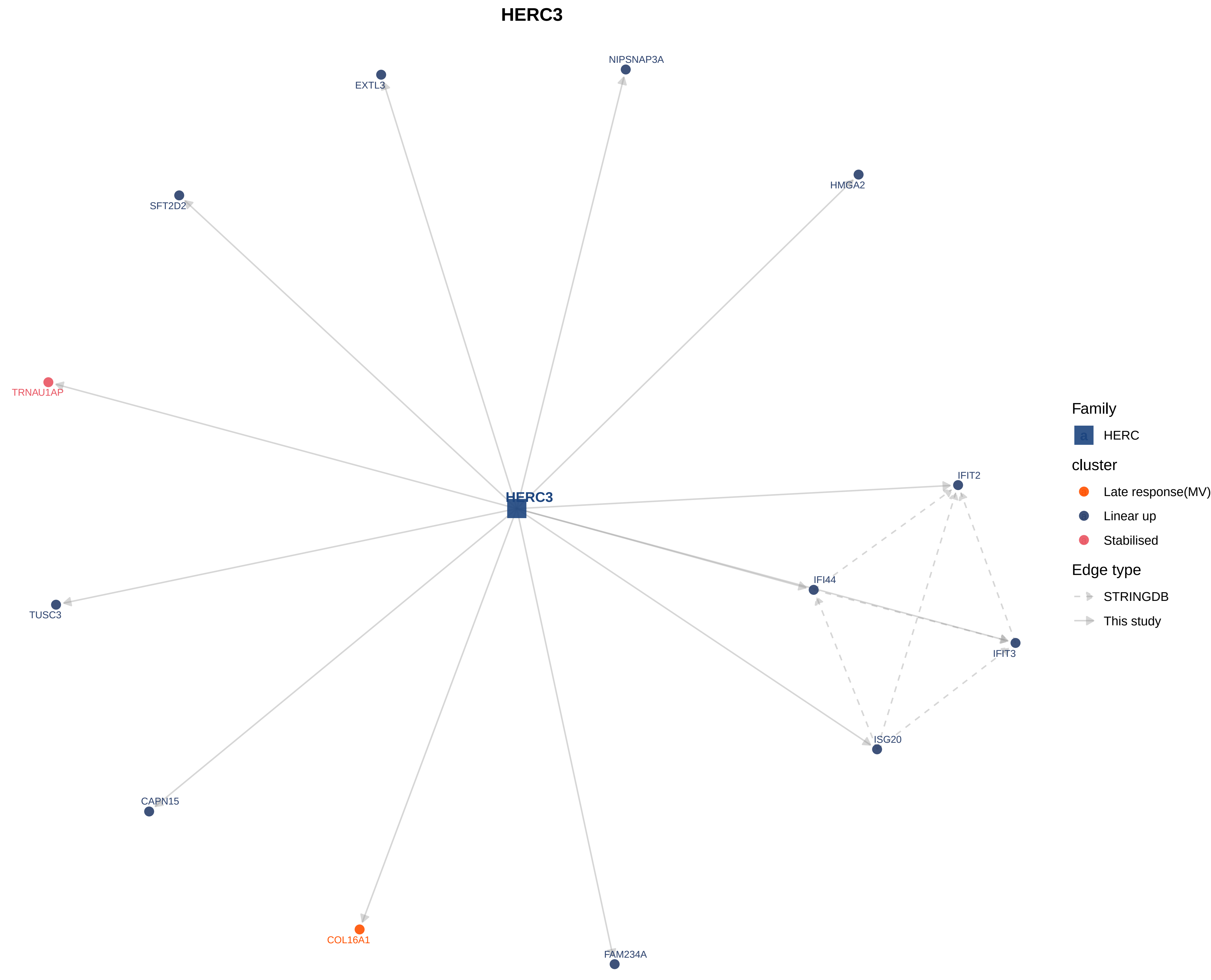
HERC3
NIPSNAP3A
EXTL3
HMGA2
SFT2D2
TRNAU1AP
HERC3
IFIT2
IFI44
IFIT3
TUSC3
ISG20
CAPN15
COL16A1
FAM234A
Family
HERC
cluster
Late response(MV)
Linear up
Stabilised
Edge type
STRINGDB
This study

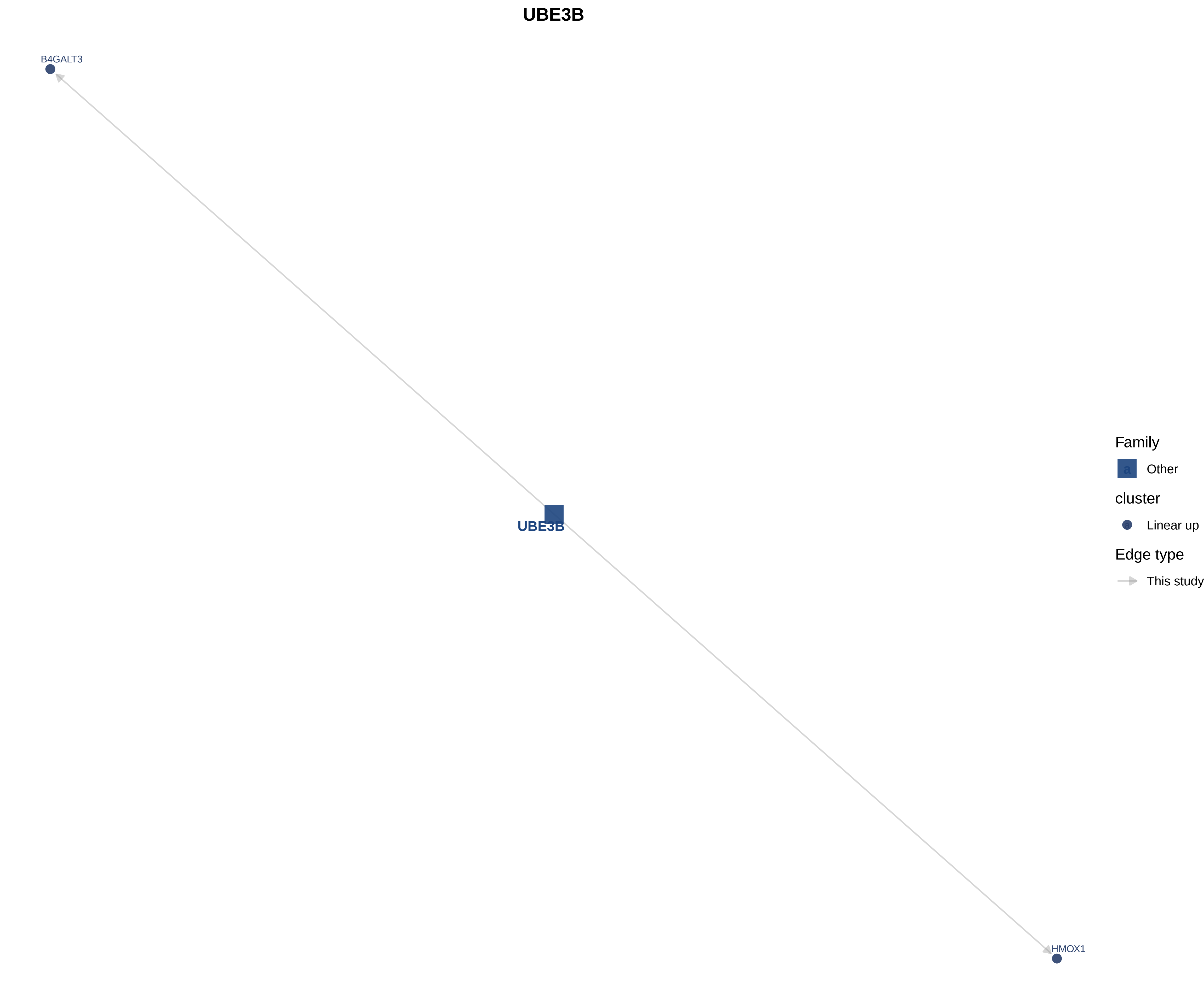
UBE3B
B4GALT3
UBE3B
HMOX1
Family
Other
cluster
Linear up
Edge type
This study

### HECTD4

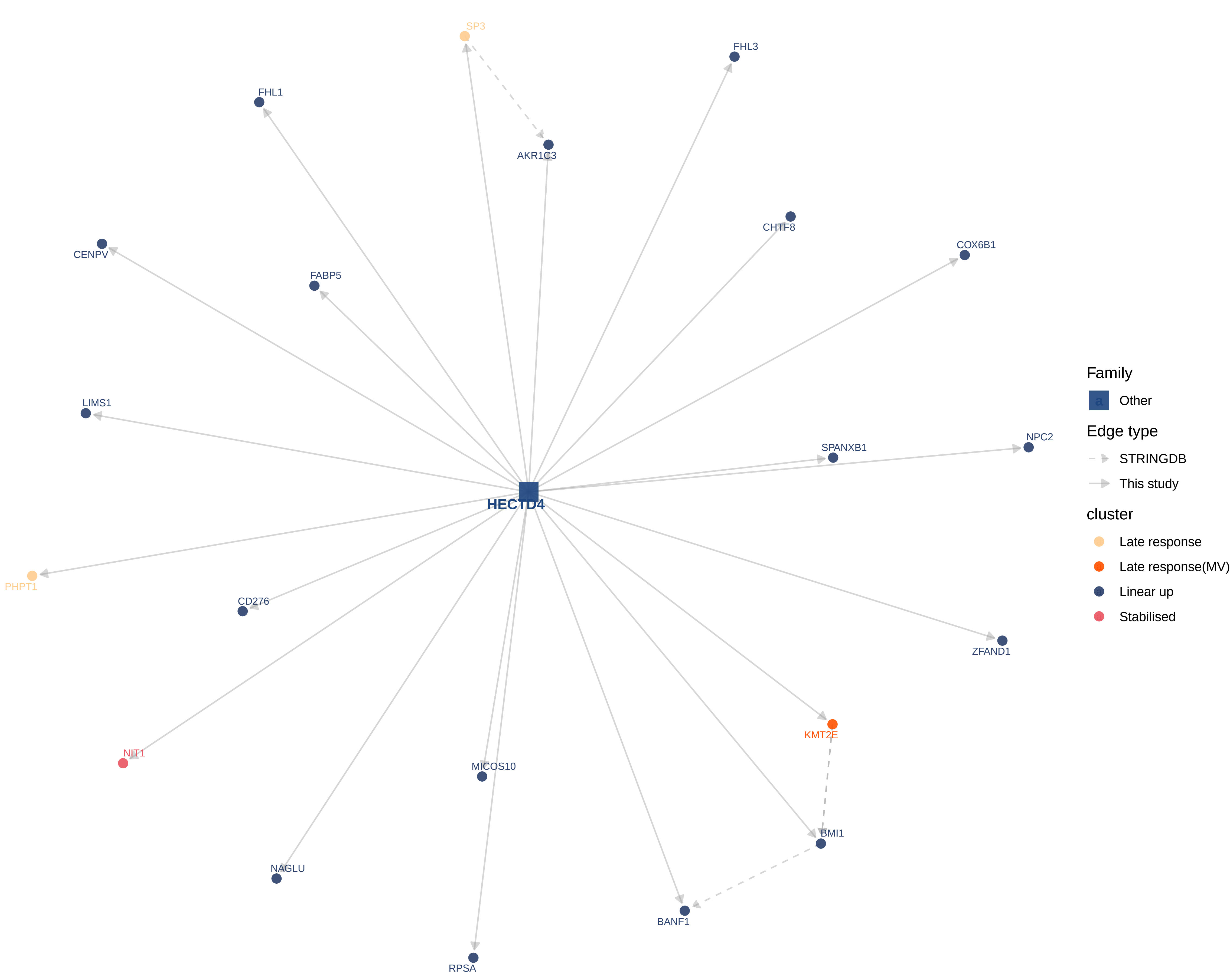

SP3
FHL3
FHL1
AKR1C3
CHTF8
CENPV
COX6B1
FABP5
LIMS1
SPANXB1
NPC2
HECTD4
PHPT1
CD276
ZFAND1
KMT2E
NIT1
MICOS10
BMI1
NAGLU
BANF1
RPSA
Family
Other
Edge type
STRINGDB
This study
cluster
Late response
Late response(MV)
Linear up
Stabilised

### HECTD1

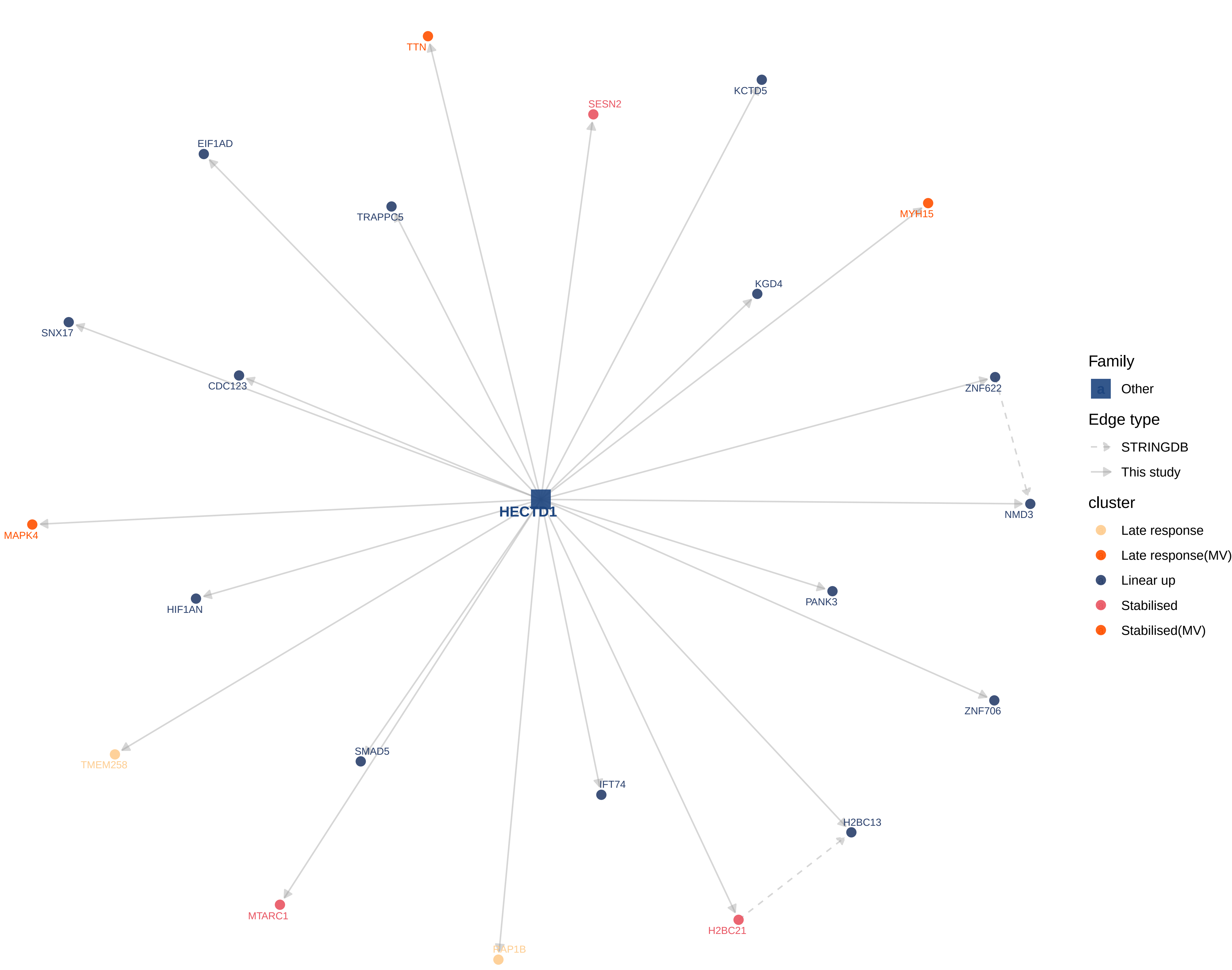
TTN
KCTD5
SESN2
EIF1AD
TRAPPC5
MYH15
KGD4
SNX17
CDC123
ZNF622
HECTD1
NMD3
MAPK4
HIF1AN
PANK3
ZNF706
TMEM258
SMAD5
IFT74
H2BC13
MTARC1
H2BC21
RAP1B
Family
Other
Edge type
STRINGDB
This study
cluster
Late response
Late response(MV)
Linear up
Stabilised
Stabilised(MV)

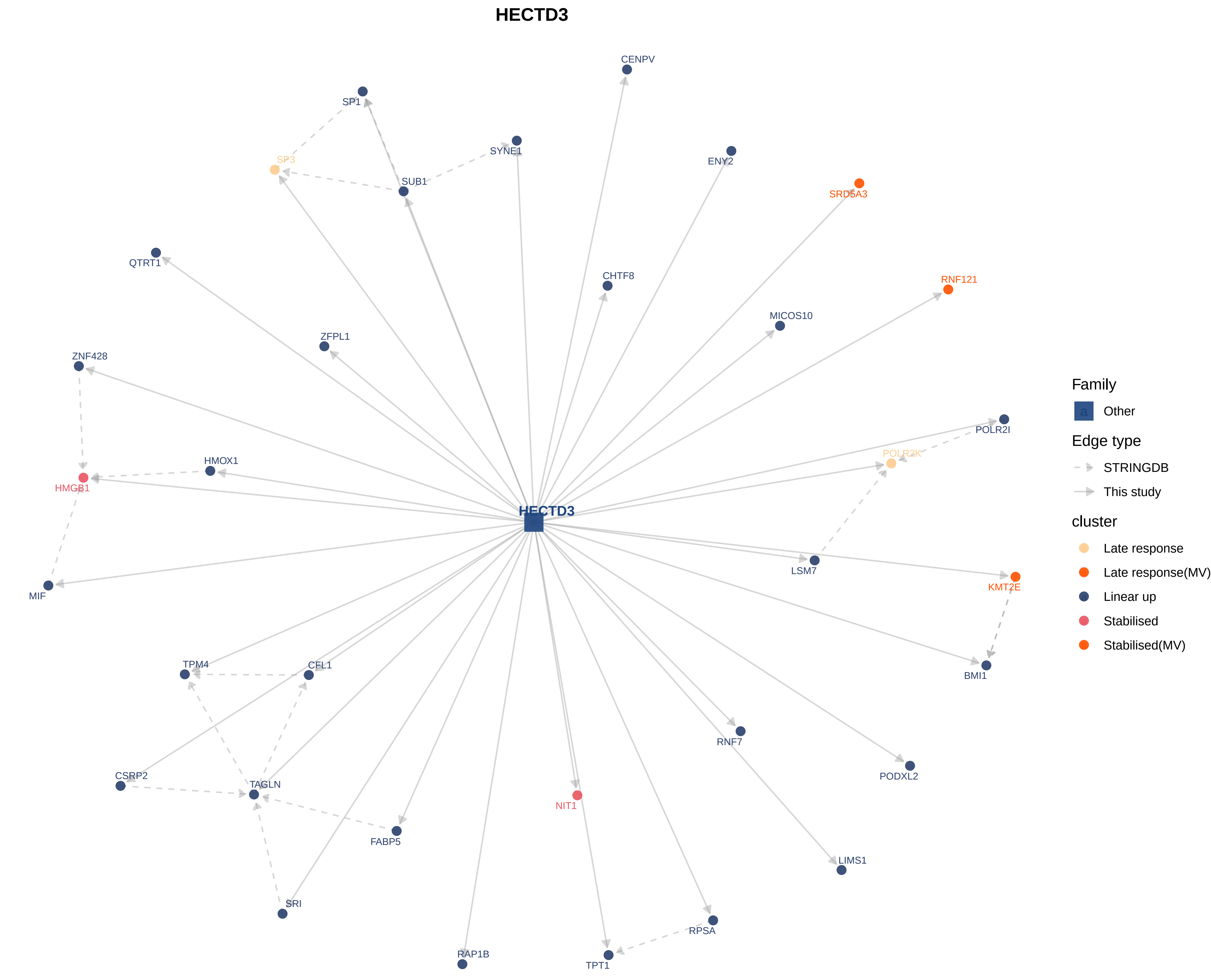
HECTD3
CENPV
SP1
SYNE1
ENY2
SP3
SUB1
SRD5A3
QTRT1
CHTF8
RNF121
MICOS10
ZFPL1
ZNF428
POLR2I
POLR2K
HMOX1
HMGB1
HECTD3
LSM7
KMT2E
MIF
BMI1
TPM4
CFL1
RNF7
PODXL2
CSRP2
TAGLN
NIT1
FABP5
LIMS1
SRI
RPSA
RAP1B
TPT1
Family
Other
Edge type
STRINGDB
This study
cluster
Late response
Late response(MV)
Linear up
Stabilised
Stabilised(MV)

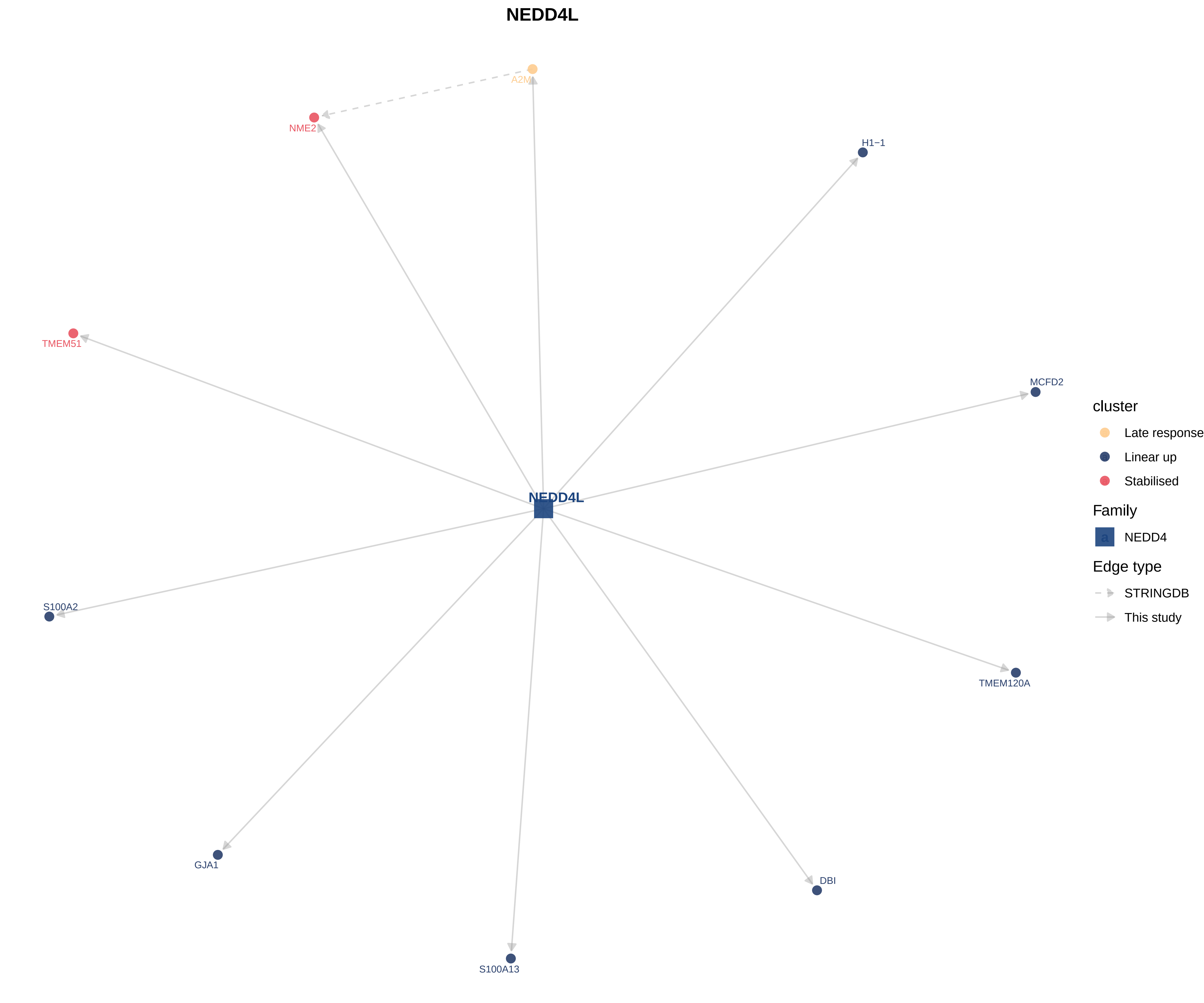
NEDD4L
A2M
NME2
H1−1
TMEM51
MCFD2
NEDD4L
S100A2
TMEM120A
GJA1
DBI
S100A13
cluster
Late response
Linear up
Stabilised
Family
NEDD4
Edge type
STRINGDB
This study

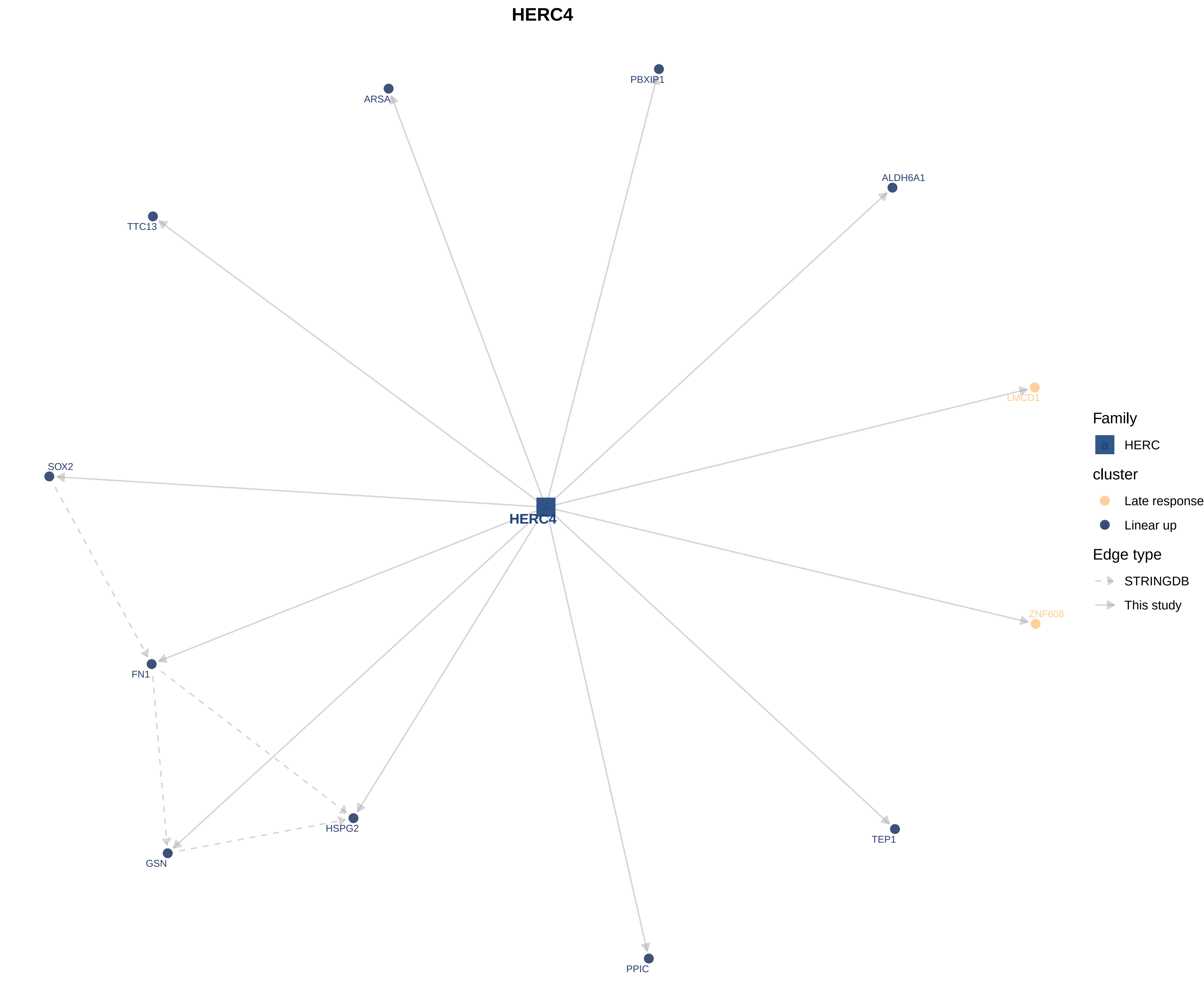
HERC4
PBXIP1
ARSA
ALDH6A1
TTC13
LMCD1
SOX2
HERC4
ZNF608
FN1
HSPG2
GSN
TEP1
PPIC
Family
HERC
cluster
Late response
Linear up
Edge type
STRINGDB
This study

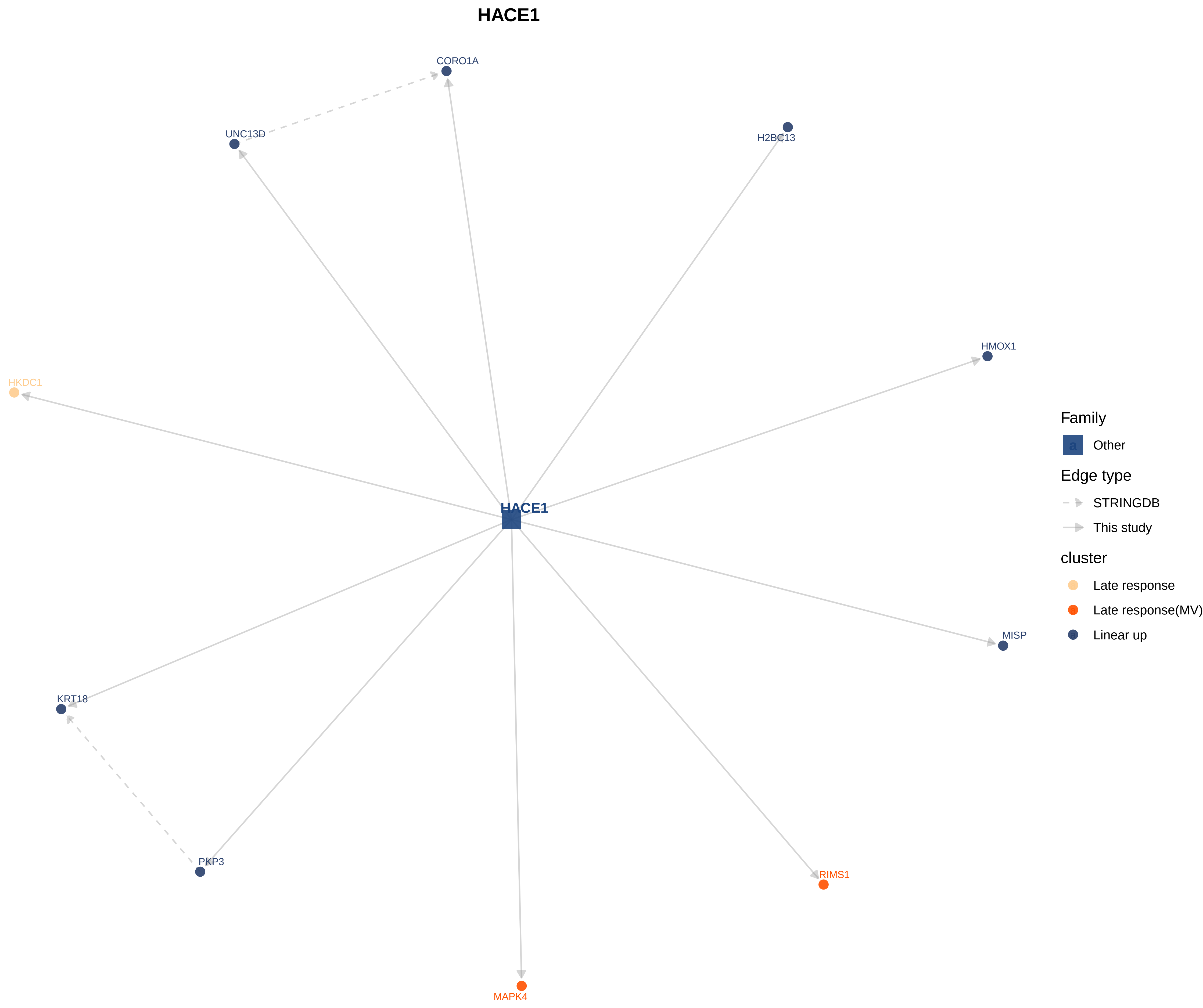
HACE1
CORO1A
UNC13D
H2BC13
HMOX1
HKDC1
HACE1
MISP
KRT18
PKP3
RIMS1
MAPK4
Family
Other
Edge type
STRINGDB
This study
cluster
Late response
Late response(MV)
Linear up

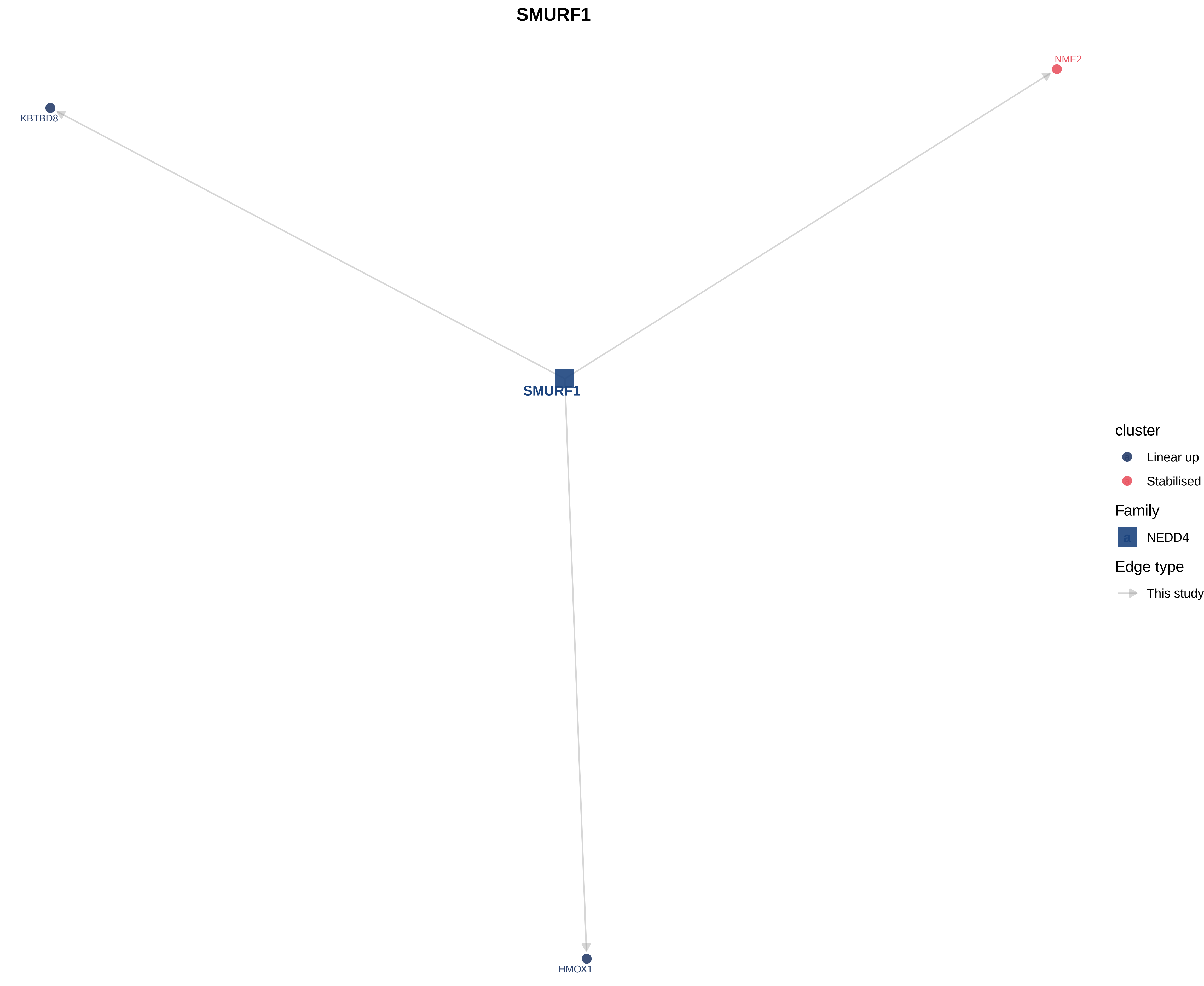
SMURF1
NME2
KBTBD8
SMURF1
HMOX1
cluster
Linear up
Stabilised
Family
NEDD4
Edge type
This study

### ITCH

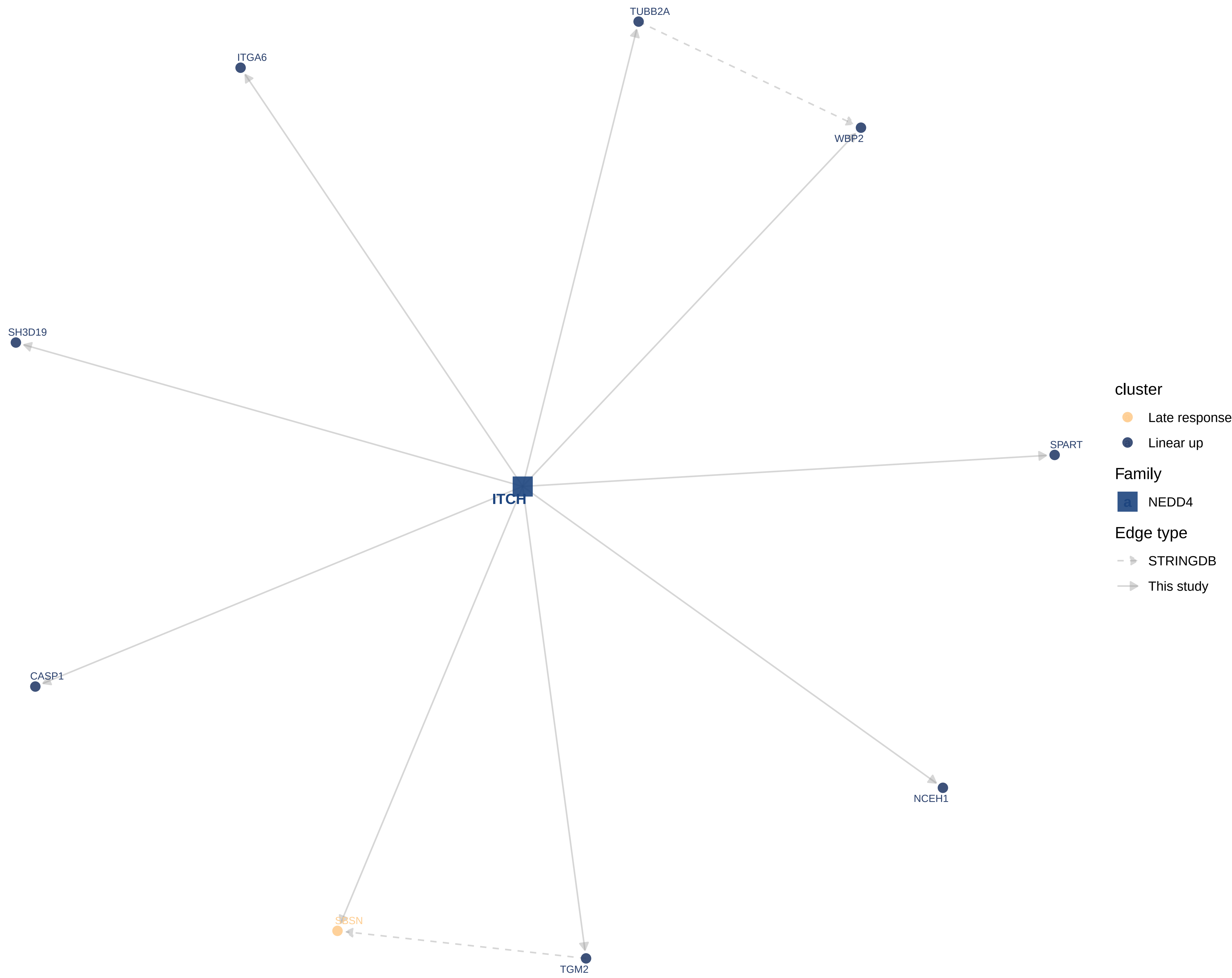

TUBB2A
ITGA6
WBP2
SH3D19
SPART
ITCH
CASP1
NCEH1
SBSN
TGM2
cluster
Late response
Linear up
Family
NEDD4
Edge type
STRINGDB
This study

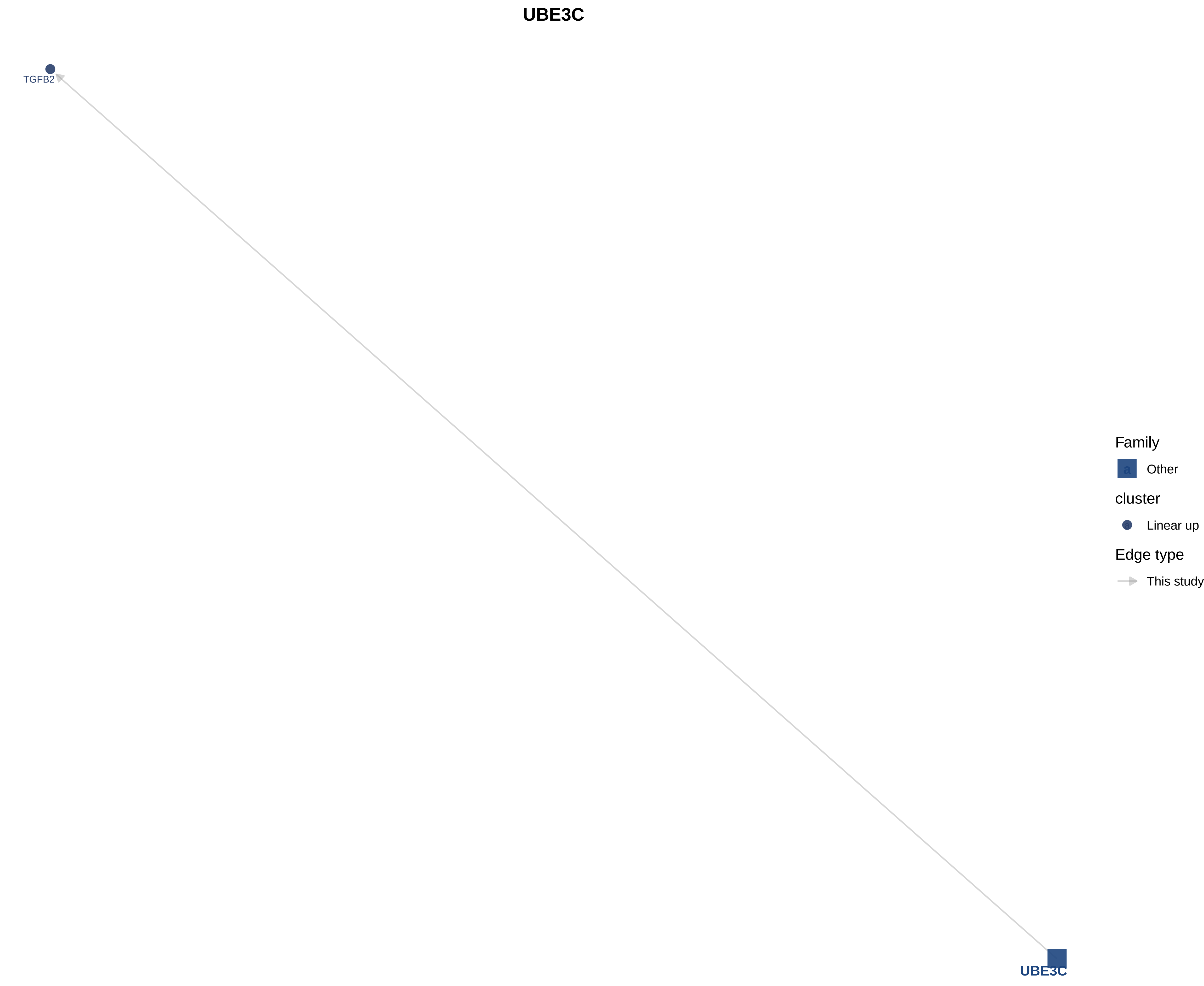

UBE3C
TGFB2
UBE3C
Family
Other
cluster
Linear up
Edge type
This study

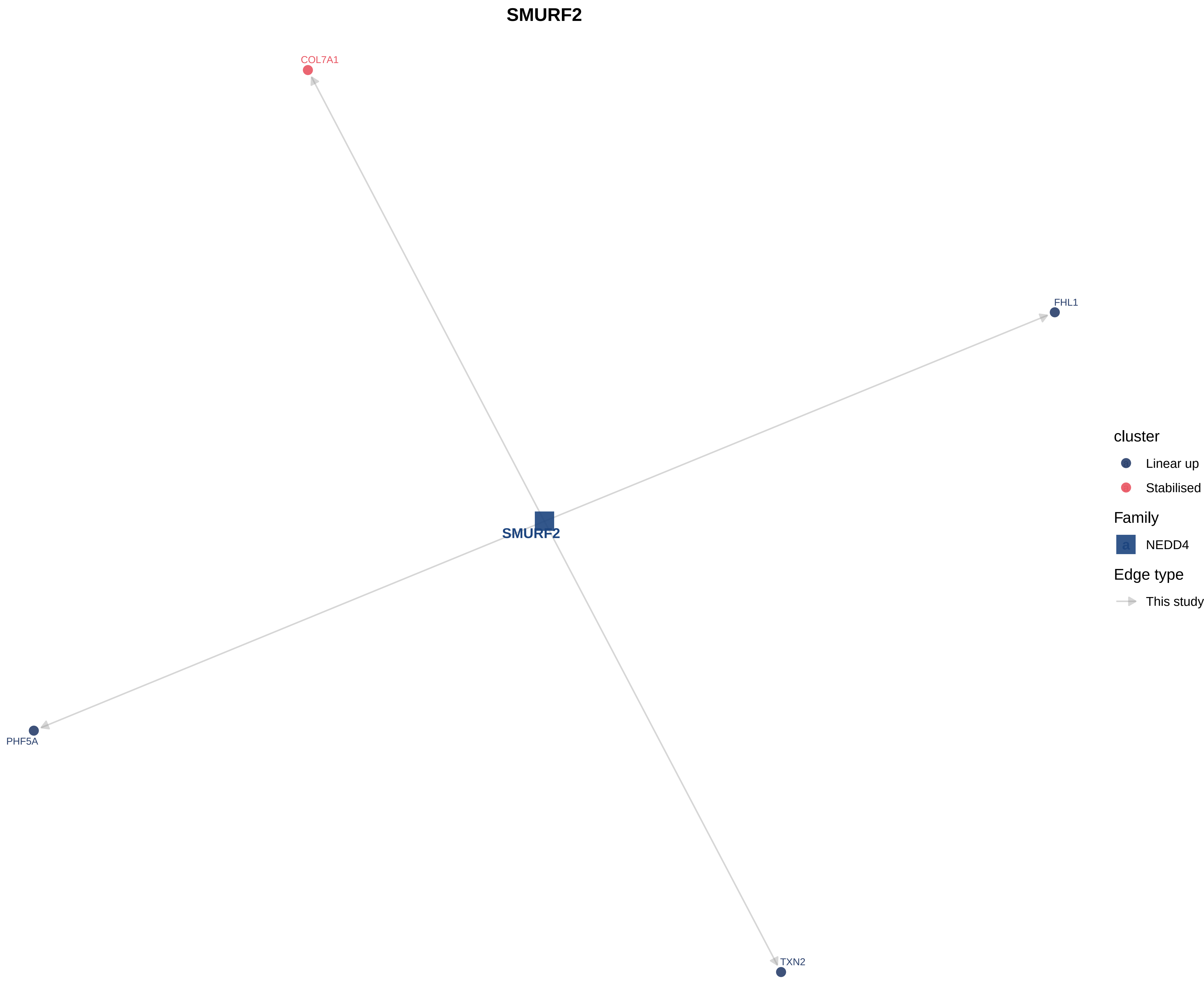

SMURF2
COL7A1
FHL1
SMURF2
PHF5A
TXN2
cluster
Linear up
Stabilised
Family
NEDD4
Edge type
This study

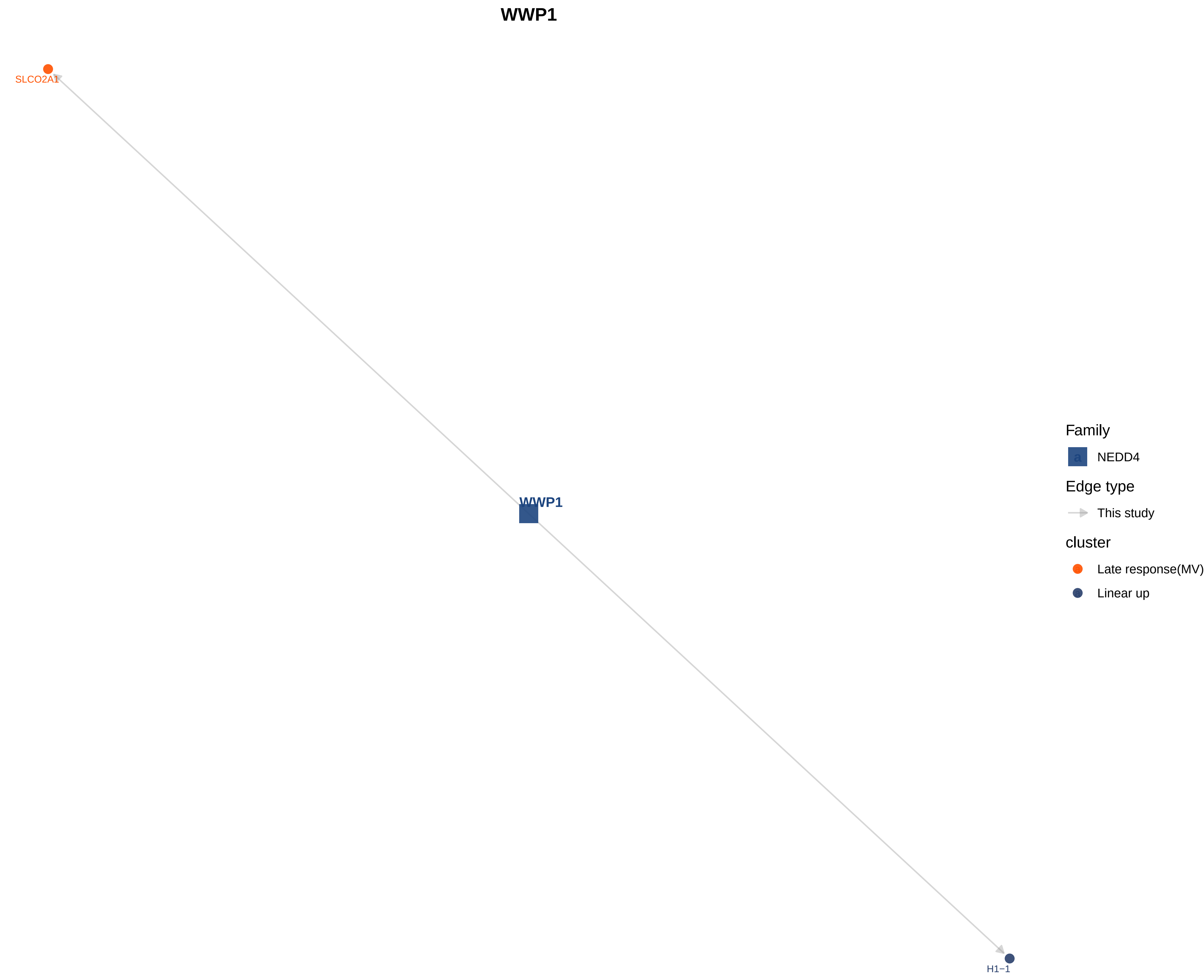

WWP1
SLCO2A1
WWP1
H1−1
Family
NEDD4
Edge type
This study
cluster
Late response(MV)
Linear up

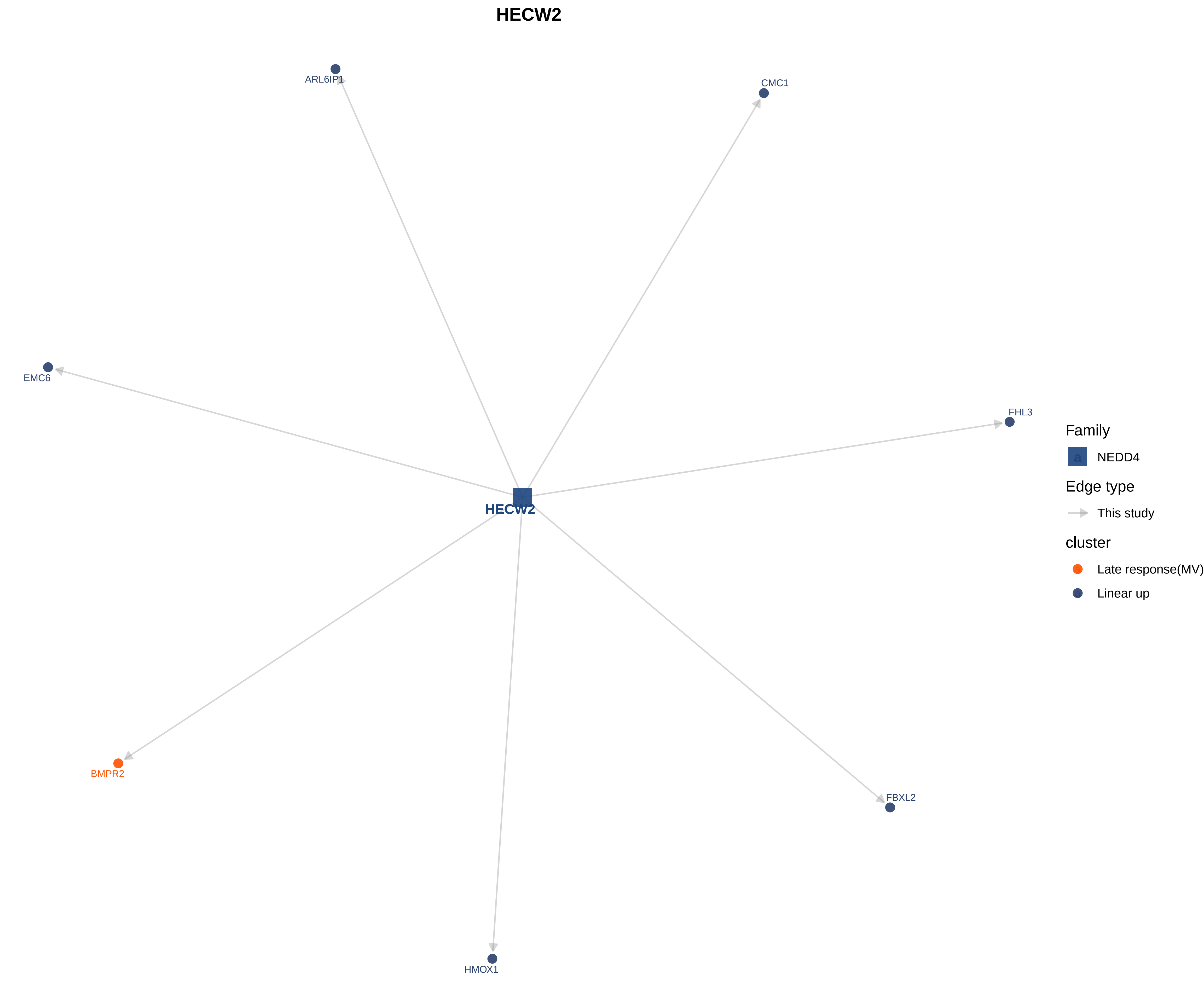
HECW2
ARL6IP1
CMC1
EMC6
FHL3
HECW2
BMPR2
FBXL2
HMOX1
Family
NEDD4
Edge type
This study
cluster
Late response(MV)
Linear up

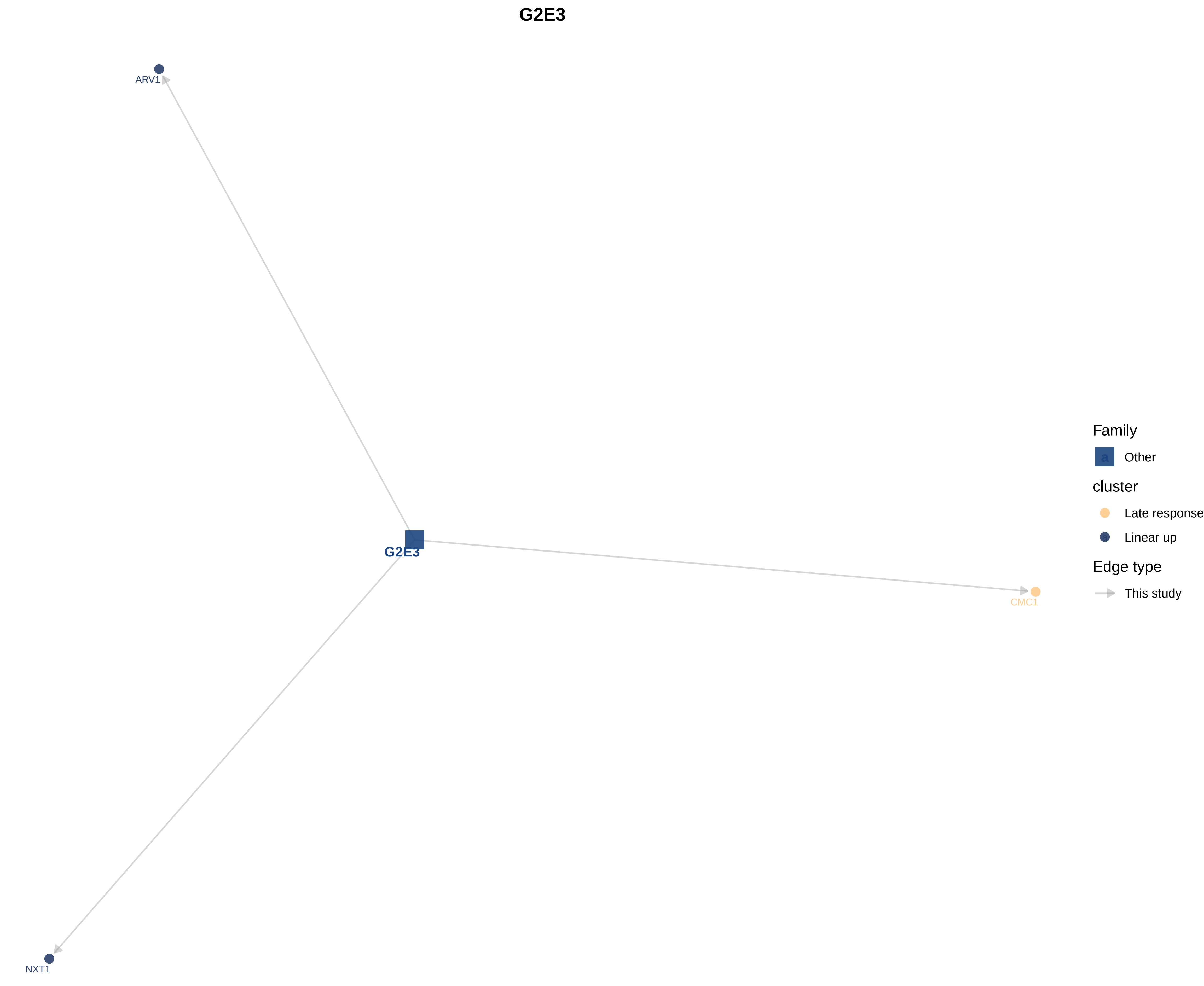
G2E3
ARV1
G2E3
CMC1
NXT1
Family
Other
cluster
Late response
Linear up
Edge type
This study

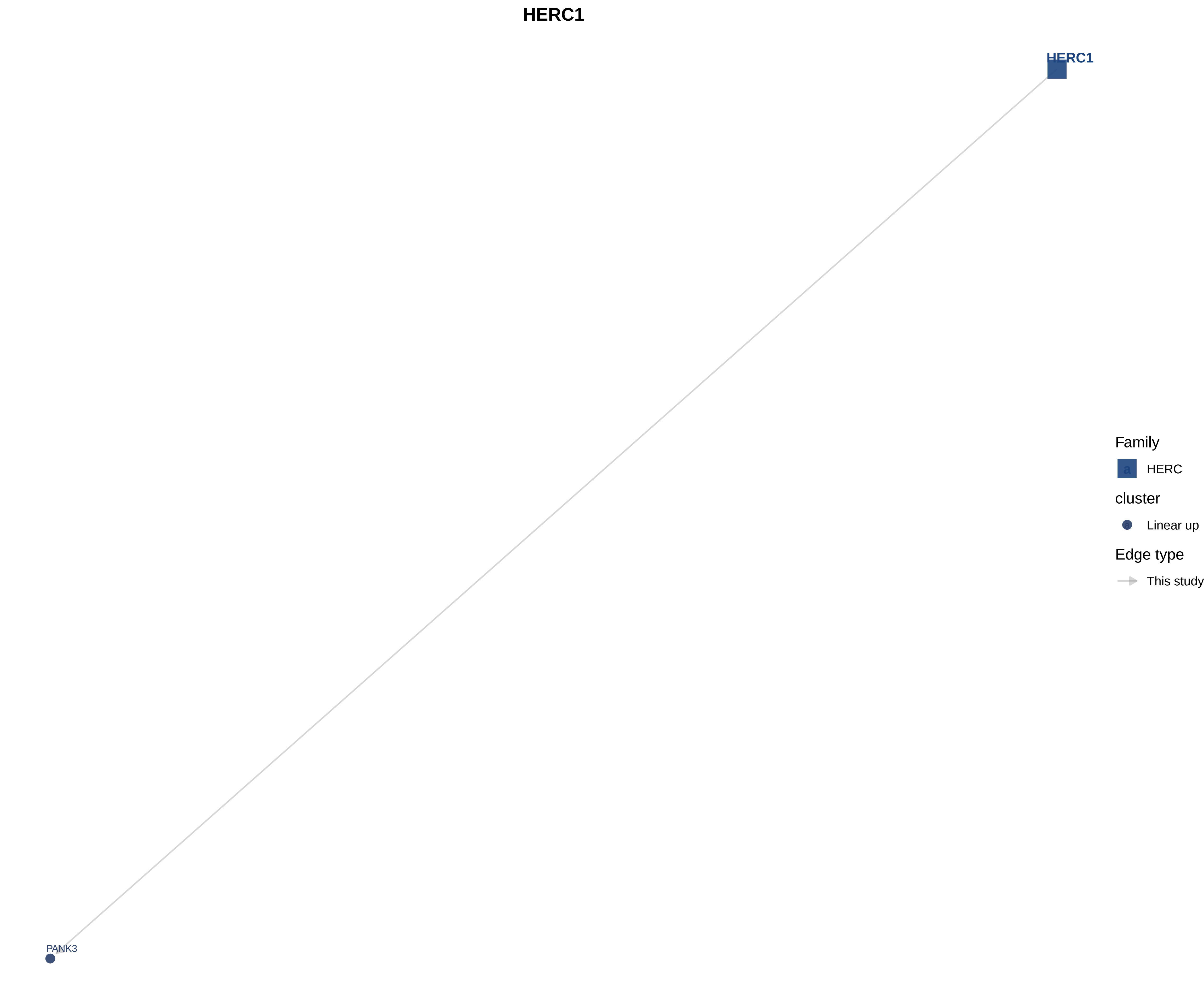
HERC1
HERC1
PANK3
Family
HERC
cluster
Linear up
Edge type
This study
