## Supplemental Figures and Captions for "A dynamic proteomic HECT E3 ligase screen reveals substrate exclusivity and redundancies"

Table of contents:

Table.S1 – TMTpro plex design for HECT E3 knockdown screen with plex-wise control

Table.S2 – Summarized data from dynamic HECT E3 knockdown (KD) screen. Protein and their corresponding gene names for each siRNA along with their log2FC and q value (vs inter-plex control) are calculated using PBLMM for both light and heavy SILAC channel (_L and _H respectively). Additional information such as gene name (Gene), HECT family annotation, psm count (psm_),‘confidence’ based on psm count, median log2FC (med_), difference in log2fc between light and heavy (diff), number of siRNAs with detection (n_), and outlier scores from statistical and ML based algorithms are included.

Table.S3 – Subset of Table.S2 with annotated interactors of individual HECT E3s from Biogrid, DeepUSI and Ubibrowser.

Table.S4 and Table.S5 – Light and Heavy Protein log2FC data from pSILAC-DIA experiment of combinatorial KD in section 4, respectively. For each siRNA, the log2FC (Median) are calculate against non-targeting control without imputation using eBayes modelling and corresponding q value (pLTcomp) are included.

Table.S6 – PRM based log2FC and q value data from 69 selected proteins for targeted assay on combinatorial KD exported from Skyline. The foldchanges are calculated against same control as before using Anova and Tukey median polish within Skyline.

Table.S7 – RNAseq fold change data

Table.S8 and Table.S9 - Light and Heavy Protein log2FC data from pSILAC-DIA experiment coIP experiment against MCL1 and IgG (IP control), for HUWE1 siRNA against non-targeting control. Inter IP (MCL1 vs IgG) and inter condition (HUWE1 vs NTC) log2FC (Median_) and corresponding q value based on eBayes modelling are reported.

Supplementary Figures:

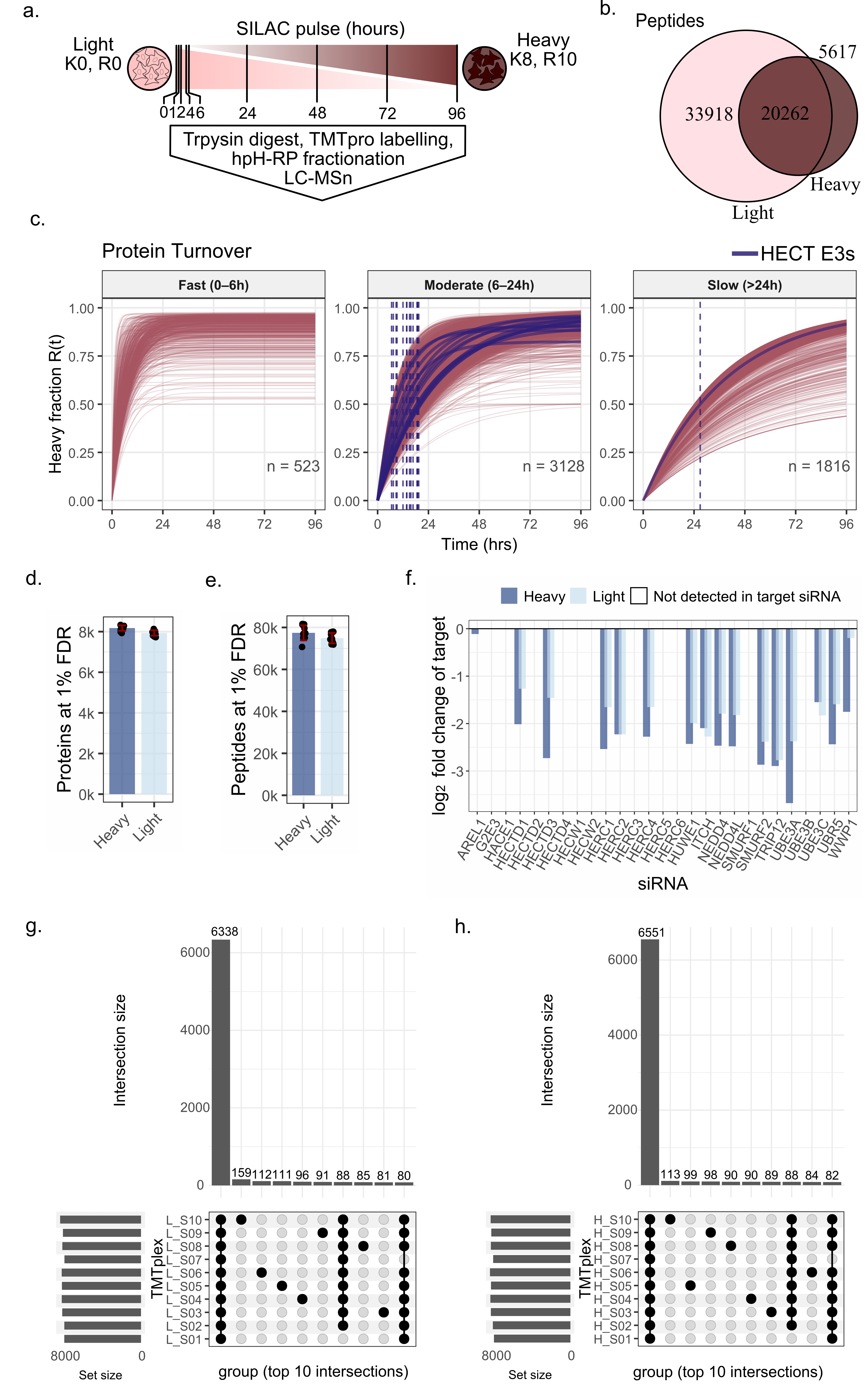

Fig.S1: HECT E3 essentiality is coupled with proteome turnover and cell proliferation. a. Timepoints and workflow for hyperplexed turnover proteomics. Cells were supplemented with heavy SILAC media for 8 time points from 1 h to 96 h (2 biological replicates), followed by trypsin digestion, TMTpro labelling, high pH fractionation and LC-MS with real-time search enabled SPS-MS3. b. Number of peptides identified as SILAC light and heavy in turnover experiment. c. Protein level turnover curves separated by apparent half-lives between 0-6 h (fast), 6-24 h (moderate) and >24 h (slow). 16 identified HECT E3s curves are marked in purple with apparent half-life as vertical line, showing moderate to slow turnover. d. Quantified proteins and e. peptides at 1% FDR across the HECT E3 KD screen. f. log_2_FC of target HECT E3s, showing successful knockdown in light and heavy proteome for 16 quantified HECT E3s. g. Total overlap of quantified proteins in light and h. heavy SILAC channel across all 10 TMT-pro plexes from HECT E3 screen.

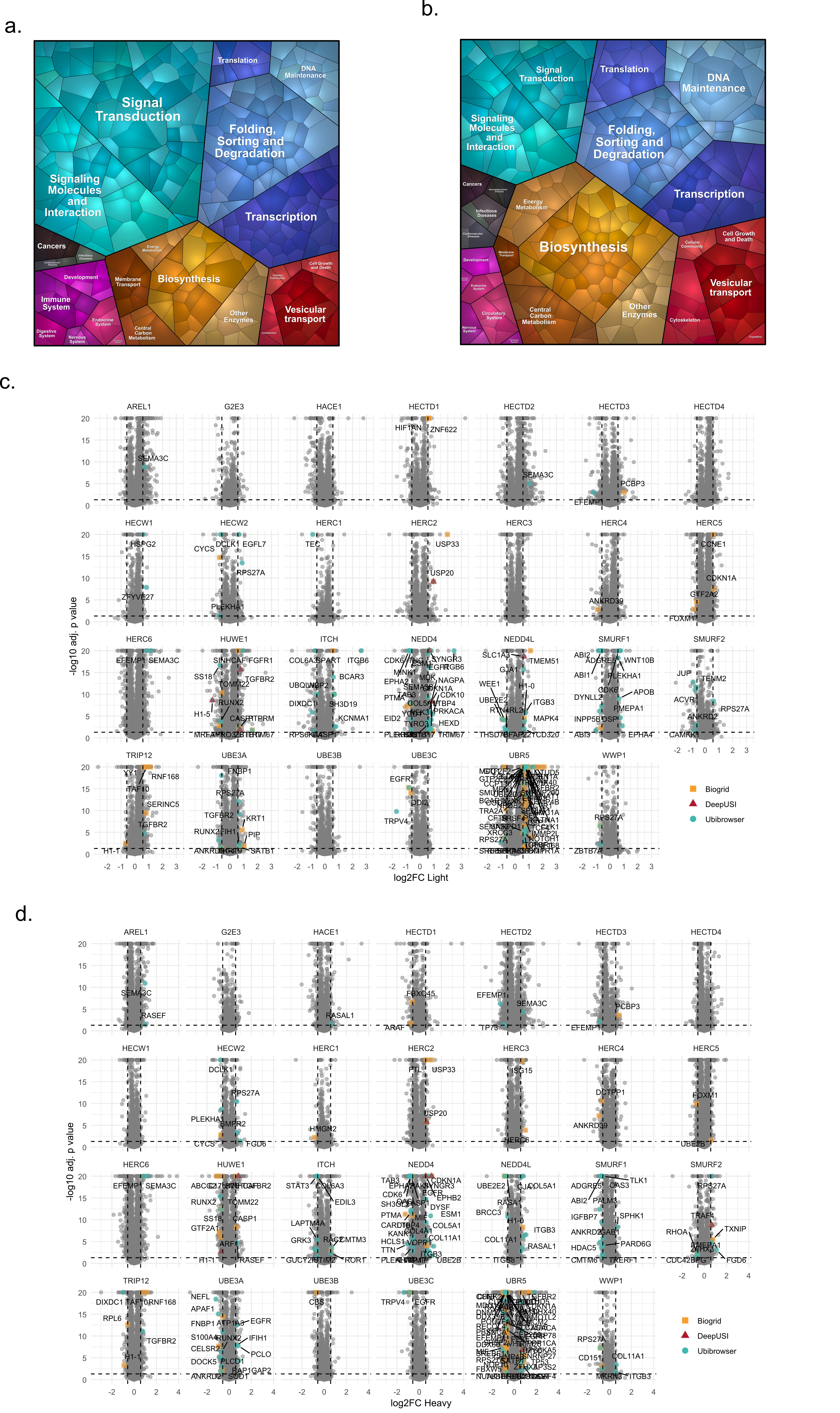

Fig.S2: Many UPS related processes are co-regulated by HECT E3s. a. & b. Voronoi plot showing major KEGG pathways represented by differentially expressed up and down-regulated proteins respectively. Only protein differentially expressed in the same trajectory in two or more knockdowns were filtered and run on proteomaps.net. Each polygon represents a protein and size represents occurrences. Proteins belonging to similar KEGG pathways are grouped under larger polygon. c & d. Log_2_FC (cut-off line 0.58) and adjusted p value (maxxed to -log_10_(20)) of light and heavy proteins respectively. Genes annotated as known substrate or interactor of each HECT E3 on Biogrid, Ubibrowser and DeepUSI are marked by shapes and color and labelled.

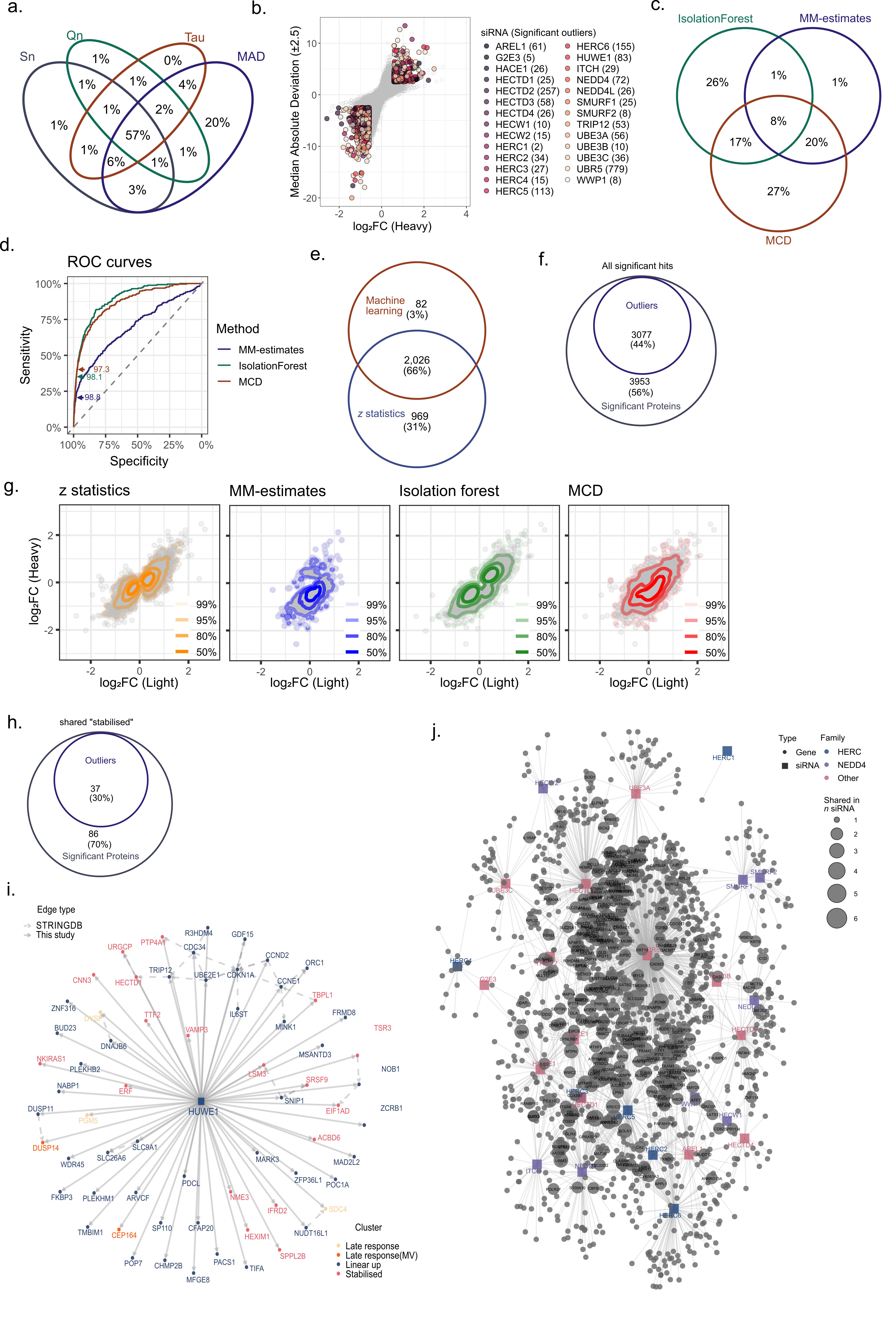
Fig.S3: Robust statistics and machine learning resolves exclusivity in HECT E3-substrate relationships. a. Overlap of all outliers above z threshold (±2.5) with all four z methods namely (Sn, Qn, Tau and MAD) b. Distribution of significant DEPs (log_2_FC ±0.58 q value < 0.05) for heavy proteome on x-axis versus corresponding MAD score on y-axis. Dots colored for individual siRNA and above log_2_FC, q value and z threshold. Legend indicates the number of outliers per knockdown. c. Overlap of all dynamic outliers above score threshold for three ML approaches (score cutoffs & specificity %: MM-estimate: 0.1 = 98.8%, Isolation forest: 0.7 = 98.1% and MCD: 23.0 = 97.3%) .d Receiver operating characteristics (ROC) curves for ML results against annotated, significantly expressed substrates and interactors as true positive i.e. sensitivity on y-axis and (1 - false discovery rate) of score distribution i.e. specificity on x-axis. e. Overlap of significant DEPs above outlier threshold for z statistics and machine learning approaches. f. subset of outliers out of significant DEPs across all HECT KDs. g. Distribution of all outliers classified by z statistics (MAD, Qn, Sn and tau combined), MM-estimates, isolation forest and MCD, with density probability (%) annotated in each panel. h. subset of shared outliers out of all shared significant DEPs across all HECT KDs. i. Example of exclusive targets of HECT E3s and string network incorporation. Outliers belonging to HUWE1 KD are mapped (solid) along with annotated STRING (scoring threshold 400 v.12.0) interaction amongst them. j. Network plot of downregulated outliers (dot) (n =1,779) per HECT E3 siRNA (square, colored by subfamily). The size of the dot indicates whether the protein is a shared outlier between HECT E3s or unique. Shared outliers labeled by their respective gene name.

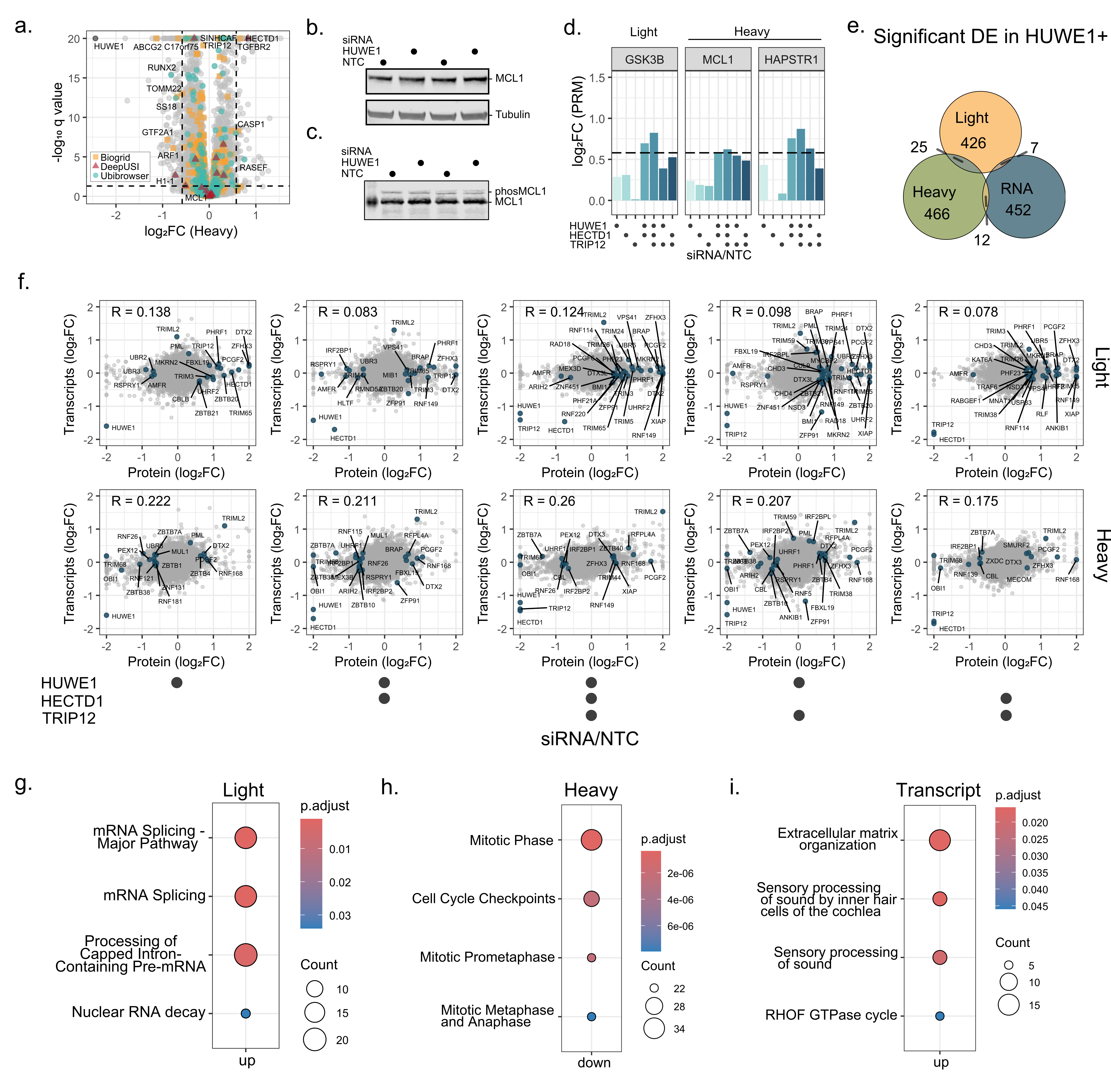

Fig.S4: Combinatorial knockdown reveals crosstalk along the HUWE1-TRIP12-HECTD1 axis. a. Volcano plot showing log_2_FC and -log_10_ q value of heavy proteins in HUWE1 KD, including annotated targets and substrates from Biogrid, Ubibrowser and DeepUSI. MCL1 remains unchanged and TRIP12 and HECTD1 increase. b. Western blot confirming stable MCL1 expression and c. phosphorylation under HUWE1 KD. d. Parallel reaction monitoring (PRM)-based confirmation of GSK3B expression in light and MCL1 and HAPSTR1 expression in heavy proteome across combination of HUWE1, HECTD1 and TRIP12 KDs. e. Overlap of significant DE proteins and transcripts in HUWE1+ combinatorial KD. f. Dotplot with transcript log_2_FC from RNAseq on y-axis and protein log_2_FC (light and heavy) on x-axis for combinations of KDs, with significantly expressed E3s in dark green and spearman R annotated at the top of each panel. g-i. Reactome pathways enriched in synergistically (HUWE1+ KD) upregulated proteins in light, downregulated proteins in heavy and upregulated transcripts respectively.

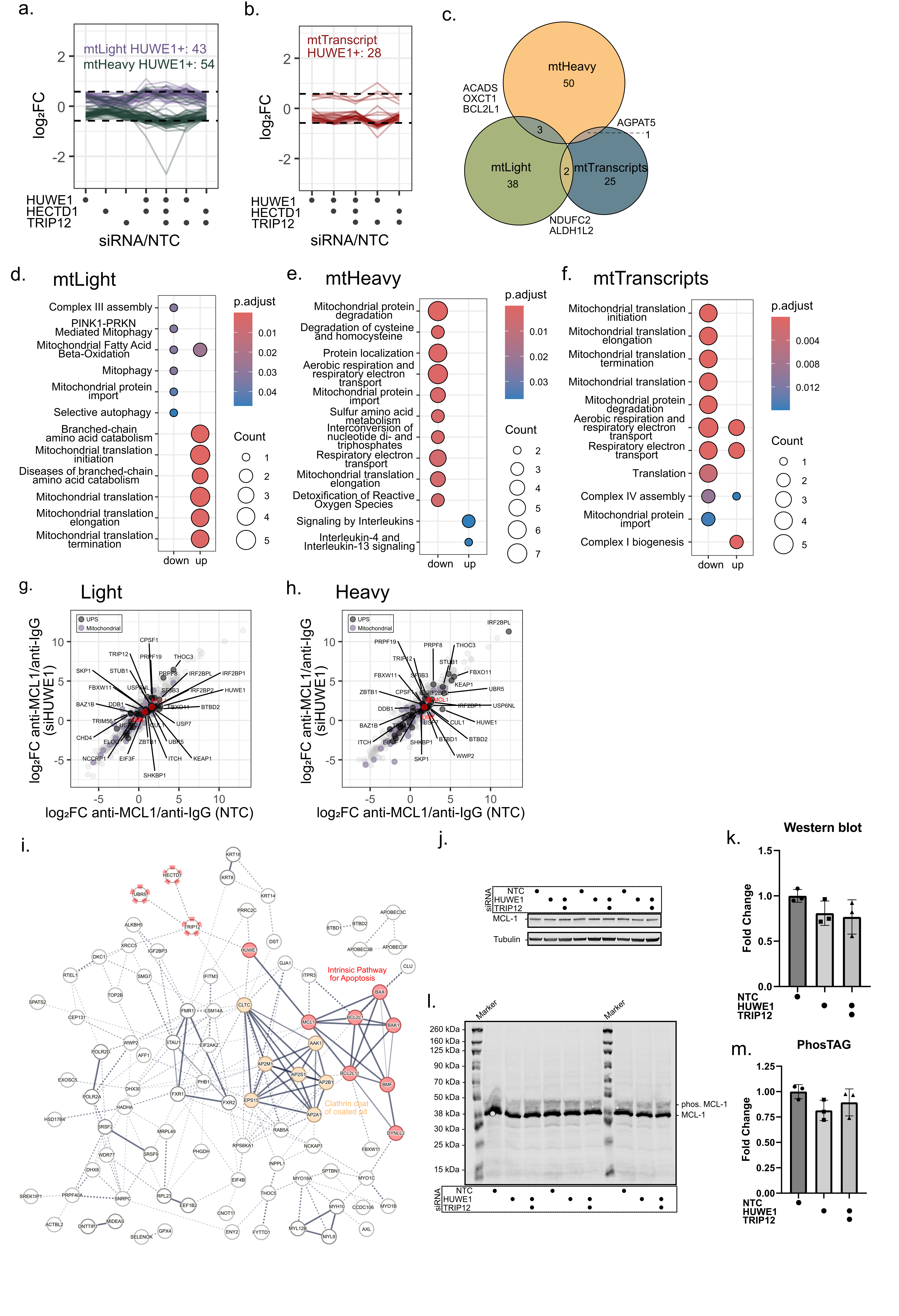

Fig.S5: Combinatorial knockdown reveals HUWE1-TRIP12-HECTD1 involvement in mitophagy, ROS clearance and apoptosis. a. Spaghetti plot of log_2_FC across KDs of mitochondrial (mt) subset of synergistic DEPs (HUWE1+) in light and heavy proteome and b. Transcriptome. c. Overlap between synergistic mt DEPs in light and heavy proteome and transcriptome. Shared ones are labelled with gene names. d-f. Biological processes (Gene ontology) represented in up and downregulated synergistic mt genes in light proteome, heavy proteome and transcriptome respectively. g-h. Correlation of log_2_FC (anti-MCL1/anti-IgG) of NTC (x-axis) vs HUWE1 KD (y-axis) in light and heavy SILAC channels for MCL1 and IgG co-IP. Mitochondrial proteins are colored purple and UPS related proteins are colored and labelled in black. i. STRING annotated interaction between all the DEPs in HUWE1 KD vs NTC anti-MCL1 co-IP, with UBR5, HECTD1 and MCL1 added to the list (from Fig.4g-h). Dotted/solid lines indicate annotated connections and thickness of the line indicates confidence. Solid lines indicate grouping by DBscan based on annotated pathway. j-k. SDS-PAGE and l-m. PhosTAG western blot for MCL1, showing consistent abundance of MCL1 and phosphorylated MCL1 in NTC control, HUWE1 KD and HUWE1+TRIP12 KD. k,m show mean values with standard deviation (n=3 biological replicates).
